# Analysis of structural brain asymmetries in Attention-Deficit/Hyperactivity Disorder in 39 datasets

**DOI:** 10.1101/2020.03.03.974758

**Authors:** Merel C. Postema, Martine Hoogman, ENIGMA ADHD Working Group, David C. Glahn, Neda Jahanshad, Sarah E. Medland, Paul M. Thompson, Simon E. Fisher, Barbara Franke, Clyde Francks

## Abstract

**Objective:** Some studies have suggested alterations of structural brain asymmetry in attention-deficit/hyperactivity disorder (ADHD), but findings have been contradictory and based on small samples. Here we performed the largest-ever analysis of brain left-right asymmetry in ADHD, using 39 datasets of the ENIGMA consortium.

**Methods:** We analyzed asymmetry of subcortical and cerebral cortical structures in up to 1,978 people with ADHD and unaffected 1,917 controls. Asymmetry Indexes (AIs) were calculated per participant for each bilaterally paired measure, and linear mixed effects modelling was applied separately in children, adolescents, adults, and the total sample, to test exhaustively for potential associations of ADHD with structural brain asymmetries.

**Results:** There was no evidence for altered caudate nucleus asymmetry in ADHD, in contrast to prior literature. In children, there was less rightward asymmetry of the total hemispheric surface area compared to controls (*t*=2.4, *P*=0.019). Lower rightward asymmetry of medial orbitofrontal cortex surface area in ADHD (*t*=2.4, *P*=0.007) was similar to a recent finding for autism spectrum disorder. There were also some differences in cortical thickness asymmetry across age groups. In adults with ADHD, globus pallidus asymmetry was altered compared to those without ADHD. However, all effects were small (Cohen’s *d* from −0.18 to 0.18) and would not survive study-wide correction for multiple testing.

**Conclusion:** Prior studies of altered structural brain asymmetry in ADHD were likely underpowered to detect the small effects reported here. Altered structural asymmetry is unlikely to provide a useful biomarker for ADHD, but may provide neurobiological insights into the trait.

## Introduction

Attention-deficit/hyperactivity disorder (ADHD) is among the most frequently diagnosed childhood-onset mental disorders, affecting 5% of individuals worldwide (1). ADHD is characterized by developmentally inappropriate and impairing levels of inattention and/or hyperactivity, impulsivity, and emotional dysregulation (2). At least 15% of children diagnosed with ADHD retain the diagnosis into adulthood (3, 4).

Left-right asymmetry (laterality) is an important feature of human brain organization (5-7), and altered structural or functional asymmetry has been reported for a range of psychiatric conditions (6). The right hemisphere is typically dominant for some aspects of attention and arousal (8), and it was observed in the 1980s that people with unilateral lesions in the right hemisphere can show ADHD-like symptoms (8). Since then, various neuropsychological and functional imaging studies have found lateralized differences between people with ADHD compared to controls (e.g., (9-14)), with some pointing to a particular involvement of right hemisphere alterations (15-20). However, not all functional data fit a primarily right-hemisphere model (21).

In terms of brain anatomy, several studies have reported altered asymmetry of the caudate nucleus in ADHD, although not consistently in the direction of effect (9, 10, 22-25). Altered asymmetry of grey matter volumes in the superior frontal and middle frontal gyri has been reported in ADHD (26), as well as decreased asymmetry of cortical convolution complexity in the prefrontal cortex (27). Douglas *et al.* (14) performed the largest study of brain anatomical asymmetry in ADHD to date, including 192 cases with ADHD with a history of pharmacotherapy, 149 medication-naïve cases with ADHD, and 508 typically developing controls (ages 6-21 years), from eight separate datasets. They calculated per-subject Asymmetry Indexes (AI) for various regional grey matter volumes, AI=(Left-Right)/((Left+Right)/2) (a widely used approach in studies of brain asymmetry (28-31)), but did not find any significant alterations of AIs in ADHD (14). However, in a subset of their dataset (56 cases and 48 controls), Douglas *et al.* (14) analyzed diffusion tensor imaging (DTI) data, including fractional anisotropy and mean diffusivity measures, and reported alterations in the asymmetry of six white matter tracts, again not specifically driven by alterations in the right hemisphere.

In the current study, we measured cortical regional AIs in 1,978 cases and 1,917 controls from 39 datasets, and subcortical AIs in 1,736 cases and 1,654 controls from 35 datasets, made available via the ADHD working group of the ENIGMA (Enhancing NeuroImaging Genetics through MetaAnalysis) consortium. The same datasets were recently analyzed in two other studies, by Hoogman *et al*.(32, 33), that investigated bilateral changes in subcortical volumes and cortical measures, but not alterations of asymmetry. They found that ADHD was associated with lower average volumes of various subcortical structures (33), as well as lower total and regional cortical surface areas (including frontal, cingulate and temporal regions), and decreased cortical thickness in fusiform gyrus and temporal pole (32). These effects were largest in children, and even child-specific for the cortical findings, so that for the present study of asymmetries, we followed the age-group division of Hoogman *et al.* (32) into children (<15 years), adolescents (15-21 years) and adults (>21 years), as well as performing analysis in the total combined sample to explore age-general effects. Bilateral effect sizes reported by Hoogman *et al.* (32, 33) were small, i.e. case-control Cohen’s *d* values between −0.21 and 0.06. This suggests that, if associations between ADHD diagnosis andregional brain asymmetries are similarly subtle, many previous studies of anatomical asymmetries in this disorder were underpowered, and the described effects may have been unreliable. The current study aims to address this by providing detailed information on how laterality is affected in ADHD, based on the largest ever sample size for this question, comprised of multiple independent cohorts from around the world.

## Methods and Materials

### Datasets

Bilateral brain measures derived from structural MRI were available from 39 different datasets via the ENIGMA-ADHD Working Group (**ST1**). The 39 datasets comprised cortical data from a total of 1,978 participants with ADHD (1,426 males; median age = 15 y; range = 4 y to 62 y) and 1,917 healthy individuals (1,146 males; median age = 14 y; range = 4 y to 63 y). Subcortical data were available from 35 of the 39 datasets and comprised 1,736 cases (1,246 males, median age = 15 y; range = 5 y to 62 y) and 1654 controls (983 males, median age = 13 y; range= 4 y to 63 y). The previous study by Douglas *et al.* (14) (see introduction) included five datasets that were also analyzed in the present study (**ST1**).

For all but 4 of the 39 datasets, ADHD diagnosis was based on the Diagnostic and Statistical Manual of Mental Disorders 4^th^ Edition (DSM-IV) (34). Other instruments used were DSM5^th^ Edition (DSM-5), or the International Classification of Diseases (ICD)10^th^ Edition) (35). For information per dataset see **ST1**.

All participating sites had obtained ethical approval from local institutional review boards, and all datasets involved informed consent procedures for all participants.

In terms of age groups, for children (<15 y) there were subcortical data from 843 cases and 928 controls, and cortical data from 953 cases and 1,036 controls; for adolescents (15 y – 21 y) there were subcortical data from 330 cases and 234 controls, and cortical data from 412 cases and 342 controls; for adults (> 21 y) there were subcortical data from 563 cases and 492 controls, and cortical data from 613 cases and 539 controls.

Eleven additional datasets, comprising cases-only or controls-only, were excluded for the purpose of the present study (these are not listed in **ST1**). This was because our analysis models included random intercepts for ‘dataset’ (below), and diagnosis would be fully confounded with ‘dataset’ for case-only or control-only datasets.

### MRI-based measures

Structural T1-weighted brain MRI scans were acquired at each study site. Images were obtained at different field strengths (1.5 T or 3 T). An overview of the different field strengths is given in **ST1**. All sites then applied a harmonized image processing and quality-control protocol from the ENIGMA consortium (http://enigma.ini.usc.edu/protocols/imaging-protocols), which included, briefly, subcortical segmentation and cortical parcellation using the freely available and validated software FreeSurfer (versions 5.1 or 5.3) (36), with the default ‘recon-all’ pipeline, followed by visual inspection of both internal and external segmentations (**Supplementary Methods**). Exclusions on the basis of these quality control steps resulted in the sample sizes given above. The data analyzed in the current study were left and right volumes of seven bilaterally paired subcortical structures, and thickness and surface area measures for each of 34 bilaterally paired cortical regions, the latter as defined by the Desikan-Killiany atlas (37). In addition, the average cortical thickness and total surface area per hemisphere were analyzed.

The Desikan-Killiany atlas (37) was derived from manual segmentations of reference brain images. The labeling system incorporates hemisphere-specific information on sulcal and gyral geometry with spatial information regarding the locations of brain structures (37). Accordingly, the mean regional asymmetries in our data might be influenced by left-right differences present in the reference dataset used for constructing the atlas. Nonetheless, this approach was appropriate for our study focused on comparing relative asymmetry between groups, rather than trying to measure absolute asymmetry. The use of a ‘real-world’ asymmetrical atlas has the advantage that regional identification is likely to be accurate for structures that are asymmetrical both in the atlas and, on average, in the study population.

### Asymmetry indexes

Left and right data per brain region and individual subject were loaded into R (version 3.5.3), and null values were removed. An asymmetry index (AI) was calculated for each subject and each paired left-right measure using the following formula: (Left-Right)/(Left+Right). Negative AIs therefore indicate a right>left asymmetry, while positive AIs indicate a left>right asymmetry. We did not divide the denominator by 2, in contrast to some previous formulations of AIs (see Introduction), but this makes no difference in terms of deriving Cohen’s *d* effect sizes and *P*-values for group comparisons. Distributions of each of the AIs in the total study sample are plotted in **SF1**.

Correlations between AI measures in the total study sample were calculated using Pearson’s R and visualized using the *corrplot* package in R (**SF2-SF4**). Most pairwise correlations between AIs were of low magnitude, with only 22 pairwise correlations either less than −0.3 or greater than 0.3, with a minimum R = −0.418 between caudal anterior cingulate surface area and superior frontal surface area, and maximum R = 0.486 between rostral middle frontal thickness and total average thickness.

### Linear mixed effects random-intercept models

#### Main analysis

Linear mixed effects analyses were performed separately for each subcortical volume AI, cortical regional surface and thickness AI, and the total hemispheric surface area and average thickness AI, using the *nlme* package in R (version 3.5.3). Analyses were conducted separately within children, adolescents, and adults, as well as on the total study sample. All models included *diagnosis* (a binary variable, i.e. case or control), *sex* (binary) and *age* (numeric) as fixed factors, and *dataset* as a random factor (39 categories for cortical data, 34 categories for subcortical data):

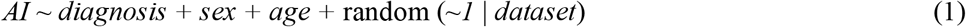

The Maximum Likelihood (ML) method was used to fit the models. Whenever any of the predictor variables was missing in a given subject, the subject was omitted from the analysis (method = ‘na.omit’).

The t-statistic for the factor ‘diagnosis’ in each linear mixed effects model was derived and used to calculate Cohen’s *d*(**Supplementary Methods**). For visualization of cerebral cortical results, Cohen’s *d* values were loaded into Matlab (v. R2019a) and 3D images of left hemisphere inflated cortical and subcortical structures were obtained using Freesurfer-derived ply files.

We did not include handedness as a covariate in our analysis, as previous studies of subcortical and cortical anatomical asymmetry in over 15,000 subjects from healthy control and population datasets, also performed by the ENIGMA consortium (30, 38), found no significant effects of handedness.

We also did not include overall brain size as a covariate, as we were interested in case-control associations with asymmetry regardless of whether they might involve other aspects of brain anatomy. In addition, in the AI formula (L-R)/(L+R), the L-R difference (numerator) is adjusted relative to the bilateral measure L+R (denominator), such that its magnitude no longer scales with L+R.

#### Sensitivity analyses

The relationships between AIs and age appeared roughly linear across all age groups combined (**SF2-4**). Therefore, no polynomials for age were incorporated in the main model (**Supplementary Methods**). However, analyses were repeated (all age groups combined) using an additional non-linear term for age, to check whether this choice had affected the results. The variables age and age^2^ would be highly correlated. To model age and non-linear effects of age in the same model, we made use of the poly()-function in R for these two predictors, which created a pair of uncorrelated variables (so-called orthogonal polynomials) (39), where one variable was linear and one non-linear:

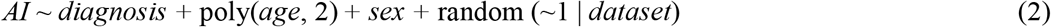

No AI outliers were removed for the main analysis, but to confirm that results were not dependent on outliers, the main analyses were also repeated (all age groups combined) after having winsorized using a threshold of k= 3, for each AI measure separately.

#### Associations between brain asymmetries and IQ, comorbidity, ADHD severity and psychostimulant medication

Within the set of ADHD participants (all age groups combined), brain asymmetries were tested in relation to several potentially associated variables (IQ, comorbidity, severity, medication use; see **SF5**), using separate models in which each variable was considered as a fixed effect:

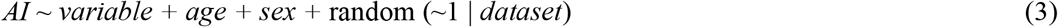

See **Supplementary Methods** for the derivation of these variables. For the dichotomous variables, i.e. comorbidities and psychostimulant medication use, datasets were removed if they had < 1 subject per category, to avoid the random variable ‘dataset’ being fully confounded with the binary variable for any datasets. Depending on the availability of each specific AI measure, data for testing association with IQ were available for up to 1,788 ADHD individuals (exact numbers per AI are in **ST18-20** and depend on image quality control for that region). For the presence/absence of comorbidities, four different binary variables were constructed: mood disorder (up to 179 yes, 384 no), anxiety disorder (up to 82 yes, 503 no), oppositional defiant disorder (ODD; up to 80 yes, 151 no), and substance use disorder (SUD; up to 77 yes, 335 no) (**ST21-23**). For ADHD symptom severity, two continuous variables were used: hyperactivity/impulsivity (up to 1,051 ADHD participants) and inattention (1,048 ADHD participants) (**ST24-26**). For psychostimulant medication use, two binary variables were constructed: lifetime use: (up to 337 yes, 188 no), and current use (up to 361 yes, 377 no) (see **SF5** for the distributions) (**ST27-29**).

IQ was also examined in controls only (all age groups combined) to explore the relationships between IQ and brain asymmetries in typically developing individuals. IQ and AI data were available for up to 1,727 controls.

We did not adjust for IQ as a covariate effect in our main, case-control analysis (further above). This was because slightly lower average IQ was part of the ADHD phenotype in our total combined dataset (Welch two-sample *t*-test for case-control difference: *t*=-12.64, df=3501.2, *P*=2.2×10^−16^) (**SF5**), so that including IQ as a covariate in case-control analysis might have reduced the power to detect an association of diagnosis with asymmetry. This would occur if underlying susceptibility factors contribute both to altered asymmetry and reduced IQ, as part of the ADHD phenotype.

#### Significance and detectable effect sizes

Significance was assessed based on the *P*-value for the diagnosis term within each model. Within each age group, and again within all age groups combined, the False Discovery Rate (FDR) (40) was estimated separately for the seven subcortical structures, for the 35 cortical surface area AIs (i.e. 34 regional AIs and one hemispheric total AI), and again for the 35 cortical thickness AIs, each time with an FDR threshold of 0.05.

As each linear model included multiple predictor variables, the power to detect an effect of diagnosis on AI could not be computed exactly, but we obtained an indication of the effect size that would be needed to provide 80% power had we been using simple t-tests and Bonferroni correction for multiple testing, using the *pwr* command in R (**Supplementary Methods**). For this purpose, a significance level of 0.0071 (i.e., 0.05/7) or 0.0014 (i.e. 0.05/35) was set in the context of multiple testing over the seven subcortical volumes, or the regional and total cortical surface areas (N = 35) or thicknesses (N = 35). This showed that, in the total study sample, a case-control effect size of roughly Cohen’s *d*=0.12 (subcortical), or *d*=0.13 (cortical), would be detectable with 80% power. For the analyses in the different age groups, this was, respectively, *d*=0.17 and *d*=0.18 in children, *d*=0.30 and *d*=0.30 in adolescents, and *d*=0.22 and *d*=0.24 in adults.

#### Directions of asymmetry changes

For any AIs showing nominally significant effects (i.e., unadjusted P<0.05) of diagnosis in any of the primary analyses, *post hoc* linear mixed effects modelling was also performed on the corresponding L and R measures separately, to understand the unilateral changes involved. The models included the same terms as were used in the main analysis of AIs (i.e., diagnosis, age and sex as fixed factors, and dataset as random factor). Again, the Cohen’s *d* effect sizes for diagnosis were calculated based on the *t*-statistics. The raw mean AI values were calculated separately in controls and cases, to describe the reference direction of healthy asymmetry in controls, and whether cases showed lower, higher, or reversed asymmetry relative to controls.

## Results

### Associations of brain asymmetry with ADHD

#### Children

There were no associations of diagnosis with AIs that had FDR <0.05 in children (**T1-T3, ST2-ST4**). The children showed nominally significant associations (unadjusted P<0.05) of diagnosis with the AIs of total hemispheric surface area (*t*=2.4, *P*=0.019) and medial orbitofrontal cortex surface area (*t*=2.7, *P*=0.007) (**T2, ST3**). The Cohen’s *d* for these effects were 0.11 and 0.12 respectively (**F1, SF6, ST3**). *Post hoc* analysis showed that both of these effects involved reductions of rightward asymmetry compared to controls, driven by a relatively greater reduction of area on the right-side than left-side in ADHD compared to controls (**ST14**).

The children also showed nominally significant associations of diagnosis with four regional cortical thickness AIs, which were the banks of the superior temporal sulcus (*t*=-2.1, *P*=0.04; increased rightward asymmetry), caudal middle frontal cortex (*t*=2.08, *P*=0.037; increased leftward asymmetry), and precentral gyrus (*t*=2.6, *P*=0.008; increased leftward asymmetry). Decreased leftward asymmetry in ADHD was observed for insula thickness AI (*t*=-2.1, *P*=0.038) (**T2, ST14**).

#### Adolescents

There were two nominally significant associations between diagnosis and AIs in adolescents, but none with FDR <0.05 (**T1-T3, ST5-ST7**).These involved the *pars orbitalis* of inferior frontal gyrus surface area (*t*=2.3, *P*=0.022), which showed lower rightward asymmetry in ADHD compared to controls, and cuneus thickness (*t*=-2.0, *P*=0.044), which showed higher rightward asymmetry in ADHD compared to controls.

#### Adults

In adults, the globus pallidus AI was significantly associated with ADHD diagnosis with FDR <0.05 (*t*=-3.1, *P*=0.002, uncorrected) (**T1, ST8**). The Cohen’s *d* effect size for this association was −0.18 (**T1, F1, SF6**). This effect involved a decrease in leftward asymmetry in ADHD compared to controls, driven by a reduction of left-side volume accompanied by an increase in right-side volume, in ADHD compared to controls (left: *d*=-0.048, 95% CI [-0.17, 0.07]; right: *d*=0.124, 95% CI [0.002, 0.24]) **(ST14)**. Note this association was only significant in the context of FDR correction for 7 subcortical AIs in adults, but would not be significant with further correction for study-wide multiple testing.

There were other nominally significant associations of AIs with diagnosis in adults: lateral occipital cortex surface area (*t*=2.0, *P*=0.048; increased leftward) (**T2, ST9, ST14**) and thickness (*t*=2.3, *P*=0.021; decreased rightward) (**T3, ST10, ST14**), medial orbitofrontal cortex thickness (*t*=2.0, *P*=0.043; increased leftward), middle temporal gyrus thickness (*t*=-2.6, *P*=0.011; increased rightward), pericalcarine cortex thickness (*t*=2.8, *P*=0.005; decreased rightward), and postcentral gyrus thickness (*t*=-2.5, *P*=0.012; decreased leftward).

#### All age groups combined

When combining all age groups, and using the same model as in the separate age groups (i.e. *AI ~ diagnosis + sex + age +* random (~1 | *dataset*)), there were nominally significant associations of AIs with diagnosis for the medial orbitofrontal cortex surface area (*t*=2.2, *P*=0.031; decreased rightward), thicknesses of the banks of the superior temporal sulcus (*t*=-2.0, *P*=0.045; increased rightward), caudal middle frontal cortex (*t*=2.3, *P*=0.024; increased leftward), insula (*t*=-2.1, *P*=0.040; decreased leftward), and precentral gyrus (*t*=2.1, *P*=0.033; increased leftward), as well as the volume of the globus pallidus (*t*=-2.1, *P*=0.033; decreased leftward) (**T1-T3, ST11-ST13)**.

The addition of non-linear effects of age to the model had negligible influences on the six nominally significant associations with diagnosis in the total study sample, all of which remained nominally significant. Winsorizing outliers (using a threshold k=3, see **Methods**) also had negligible influences on the results (**ST15-ST17**).

### Associations of brain asymmetries with IQ, comorbidity, ADHD severity, and psychostimulant medication

These analyses were carried out in all age groups combined. When testing associations of IQ, comorbidity, ADHD severity, and psychostimulant medication with brain asymmetries within ADHD individuals (**ST21-29**), only one significant (FDR <0.05) association was found, namely, between mood disorder and the rostral middle frontal gyrus thickness AI (*P*=0.0002, *t*=3.70) (**ST18**). Furthermore, various nominally significant (P<0.05) associations were observed:

IQ was associated with the accumbens volume AI (*t*=2.27, *P*=0.02), hippocampus volume AI (*t=*-1.97, *P*=0.05) (**ST18**) and insula thickness AI (*t*=2.04, *P*=0.04) (**ST20**).

ADHD severity was associated with the AI of the entorhinal cortex surface area (*t*=2.12, *P*=0.03; hyperactivity/impulsivity) (**ST25**). ADHD severity was also associated with three regional cortical thickness asymmetries: the caudal anterior cingulate thickness AI (*t*=2.66, *P*=0.01; hyperactivity/impulsivity), *pars opercularis* of the inferior frontal gyrus thickness AI (*t*=2.12, *P*=0.03; hyperactivity/impulsivity, and *t*=2.04, *P*=0.04; inattention), and pericalcarine cortex thickness AI (*t*=2.41, *P*=0.02; hyperactivity/impulsivity) (**ST26**).

Current psychostimulant medication use was associated with two cortical regional surface area asymmetries, i.e., precuneus (*t*=-2.25, *P*=0.02) and transverse temporal gyrus (*t*=-2.34, *P*=0.02) (**ST28**), and with two thickness asymmetries, i.e., inferior parietal cortex (*t*=-2.33, P=0.02) and precentral gyrus (*t*=-2.16, *P*=0.03) (**ST29**). Lifetime psychostimulant medication use was associated with three cortical surface area asymmetries (insula (*t*=-2.03, *P*=0.04), supramarginal gyrus (*t*=-2.08, *P*=0.04), and rostral anterior cingulate cortex (*t*=1.97, *P*=0.05) (**ST28**), and the thickness asymmetry of the paracentral lobule (*t*=2.16, *P*=0.03) (**ST29**). Among the AIs that showed nominally significant associations with medication use, one had also shown nominally significant association with diagnosis in all age groups combined, i.e., the AI of precentral gyrus thickness (see above). The direction of medication effect was negative, i.e. the opposite to the effect of diagnosis on this AI (see above).

For mood disorder, associations were observed with six thickness AIs (i.e., entorhinal cortex, pars triangularis of inferior frontal gyrus, pericalcarine cortex, precuneus, rostral middle frontal gyrus, and transverse temporal gyrus), and two surface area AIs (i.e., inferior temporal gyrus, and rostral anterior cingulate cortex), of which the association with rostral middle frontal thickness AI survived multiple testing correction (FDR < 0.05) (**ST22, 23**). Anxiety Disorder was associated with thickness AIs of the cuneus and lateral occipital cortex (**ST23**). For ODD, associations were found with the AIs of medial orbitofrontal thickness (**ST23**) and temporal pole surface area (**ST22**). Finally, SUD was associated with the thickness AIs of the cuneus and paracentral lobule (**ST23**), and with surface area AIs of the postcentral gyrus and supramarginal gyrus (**ST22**). None of these regions showed a nominally significant effect of diagnosis in the main analysis of all age groups combined.

Within controls, IQ was associated with the middle temporal gyrus surface area AI (*t*=-2.43, *P*=0.02) (**ST19**), rostral anterior cingulate thickness cortex AI (*t*=2.31, *P*=0.02), supramarginal gyrus thickness AI (*t* = −2.23, *P*=0.03) (**ST20**), and globus pallidus volume AI (*t*=-2.05, *P*=0.04) (**ST18**). As noted in the Methods, we did not analyze IQ-AI associations in cases and controls together, as IQ was associated with case-control status.

## Discussion

We conducted the largest study to date of associations between anatomical brain asymmetries and ADHD. Linear mixed effects model mega-analyses were carried out separately in children, adolescents, and adults, following previous ENIGMA ADHD working group studies of bilateral brain differences that showed contrasting effects in these age groups (32, 33). We also analyzed the total study sample for age-general effects. All statistical effects of diagnosis on asymmetries were very small, with Cohen’s *d* ranging from −0.18 to 0.18. Only one of these associations was significant with a false discovery rate <0.05 (globus pallidus asymmetry in adults), and this would not be significant with study-wide correction for multiple testing. Therefore, all effects remain tentative, even in this unprecedented sample size. The small effect sizes mean that altered brain asymmetry is unlikely, in itself, to be a useful biomarker or clinical predictor of ADHD. Furthermore, effect sizes reported in prior studies, based on much smaller samples, may have been unrealistically large. Low power not only reduces the chance of detecting true effects, but also increases the likelihood that statistically significant results do not reflect true effects (41).

However, there were some notable associations of diagnosis with cortical asymmetry that reached nominal significance in our study. Among these, children with ADHD showed lower rightward asymmetry of total hemispheric surface area, and medial orbitofrontal surface area. In a recent ENIGMA consortium study of autism spectrum disorder (ASD), medial orbitofrontal cortex surface area asymmetry was altered in the same direction, and to a similar extent, as in the present study (31). ADHD and ASD often co-occur (42) and are known to share genetic influences (43, 44), such that the two diagnostic labels are likely to capture a partly overlapping spectrum of related disorders (45, 46). Studies that aimed to identify shared brain structural traits between ADHD and ASD have found mixed results (47, 48), with perhaps the greatest overlap involving regions of the ‘social brain’, including orbitofrontal cortex (49). However, laterality has not been specifically studied in this regard, so that our finding of reduced rightward medial orbitofrontal cortex surface area in both disorders may be a new insight into shared neurobiology between ADHD and ASD. Altered lateralized neurodevelopment may play a causal role in disorder susceptibility, or else may arise as a correlated trait due to other underlying susceptibility factors, or even be a downstream consequence of having the disorder (50). Some aspects of brain asymmetry are partly heritable (30, 38), so that future gene mapping studies for brain asymmetry and disorder susceptibility may help to resolve causal relations underlying their associations.

One functional imaging study (94 cases, 85 controls) reported lower rightward lateralization in medial orbitofrontal cortex in ADHD compared to controls, based on temporal variability during resting-state (21). Furthermore, a study of 218 participants with ADHD and 358 healthy controls reported that orbitofrontal cortex thickness, but not surface area, showed a left>right asymmetry in childhood controls that switched to right>left asymmetry by late adolescence, while this change did not occur to the same extent in ADHD (51). However, in the present study, we saw no effect of diagnosis on this regional thickness asymmetry, but rather its surface area asymmetry. For other cortical asymmetries too, our findings in this large-scale study were discrepant with what might have been expected from previous reports in smaller samples (see references in the Introduction). For example, prior studies have reported reversed grey matter volume asymmetry (i.e., leftward instead of rightward) in the superior frontal and middle frontal gyri in ADHD (26), and decreased leftward asymmetry of cortical convolution complexity in prefrontal cortex, as compared to healthy controls (27).

The most often reported alteration of brain asymmetry in ADHD has involved the caudate nucleus, although the direction of the effect has not been consistent (9, 10, 22-25). We did not find evidence for altered asymmetry of caudate nucleus volume in the present study, again suggesting that prior findings were false positives in smaller samples. We found a significant association with diagnosis for another region of the basal ganglia, namely the globus pallidus, in adults-only, although this also remained nominally significant in all groups combined. The globus pallidus is involved in movement and reward processing (52), both of which are involved in the symptomatology of ADHD. A previous meta-analysis comprising data from a total of 114 participants with ADHD (or a related disorder) and 143 control participants, noted a significantly lower average right putamen and right globus pallidus volumes in ADHD (53), although asymmetry was not quantified. Regardless, our finding of lower leftward asymmetry seems discrepant with this earlier report.

We have already remarked on the limited statistical power of previous studies as a likely explanation for their findings being discrepant with the current study. Low sample sizes in relation to subtle effects can result in poor reproducibility (41, 54). Here, we had 80% power to detect case-control Cohen’s *d* effect sizes as low as roughly 0.12, or as high as 0.3 in the smallest subset by age (see **Methods**). In addition to limited sample sizes, there are various other possible explanations for discrepancies with previous studies. Methodological differences in hardware, software, and data processing pipelines can influence results (55). In terms of brain structural quantification, the cortical atlas that we used did not have perfect equivalents for some regions or measures defined in many of the earlier studies. For example, we did not consider gyral/sulcal patterns or cortical grey matter volumes as such. Rather, we studied regional cortical thicknesses and surface areas as distinct measures, which together drive grey matter volumetric measures. Since area and thickness have been shown to vary relatively independently (56), separate analyses are advisable.

The conceptualization of laterality can also differ across studies. In terms of AIs, our cortical results are largely in line with a previous report based on measuring grey matter volume asymmetries in 192 participants with ADHD and 508 controls (14), insofar as no FDR-significant results were found (five of those datasets were in common with the present study, see **Methods**). However, the authors of that study also calculated the unsigned magnitudes of the AIs (i.e., absolute degrees of asymmetry, regardless of directions). They reported significant differences in absolute asymmetry for various cortical and subcortical structures (14). In the present study, we did not calculate absolute AIs, in order not to compound multiple testing, and because these measures are highly non-normal with a floor effect at value zero, which would violate the assumptions of the modelling that we applied. It is not clear whether this issue may have affected the results in the earlier study (14). Future studies may consider the unsigned magnitude of brain asymmetry indexes further in ADHD, but it will be necessary to use statistical methods that can account for non-normal distributions. Discrepancies with earlier studies may also be due to differences in clinical features of the disorder that arise from case recruitment and diagnosis, for example with respect to medication use (57, 58), comorbidities (59), symptom severity, and/or IQ. Some asymmetries showed tentative associations with some of these clinical variables in the present study, although none of these results survived correction for multiple testing, apart from mood disorder with the rostral middle frontal thickness AI. Also, some of the clinical variables (medication, comorbidity) were missing for many ADHD individuals in this study. Regardless, it remains possible that certain subsets of ADHD might be associated more strongly with altered brain asymmetry than was apparent in our large-scale analysis of average changes over many datasets, comprising many and varied collections of ADHD individuals and controls.

In general, between-centre heterogeneity (in terms of methods used, patient subgroups, demographics) may result in reduced statistical power to detect effects that are specific to certain subgroups of datasets, or to individual datasets, when tested in mega-analysis over all datasets. Here we used random-intercept models to adjust for heterogeneity between datasets, but this cannot fully rescue power in the case that effects are truly specific to certain subsets. However, no single centre has been able to collect such a large ADHD-control sample alone, while our large sample size yields more precise estimates of effect sizes with respect to the overall case-control population, as represented across many research centres. In this way, the findings from multi-centre studies such as ours can be considered more generalizable than single-centre studies (60). In any case, as long as researchers publish separate papers based on many single, smaller datasets, collected in particular ways, the field overall has the same issue of heterogeneity.

Although not a longitudinal study, our data spanned a wide age range from childhood through to older adulthood, which allowed us to study different age groups separately, as the disorder may be neurobiologically distinct in different age groups (32, 61). The previous ENIGMA study of bilateral cortical differences in ADHD found children to be most affected, particularly in frontal, cingulate, and temporal regions, as well as the total hemispheric surface area, which was lower in ADHD (32). In the children-only analysis in our present study of asymmetries, we also found associations with diagnosis for some frontal and temporal regions (caudal middle frontal cortex thickness, precentral gyrus thickness, medial orbitofrontal cortex surface area, banks of the superior temporal sulcus thickness), as well as a change in the asymmetry of total hemispheric surface area, driven by a greater decrease of area in ADHD on the right-side than the left-side. These findings offer a more nuanced description of brain changes in childhood ADHD, which may involve altered lateralized neurodevelopment.

Considering all brain asymmetry measures, the effect sizes in the present study were not stronger in children as compared to adolescents or adults. Furthermore, bilateral case-control differences are not necessarily a good guide to case-control differences in asymmetry, since a difference in asymmetry can arise, for example, from a simultaneous left-sided increase and right-sided decrease in a brain measure, which can involve no change at all in the bilateral measure. Hence, we took a screening approach to the present study, rather than constraining our search on prior bilateral findings. It is also not entirely clear how/whether to statistically adjust the test for total hemispheric surface asymmetry, in the context of also testing multiple sub-regions, and also with respect to study-wide multiple testing. Therefore, we present all *P*-values unadjusted, while also being mindful of the tentative nature of these findings in the context of our survey across many brain asymmetry measures. Together with the corresponding effect size estimates, this mapping information should be useful for the field. In summary, we have carried out the largest case-control study of structural brain asymmetry in ADHD. We describe average changes of asymmetry that are small, but helpful towards a more complete description of brain anatomical changes in this disorder. Results were largely discrepant with earlier, inconsistent findings from smaller-scale studies, which illustrates the value of taking a large-scale approach to human clinical neuroscience. The small effects that we found remain statistically tentative in the context of multiple testing, even in this unprecedented sample size. Future longitudinal and genetic studies may probe causative relations between ADHD and brain asymmetry, focused on measures defined from this study, such as total hemispheric surface area asymmetry, medial orbitofrontal area asymmetry, or globus pallidus volume asymmetry.

## Acknowledgments

Data were made available for this study by participants of the ENIGMA-ADHD working group (http://enigma.ini.usc.edu/ongoing/enigma-adhd-working-group/). Acknowledgements in relation to the many separate research groups and datasets are listed in the Supplementary Information.

---

**Table 1.** Linear mixed model results for subcortical volume AIs.

| Subcortical volume AI | Children only |  | Adolescents only |  | Adults only |  | Total study sample |  |
| --- | --- | --- | --- | --- | --- | --- | --- | --- |
|  | P<sup>1</sup> | d<sup>2</sup> | P<sup>1</sup> | d<sup>2</sup> | P<sup>1</sup> | d<sup>2</sup> | P<sup>1</sup> | d<sup>2</sup> |
| Accumbens | 0.22 | -0.06 | 0.40 | -0.07 | 0.94 | 0.00 | 0.29 | -0.04 |
| Amygdala | 0.96 | -0.002 | 0.64 | 0.04 | 0.74 | -0.02 | 0.79 | -0.01 |
| Caudate Nucleus | 0.66 | 0.02 | 0.93 | 0.01 | 0.40 | 0.05 | 0.44 | 0.03 |
| Globus Pallidus | 0.91 | 0.01 | 0.44 | -0.07 | **0.003** | -0.18 | **0.03** | -0.07 |
| Hippocampus | 0.71 | -0.02 | 0.23 | 0.11 | 0.46 | 0.05 | 0.71 | 0.01 |
| Putamen | 0.37 | -0.04 | 0.75 | -0.03 | 0.53 | -0.04 | 0.16 | -0.05 |
| Thalamus* | 0.47 | 0.04 | 0.28 | 0.10 | 0.49 | 0.04 | 0.20 | 0.05 |

<sup>1</sup> Uncorrected P-values for diagnosis are indicated, with in **bold** those that are significant at the uncorrected level ( $P < 0.05$ ), and in **bold-italic** those that survive multiple testing correction. <sup>2</sup> Cohen's *d* value for the effect of diagnosis. \*Thalamus volume was not available from the NIH sample.

**Table 2.** Linear mixed model results for the cortical surface area AIs.

| Cortical surface area AI | Children only |  | Adolescents only |  | Adults only |  | Total study sample |  |
| --- | --- | --- | --- | --- | --- | --- | --- | --- |
|  | P<sup>1</sup> | d<sup>2</sup> | P<sup>1</sup> | d<sup>2</sup> | P<sup>1</sup> | d<sup>2</sup> | P<sup>1</sup> | d<sup>2</sup> |
| Banks of superior temporal sulcus | 0.68 | -0.02 | 0.26 | -0.09 | 0.82 | 0.01 | 0.38 | -0.03 |
| Caudal anterior cingulate cortex | 0.65 | -0.02 | 0.27 | -0.08 | 0.71 | 0.02 | 0.58 | -0.02 |
| Caudal middle frontal cortex | 0.37 | 0.04 | 0.82 | -0.02 | 0.22 | 0.07 | 0.17 | 0.04 |
| Cuneus | 0.10 | 0.07 | 0.89 | 0.01 | 0.07 | -0.11 | 0.87 | -0.01 |
| Entorhinal cortex | 0.84 | -0.01 | 0.66 | -0.03 | 0.10 | -0.10 | 0.27 | -0.04 |
| Frontal pole | 0.07 | -0.08 | 0.42 | 0.06 | 0.25 | -0.07 | 0.12 | -0.05 |
| Fusiform gyrus | 0.14 | -0.07 | 0.32 | 0.07 | 0.11 | -0.10 | 0.12 | -0.05 |
| Inferior parietal cortex | 0.16 | 0.06 | 0.94 | -0.01 | 0.89 | -0.01 | 0.31 | 0.03 |
| Inferior temporal gyrus | 0.40 | 0.04 | 0.95 | 0.00 | 0.26 | 0.07 | 0.19 | 0.04 |
| Insula | 0.07 | 0.08 | 0.73 | 0.03 | 0.62 | -0.03 | 0.24 | 0.04 |
| Isthmus cingulate cortex | 1.00 | 0.00 | 0.08 | -0.13 | 0.48 | 0.04 | 0.72 | -0.01 |
| Lateral occipital cortex | 0.38 | -0.04 | 0.82 | 0.02 | 0.05 | 0.12 | 0.66 | 0.01 |
| Lateral orbitofrontal cortex | 0.27 | -0.05 | 0.46 | -0.06 | 0.40 | -0.05 | 0.09 | -0.06 |
| Lingual gyrus | 0.85 | 0.01 | 0.17 | -0.10 | 0.49 | 0.04 | 0.89 | 0.00 |
| Medial orbitofrontal cortex | **0.01** | 0.12 | 0.35 | 0.07 | 0.77 | -0.02 | **0.03** | 0.07 |
| Middle temporal gyrus | 0.16 | 0.07 | 0.63 | -0.04 | 0.93 | -0.01 | 0.44 | 0.03 |
| Paracentral lobule | 0.07 | -0.08 | 0.98 | 0.00 | 0.29 | -0.06 | 0.06 | -0.06 |
| Parahippocampal gyrus | 0.29 | 0.05 | 0.43 | -0.06 | 0.13 | -0.09 | 0.85 | -0.01 |
| Pars opercularis of inferior frontal gyrus | 0.88 | -0.01 | 0.16 | 0.11 | 0.57 | 0.03 | 0.46 | 0.02 |
| Pars orbitalis of inferior frontal gyrus | 0.44 | 0.04 | **0.02** | 0.18 | 0.54 | 0.04 | 0.05 | 0.06 |
| Pars triangularis of inferior frontal gyrus | 0.30 | 0.05 | 0.13 | 0.11 | 0.57 | -0.03 | 0.22 | 0.04 |
| Pericalcarine cortex | 0.22 | 0.06 | 0.11 | -0.12 | 1.00 | 0.00 | 0.84 | 0.01 |
| Postcentral gyrus | 0.45 | 0.03 | 0.42 | 0.06 | 0.98 | 0.00 | 0.37 | 0.03 |
| Posterior cingulate cortex | 0.98 | 0.00 | 0.48 | -0.05 | 0.82 | 0.01 | 0.88 | -0.01 |
| Precentral gyrus | 0.80 | 0.01 | 0.07 | -0.14 | 0.06 | -0.11 | 0.09 | -0.05 |
| Precuneus | 0.37 | 0.04 | 0.29 | -0.08 | 0.65 | 0.03 | 0.57 | 0.02 |
| Rostral anterior cingulate cortex | 0.89 | -0.01 | 0.93 | -0.01 | 0.36 | -0.05 | 0.46 | -0.02 |
| Rostral middle frontal gyrus | 0.20 | -0.06 | 0.89 | -0.01 | 0.60 | -0.03 | 0.21 | -0.04 |
| Superior frontal gyrus | 0.29 | 0.05 | 0.07 | 0.13 | 0.12 | -0.09 | 0.54 | 0.02 |
| Superior parietal cortex | 0.15 | 0.07 | 0.47 | 0.05 | 0.68 | -0.02 | 0.34 | 0.03 |
| Superior temporal gyrus | 0.36 | 0.04 | 0.91 | 0.01 | 0.20 | -0.08 | 0.99 | 0.00 |
| Supramarginal gyrus | 0.79 | 0.01 | 0.18 | -0.10 | 0.22 | -0.07 | 0.28 | -0.04 |
| Temporal pole | 0.40 | 0.04 | 0.46 | 0.06 | 0.36 | -0.06 | 0.73 | 0.01 |
| Transverse temporal gyrus | 0.56 | -0.03 | 0.39 | 0.07 | 0.94 | 0.00 | 0.98 | -0.0008 |
| Total average surface area | **0.02** | 0.11 | 0.90 | 0.01 | 0.24 | -0.07 | 0.43 | 0.03 |

<sup>1</sup> Uncorrected P-values for diagnosis are indicated, with in **bold** those that are significant at the uncorrected level (P < 0.05). None survived multiple testing correction. <sup>2</sup> Cohen's *d* value for the effect of diagnosis.

**Table 3.** Linear mixed model results for the cortical thickness AIs.

| Cortical thickness AI | Children only |  | Adolescents only |  | Adults only |  | Total study sample |  |
| --- | --- | --- | --- | --- | --- | --- | --- | --- |
|  | P<sup>1</sup> | d<sup>2</sup> | P<sup>1</sup> | d<sup>2</sup> | P<sup>1</sup> | d<sup>2</sup> | P<sup>1</sup> | d<sup>2</sup> |
| Banks of superior temporal sulcus | 0.04 | -0.10 | 0.48 | -0.06 | 0.64 | -0.03 | **0.05** | -0.07 |
| Caudal anterior cingulate cortex | 0.32 | 0.05 | 0.65 | -0.03 | 0.06 | 0.11 | 0.15 | 0.05 |
| Caudal middle frontal cortex | 0.04 | 0.09 | 0.06 | 0.14 | 0.77 | 0.02 | **0.02** | 0.07 |
| Cuneus | 0.63 | 0.02 | 0.05 | -0.15 | 0.07 | 0.11 | 0.59 | 0.02 |
| Entorhinal cortex | 0.14 | -0.07 | 0.86 | 0.01 | 0.63 | -0.03 | 0.27 | -0.04 |
| Frontal pole | 0.22 | 0.05 | 0.19 | -0.10 | 0.19 | 0.08 | 0.31 | 0.03 |
| Fusiform gyrus | 0.44 | -0.04 | 0.91 | 0.01 | 0.75 | 0.02 | 0.85 | -0.01 |
| Inferior parietal cortex | 0.81 | -0.01 | 0.52 | -0.05 | 0.54 | 0.04 | 0.92 | 0.00 |
| Inferior temporal gyrus | 0.30 | -0.05 | 0.58 | 0.04 | 0.87 | -0.01 | 0.77 | -0.01 |
| Insula | **0.04** | -0.09 | 0.19 | -0.10 | 0.93 | -0.01 | **0.04** | -0.07 |
| Isthmus cingulate cortex | 0.53 | -0.03 | 0.18 | 0.10 | 0.34 | -0.06 | 0.73 | -0.01 |
| Lateral occipital cortex | 0.55 | 0.03 | 0.22 | -0.09 | **0.03** | 0.13 | 0.33 | 0.03 |
| Lateral orbitofrontal cortex | 0.73 | -0.02 | 0.50 | 0.05 | 0.14 | 0.09 | 0.42 | 0.03 |
| Lingual gyrus | 0.41 | -0.04 | 0.97 | 0.003 | 0.60 | -0.03 | 0.34 | -0.03 |
| Medial orbitofrontal cortex | 0.08 | -0.08 | 0.23 | 0.09 | **0.04** | 0.12 | 0.82 | 0.01 |
| Middle temporal gyrus | 0.83 | -0.01 | 0.74 | -0.03 | 0.01 | -0.16 | 0.15 | -0.05 |
| Paracentral lobule | 0.11 | -0.07 | 0.18 | 0.10 | 0.77 | -0.02 | 0.39 | -0.03 |
| Parahippocampal gyrus | 0.14 | -0.07 | 0.06 | -0.14 | 0.40 | 0.05 | 0.18 | -0.04 |
| Pars opercularis of inferior frontal gyrus | 0.86 | 0.01 | 0.26 | 0.08 | 0.86 | -0.01 | 0.44 | 0.03 |
| Pars orbitalis of inferior frontal gyrus | 0.27 | 0.05 | 0.93 | -0.01 | 0.39 | 0.05 | 0.25 | 0.04 |
| Pars triangularis of inferior frontal gyrus | 0.84 | -0.01 | 0.40 | 0.06 | 0.90 | -0.01 | 0.72 | 0.01 |
| Pericalcarine cortex | 0.89 | 0.01 | 0.95 | 0.00 | **0.004** | 0.17 | 0.11 | 0.05 |
| Postcentral gyrus | 0.79 | -0.01 | 0.79 | -0.02 | 0.01 | -0.15 | 0.08 | -0.06 |
| Posterior cingulate cortex | 0.64 | -0.02 | 0.54 | -0.05 | 0.84 | -0.01 | 0.34 | -0.03 |
| Precentral gyrus | **0.01** | 0.12 | 0.39 | -0.06 | 0.16 | 0.08 | **0.03** | 0.07 |
| Precuneus | 0.80 | 0.01 | 0.17 | 0.10 | 0.73 | 0.02 | 0.45 | 0.02 |
| Rostral anterior cingulate cortex | 0.82 | -0.01 | 0.09 | 0.13 | 0.35 | 0.06 | 0.25 | 0.04 |
| Rostral middle frontal gyrus | 0.77 | 0.01 | 0.78 | -0.02 | 0.39 | -0.05 | 0.82 | -0.01 |
| Superior frontal gyrus | 0.64 | 0.02 | 0.08 | 0.13 | 0.56 | 0.04 | 0.20 | 0.04 |
| Superior parietal cortex | 0.85 | 0.01 | 0.48 | -0.05 | 0.88 | 0.01 | 0.82 | -0.01 |
| Superior temporal gyrus | 0.08 | 0.08 | 0.40 | 0.07 | 0.37 | -0.06 | 0.24 | 0.04 |
| Supramarginal gyrus | 0.16 | -0.06 | 0.41 | -0.06 | 0.93 | -0.01 | 0.15 | -0.05 |
| Temporal pole | 0.53 | 0.03 | 0.66 | 0.03 | 0.63 | -0.03 | 0.71 | 0.01 |
| Transverse temporal gyrus | 0.55 | 0.03 | 0.72 | 0.03 | 0.34 | -0.06 | 0.90 | 0.00 |
| Total average thickness | 0.92 | -0.004 | 0.75 | 0.02 | 0.78 | 0.02 | 0.79 | 0.01 |

<sup>1</sup> Uncorrected P-values for diagnosis are indicated, with in **bold** those that are significant at the uncorrected level (P < 0.05). None of the associations with diagnosis survived multiple testing correction. <sup>2</sup> Cohen's *d* value for the effect of diagnosis.

**Figure 1.**
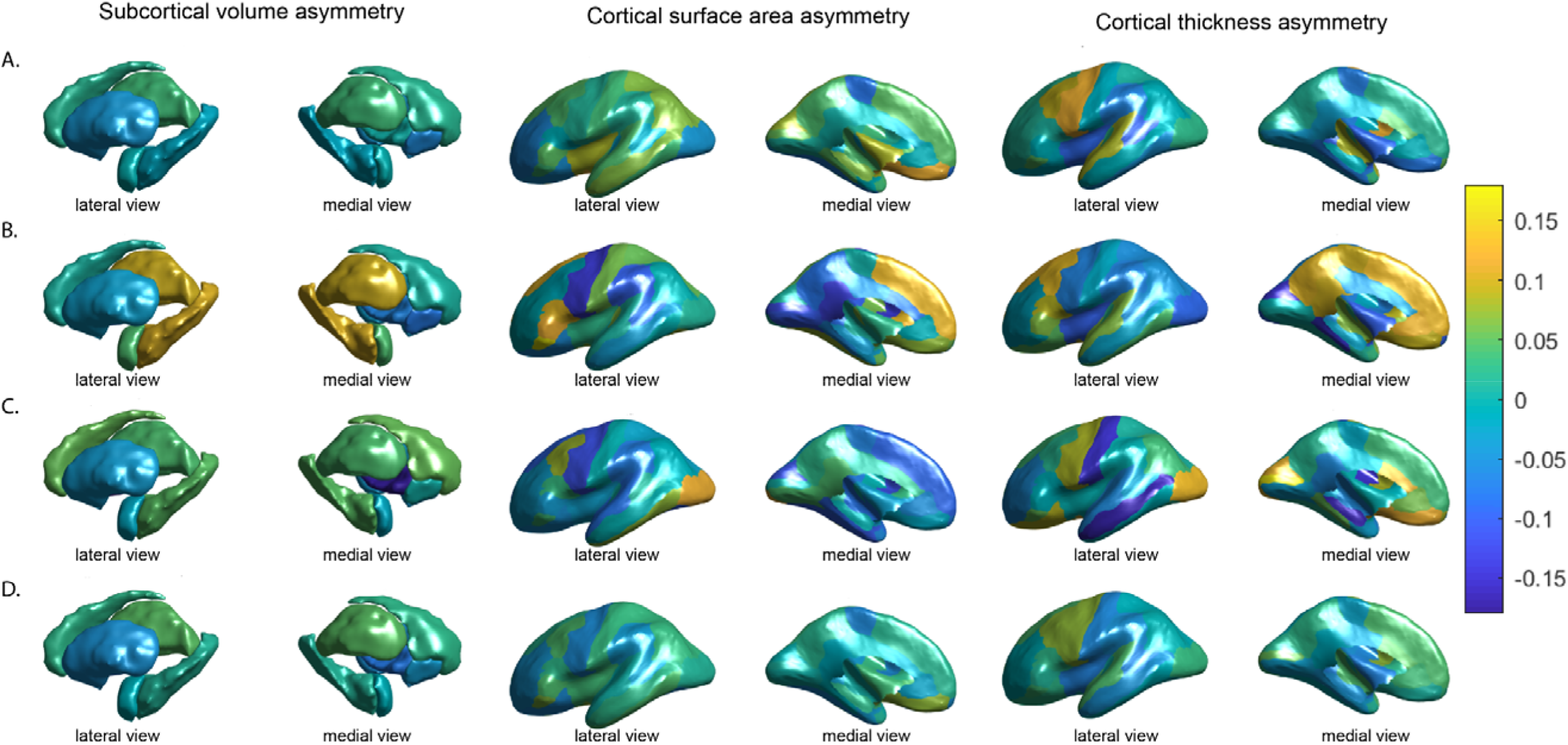
Cohen’s d effect sizes of the associations between ADHD diagnosis and AIs of subcortical volumes (left), cortical surface areas (middle) and cortical thicknesses (right) for (A) children, (B) adolescents, (C) adults, and (D) all age groups combined. Cohen’s *d* values are overlayed on left hemisphere inflated brains. Positive values (yellow) indicate mean shifts towards greater leftward or reduced rightward asymmetry in ADHD, and negative values (blue) indicate mean shifts towards greater rightward asymmetry or reduced leftward asymmetry in ADHD.

## References

1. Polanczyk G, de Lima MS, Horta BL, Biederman J, Rohde LA. The worldwide prevalence of ADHD: a systematic review and metaregression analysis. Am J Psychiatry. 2007;164(6):942–8.

2. American Psychiatric Association. Diagnostic and Statistical Manual of Mental disorders (5th ed.). Washington, DC. 2013.

3. Faraone SV, Asherson P, Banaschewski T, Biederman J, Buitelaar JK, Ramos-Quiroga JA, et al. Attention-deficit/hyperactivity disorder. Nat Rev Dis Primers. 2015;1:15020.

4. Fayyad J, Sampson NA, Hwang I, Adamowski T, Aguilar-Gaxiola S, Al-Hamzawi A, et al. The descriptive epidemiology of DSM-IV Adult ADHD in the World Health Organization World Mental Health Surveys. Attention deficit and hyperactivity disorders. 2017;9(1):47–65.

5. Renteria ME. Cerebral asymmetry: a quantitative, multifactorial, and plastic brain phenotype. Twin Res Hum Genet. 2012;15(3):401–13.

6. Toga AW, Thompson PM. Mapping brain asymmetry. Nat Rev Neurosci. 2003;4(1):37–48.

7. Duboc V, Dufourcq P, Blader P, Roussigne M. Asymmetry of the Brain: Development and Implications. Annu Rev Genet. 2015;49:647–72.

8. Heilman KM, Bowers D, Valenstein E, Watson RT. The right hemisphere: neuropsychological functions. J Neurosurg. 1986;64(5):693–704.

9. Uhlikova P, Paclt I, Vaneckova M, Morcinek T, Seidel Z, Krasensky J, et al. Asymmetry of basal ganglia in children with attention deficit hyperactivity disorder. Neuro Endocrinol Lett. 2007;28(5):604–9.

10. Schrimsher GW, Billingsley RL, Jackson EF, Moore BD, 3rd. Caudate nucleus volume asymmetry predicts attention-deficit hyperactivity disorder (ADHD) symptomatology in children. J Child Neurol. 2002;17(12):877–84.

11. Paclt I, Pribilova N, Kollarova P, Kohoutova M, Dezortova M, Hajek M, et al. Reverse asymmetry and changes in brain structural volume of the basal ganglia in ADHD, developmental changes and the impact of stimulant medications. Neuro Endocrinol Lett. 2016;37(1):29–32.

12. Li D, Li T, Niu Y, Xiang J, Cao R, Liu B, et al. Reduced hemispheric asymmetry of brain anatomical networks in attention deficit hyperactivity disorder. Brain Imaging Behav. 2018.

13. Hale TS, Smalley SL, Walshaw PD, Hanada G, Macion J, McCracken JT, et al. Atypical EEG beta asymmetry in adults with ADHD. Neuropsychologia. 2010;48(12):3532–9.

14. Douglas PK, Gutman B, Anderson A, Larios C, Lawrence KE, Narr K, et al. Hemispheric brain asymmetry differences in youths with attention-deficit/hyperactivity disorder. NeuroImage Clin. 2018;18:744–52.

15. Stefanatos GA, Wasserstein J. Attention deficit/hyperactivity disorder as a right hemisphere syndrome. Selective literature review and detailed neuropsychological case studies. Ann N Y Acad Sci. 2001;931:172–95.

16. Geeraerts S, Lafosse C, Vaes N, Vandenbussche E, Verfaillie K. Dysfunction of right-hemisphere attentional networks in attention deficit hyperactivity disorder. J Clin Exp Neuropsychol. 2008;30(1):42–52.

17. Langleben DD, Austin G, Krikorian G, Ridlehuber HW, Goris ML, Strauss HW. Interhemispheric asymmetry of regional cerebral blood flow in prepubescent boys with attention deficit hyperactivity disorder. Nucl Med Commun. 2001;22(12):1333–40.

18. Hale TS, Kane AM, Kaminsky O, Tung KL, Wiley JF, McGough JJ, et al. Visual Network Asymmetry and Default Mode Network Function in ADHD: An fMRI Study. Front Psychiatry. 2014;5:81.

19. Cortese S, Kelly C, Chabernaud C, Proal E, Di Martino A, Milham MP, et al. Toward systems neuroscience of ADHD: a meta-analysis of 55 fMRI studies. Am J Psychiatry. 2012;169(10):1038–55.

20. Vance A, Silk TJ, Casey M, Rinehart NJ, Bradshaw JL, Bellgrove MA, et al. Right parietal dysfunction in children with attention deficit hyperactivity disorder, combined type: a functional MRI study. Mol Psychiatry. 2007;12(9):826–32, 793.

21. Zou H, Yang J. Temporal Variability-Based Functional Brain Lateralization Study in ADHD. J Atten Dis. 2019: 1087054719859074.

22. Dang LC, Samanez-Larkin GR, Young JS, Cowan RL, Kessler RM, Zald DH. Caudate asymmetry is related to attentional impulsivity and an objective measure of ADHD-like attentional problems in healthy adults. Brain Struct Funct. 2016;221(1):277–86.

23. Hynd GW, Hern KL, Novey ES, Eliopulos D, Marshall R, Gonzalez JJ, et al. Attention deficit-hyperactivity disorder and asymmetry of the caudate nucleus. J Child Neurol. 1993;8(4):339–47.

24. Filipek PA, SemrudClikeman M, Steingard RJ, Renshaw PF, Kennedy DN, Biederman J. Volumetric MRI analysis comparing subjects having attention-deficit hyperactivity disorder with normal controls. Neurology. 1997;48(3):589–601.

25. Castellanos FX, Giedd JN, Marsh WL, Hamburger SD, Vaituzis AC, Dickstein DP, et al. Quantitative brain magnetic resonance imaging in attention-deficit hyperactivity disorder. Arch Gen Psychiatry. 1996;53(7):607–16.

26. Cao Q, Wang J, Sun L, Wang P, Wu Z, Wang Y. [Altered anatomical asymmetry in children with attention deficit/hyperactivity disorder: a pilot optimized voxel-based morphometric study]. Zhonghua yi xue za zhi. 2014;94(43):3387–91.

27. Li X, Jiang J, Zhu W, Yu C, Sui M, Wang Y, et al. Asymmetry of prefrontal cortical convolution complexity in males with attention-deficit/hyperactivity disorder using fractal information dimension. Brain Dev. 2007;29(10):649–55.

28. Kurth F, Gaser C, Luders E. A 12-step user guide for analyzing voxel-wise gray matter asymmetries in statistical parametric mapping (SPM). Nat Prot. 2015;10(2):293–304.

29. Leroy F, Cai Q, Bogart SL, Dubois J, Coulon O, Monzalvo K, et al. New human-specific brain landmark: the depth asymmetry of superior temporal sulcus. Proc Natl Acad Sci U S A. 2015;112(4):1208–13.

30. Kong XZ, Mathias SR, Guadalupe T, Glahn DC, Franke B, Crivello F, et al. Mapping cortical brain asymmetry in 17,141 healthy individuals worldwide via the ENIGMA Consortium. Proc Natl Acad Sci U S A. 2018.

31. Postema MC, van Rooij D, Anagnostou E, Arango C, Auzias G, Behrmann M, et al. Altered structural brain asymmetry in autism spectrum disorder in a study of 54 datasets. Nat Commun. 2019;10(1):4958.

32. Hoogman M, Muetzel R, Guimaraes JP, Shumskaya E, Mennes M, Zwiers MP, et al. Brain Imaging of the Cortex in ADHD: A Coordinated Analysis of Large-Scale Clinical and Population-Based Samples. Am J Psychiatry. 2019: appiajp201918091033.

33. Hoogman M, Bralten J, Hibar DP, Mennes M, Zwiers MP, Schweren LS, et al. Subcortical brain volume differences in participants with attention deficit hyperactivity disorder in children and adults: a cross-sectional mega-analysis. The Lancet Psychiatry. 2017;4(4):310–9.

34. American Psychiatric Association. Diagnostic and statistical manual of mental disorders, 4th edition (DSM-IV), Washington DC. 2000.

35. World Health Organization. International Classification of Diseases and Related Health Problems, 10th Revision. World Health Organization: Geneva. 1992.

36. Fischl B. FreeSurfer. NeuroImage. 2012;62(2):774–81.

37. Desikan RS, Segonne F, Fischl B, Quinn BT, Dickerson BC, Blacker D, et al. An automated labeling system for subdividing the human cerebral cortex on MRI scans into gyral based regions of interest. Neuroimage. 2006;31(3):968–80.

38. Guadalupe T, Mathias SR, vanErp TG, Whelan CD, Zwiers MP, Abe Y, et al. Human subcortical brain asymmetries in 15,847 people worldwide reveal effects of age and sex. Brain Imaging Behav. 2016.

39. Chambers JM, Hastie TJ. Statistical models in S. Pacific Grove, California, USA, Wadsworth & Brooks/Cole. 1992.

40. Benjamini Y, Hochberg Y. Controlling the False Discovery Rate - A Practical and Powerful Approach to Multiple Testing. J R Stat Soc Ser B-Methodol. 1995;57(1):289–300.

41. Munafo MR, Flint J. How reliable are scientific studies? Br J Psychiatry. 2010;197(4):257–8.

42. Leitner Y. The co-occurrence of autism and attention deficit hyperactivity disorder in children - what do we know? Front Hum Neurosci. 2014;8:268.

43. Ghirardi L, Pettersson E, Taylor MJ, Freitag CM, Franke B, Asherson P, et al. Genetic and environmental contribution to the overlap between ADHD and ASD trait dimensions in young adults: a twin study. Psychol Med. 2019;49(10):1713–21.

44. Stergiakouli E, Davey Smith G, Martin J, Skuse DH, Viechtbauer W, Ring SM, et al. Shared genetic influences between dimensional ASD and ADHD symptoms during child and adolescent development. Mol Autism. 2017;8:18.

45. van der Meer JM, Oerlemans AM, van Steijn DJ, Lappenschaar MG, de Sonneville LM, Buitelaar JK, et al. Are autism spectrum disorder and attention-deficit/hyperactivity disorder different manifestations of one overarching disorder? Cognitive and symptom evidence from a clinical and population-based sample. J Am Acad Child Adolesc Psychiatry. 2012;51(11):1160–72.e3.

46. Demopoulos C, Hopkins J, Davis A. A comparison of social cognitive profiles in children with autism spectrum disorders and attention-deficit/hyperactivity disorder: a matter of quantitative but not qualitative difference? J Autism Dev Disord. 2013;43(5):1157–70.

47. Nevena V. Radonjić, Jonathan L. Hess, Paula Rovira, Ole Andreassen, Jan K. Buitelaar, Christopher R. K. Ching, Barbara Franke, Martine Hoogman, Neda Jahanshad, Carrie McDonald, Lianne Schmaal, Sanjay M. Sisodiya, Dan J. Stein, Odile A. van den Heuvel, Theo G.M. van Erp, Daan van Rooij, Dick J. Veltman, Paul Thompson, Stephen V. Faraone. Structural Brain Imaging Studies Offer Clues about the Effects of the Shared Genetic Etiology among Neuropsychiatric Disorders. BioRxiv. 2019.

48. Premika S.W. Boedhoe MS, Daan van Rooij, Martine Hoogman, Jos W.R. Twisk, Lianne, Schmaal, Yoshinari Abe, Pino Alonso, Stephanie H. Ameis, Anatoly, Anikin, Alan Anticevic, Philip Aherson, Celso Arango, Paul D. Arnold, Guillaume Auzias, Tobias Banaschewski, Alexander, Baranov, Marcelo C. Batistuzzo, Sarah Baumeister, Ramona Baur-Streubel, Marlene, Behrmann, Mark A. Bellgrove, Francesco Benedetti, Jan C. Beucke, Joseph Biederman, et al. Subcortical brain volume, regional cortical thickness and cortical surface area across attention-deficit/hyperactivity disorder (ADHD), autism spectrum disorder (ASD), and obsessive-compulsive disorder (OCD). bioRxiv. 2019.

49. Baribeau DA, Dupuis A, Paton TA, Hammill C, Scherer SW, Schachar RJ, et al. Structural neuroimaging correlates of social deficits are similar in autism spectrum disorder and attention-deficit/hyperactivity disorder: analysis from the POND Network. Transl Psychiatry. 2019;9(1):72.

50. Bishop DV. Cerebral asymmetry and language development: cause, correlate, or consequence? Science. 2013;340(6138):1230531.

51. Shaw P, Lalonde F, Lepage C, Rabin C, Eckstrand K, Sharp W, et al. Development of cortical asymmetry in typically developing children and its disruption in attention-deficit/hyperactivity disorder. Arch Gen Psychiatry. 2009;66(8):888–96.

52. Munte TF, Marco-Pallares J, Bolat S, Heldmann M, Lutjens G, Nager W, et al. The human globus pallidus internus is sensitive to rewards - Evidence from intracerebral recordings. Brain Stim. 2017;10(3):657–63.

53. Ellison-Wright I, Ellison-Wright Z, Bullmore E. Structural brain change in Attention Deficit Hyperactivity Disorder identified by meta-analysis. BMC Psychiatry. 2008;8:51.

54. Button KS, Ioannidis JP, Mokrysz C, Nosek BA, Flint J, Robinson ES, et al. Power failure: why small sample size undermines the reliability of neuroscience. Nat Rev Neurosci. 2013;14(5):365–76.

55. Biberacher V, Schmidt P, Keshavan A, Boucard CC, Righart R, Samann P, et al. Intra- and interscanner variability of magnetic resonance imaging based volumetry in multiple sclerosis. Neuroimage. 2016;142:188–97.

56. Panizzon MS, Fennema-Notestine C, Eyler LT, Jernigan TL, Prom-Wormley E, Neale M, et al. Distinct genetic influences on cortical surface area and cortical thickness. Cereb Cortex. 2009;19(11):2728–35.

57. Nakao T, Radua J, Rubia K, Mataix-Cols D. Gray matter volume abnormalities in ADHD: voxel-based meta-analysis exploring the effects of age and stimulant medication. Am J Psychiatry. 2011;168(11):1154–63.

58. Pretus C, Ramos-Quiroga JA, Richarte V, Corrales M, Picado M, Carmona S, et al. Time and psychostimulants: Opposing long-term structural effects in the adult ADHD brain. A longitudinal MR study. Eur Neuropsychopharmacol. 2017.

59. Reale L, Bartoli B, Cartabia M, Zanetti M, Costantino MA, Canevini MP, et al. Comorbidity prevalence and treatment outcome in children and adolescents with ADHD. Eur Child Adolesc Psychiatry. 2017.

60. Costafreda S. Pooling fMRI data: meta-analysis, mega-analysis and multi-center studies. Front Neuroinform. 2009;3(33).

61. Alexander L, Farrelly N. Attending to adult ADHD: a review of the neurobiology behind adult ADHD. Iri J Psychol Medicine. 2018;35(3):237–44.

