## Supplementary Information for "Analysis of structural brain asymmetries in Attention-Deficit/Hyperactivity Disorder in 39 datasets"

**Table S21** Associations of subcortical volume AIs with comorbidities in ADHD individuals, all age groups combined. 37

**Table S22**. Associations of cortical surface area AIs with comorbidities in ADHD individuals, all age groups combined. 38

**Table S25**. Associations of cortical surface area AIs with psychostimulant medication use in all age groups combined. 38

**Table S23**. Associations of cortical thickness AIs with comorbidities in ADHD individuals, all age groups combined. 40

**Table S24**. Associations of subcortical volume AIs with disorder severity in ADHD individuals, all age groups combined. 42

**Table S25**. Associations of cortical surface area AIs with disorder severity in ADHD individuals, all age groups combined. 43

**Table S26**. Associations of cortical thickness AIs with disorder severity in ADHD individuals, all age groups combined. 44

**Table S27**. Associations of subcortical volume AIs with psychostimulant medication use in ADHD individuals, all age groups combined. 45

**Table S28**. Associations of cortical surface area AIs with psychostimulant medication use in ADHD individuals, all age groups combined. 46

**Table S29**. Associations of cortical thickness AIs with psychostimulant medication use in ADHD individuals, all age groups combined. 47

**Figure S2**. Correlations between AIs of subcortical volumes in the total study sample. 49

**Figure S3**. Correlations between AIs of cortical surface areas in the total study sample. 50

**Figure S4**. Correlations between AIs of cortical thicknes in the total study sample. 51

**Figure S5**. Distributions of age, sex and IQ in ADHD and controls, and of ADHD severity, psychostimulant medication use, and comorbidity in ADHD-only 52

**Figure S6.** Bar plots of the Cohen's *d* effect sizes for diagnosis in the different age groups analyzed. 53

**Figure S2**. Scatter plots of the relationship between age and AIs of the subcortical volumes. 54

**Figure S3**. Scatter plots of the relationship between age and AIs of the cortical surface areas. 55

**Figure S4**. Scatter plots of the relationship between age and AIs of the cortical thickness. 56

**Figure S5**. Residual plots of the linear mixed effects model analysis of subcortical volume AIs in the total study sample. The ggplot2 package in R was used to visualize residuals. 57

**Figure S6**. Residual plots of the linear mixed effects model analysis of cortical surface area AIs and the AI of the total average surface area (totalsurf) in the total study sample. 58

**Figure S7**. Residual plots of the linear mixed effects model analysis of cortical thickness AIs and the AI of the total average thickness (totalthick) in the total study sample. 59

Supplementary References 60

### Complete list of authors and affiliations

Merel C. Postema, MSc[1], Martine Hoogman, PhD[2,3], Sara Ambrosino, MD[4], Philip Asherson, PhD[5], Tobias Banaschewski, MD, PhD[6], Cibele E. Bandeira, MSc[7,8], Alexandr Baranov, PhD[9], Claiton H.D. Bau, PhD[7,8,10], Sarah Baumeister, PhD[6], Ramona Baur-Streubel, PhD[11], Mark A. Bellgrove, PhD[12], Joseph Biederman, MD[13,14], Janita Bralten, PhD[2,3], Daniel Brandeis, PhD[15,16], Silvia Brem, PhD[16,17], Jan K. Buitelaar, MD, PhD[18,19], Geraldo F. Busatto, PhD[20], Francisco X. Castellanos, MD[21,22], Mara Cercignani, PhD[23], Tiffany M. Chaim-Avancini, PhD[20], Kaylita C. Chantiluke, PhD[24], Anastasia Christakou, PhD[24,25], David Coghill, MD[26,27], Annette Conzelmann, PhD[28,29], Ana I. Cubillo, PhD[24], Renata B. Cupertino, PhD[7,8], Patrick de Zeeuw, PhD[30], Alysa E. Doyle, PhD[14,31], Sarah Durston, PhD[30], Eric A. Earl, BSc[32], Jeffery N. Epstein, PhD[33,34], Thomas Ethofer, PhD[35], Damien A. Fair, PhD[32], Andreas J. Fallgatter, MD[36,37], Stephen V. Faraone , PhD[38], Thomas Frodl, MD, PhD[39,40], Matt C. Gabel, PhD[23], Tinatin Gogberashvili, PhD[41], Eugenio H. Grevet, PhD[7,8,10], Jan Haavik, MD, PhD[42,43], Neil A. Harrison, MBBS PhD[23,44], Catharina A. Hartman, PhD[45], Dirk J. Heslenfeld, PhD[46], Pieter J. Hoekstra, MD, PhD[47], Sarah Hohmann, MD[6], Marie F. Høvik, MD[43,48], Terry L. Jernigan, PhD[49], Bernd Kardatzki, Dipl.Phys.[50], Georgii Karkashadze, PhD[9], Clare Kelly, PhD[51,52], Gregor Kohls, PhD[53], Kerstin Konrad, PhD[53,54], Jonna Kuntsi, PhD[5], Luisa Lazaro, MD, PhD[55,56], Sara Lera-Miguel, PhD[57], Klaus-Peter Lesch, MD, PhD[58,59,60], Mario R. Louza, MD, PhD[61], Astri J. Lundervold, PhD[42,62], Charles B Malpas, PhD[63,64], Paulo Mattos, MD, PhD[65,66], Hazel McCarthy, PhD[40,67], Leyla Namazova-Baranova, PhD[9,68], Rosa Nicolau, BSc[69], Joel T Nigg, PhD[32,70], Stephanie E. Novotny, MSc[71], Eileen Oberwelland Weiss, PhD[72,73], Ruth L. O'Gorman Tuura, PhD[74,75], Jaap Oosterlaan, PhD[76,77], Bob Oranje, PhD [30], Yannis Paloyelis, PhD[78], Paul Pauli, PhD[79], Felipe A. Picon, PhD[7], Kerstin J. Plessen, MD, PhD[80,81], J. Antoni Ramos-Quiroga, MD, PhD[82,83,84,85], Andreas Reif, MD[86], Liesbeth Reneman, MD, PhD[87], Pedro G.P. Rosa, MD[20], Katya Rubia, PhD[24], Anouk Schrantee, PhD[88], Lizanne J.S. Schweren, PhD[45], Jochen Seitz, MD[89], Philip Shaw, MD, PhD[90], Tim J. Silk, PhD[91,92], Norbert Skokauskas, MD, PhD[93,94], Juan Carlos Soliva Vila, PhD[95], Michael C. Stevens, PhD[71,96], Gustavo Sudre, PhD[97], Leanne Tamm, PhD[98,99], Fernanda Tovar-Moll, MD, PhD[65,100], Theo G.M. van Erp, PhD[101,102], Alasdair Vance, MD[103], Oscar Vilarroya, PhD[95,104], Yolanda Vives-Gilabert, PhD[105], Georg G. von Polier, MD[89,106], Susanne Walitza, MD[17], Yuliya N. Yoncheva, PhD[107], Marcus V. Zanetti, PhD[108,109], Georg C. Ziegler, MD[58], David C. Glahn, PhD[71,110], Neda Jahanshad, PhD[111], Sarah E. Medland, PhD[112], Paul M. Thompson, PhD[113], Simon E. Fisher, D.Phil[1,3], Barbara Franke, PhD[2,3,114], Clyde Francks, D.Phil[1,3]

| nr | aff |
|---|---|
| 1 | Language and Genetics Department, Max Planck Institute for Psycholinguistics, Nijmegen, The Netherlands |
| 2 | Department of Human Genetics, Radboud university medical center, Nijmegen, Netherlands. |
| 3 | Donders Institute for Brain, Cognition and Behaviour, Radboud University, Nijmegen, Netherlands. |
| 4 | NICHE lab, Department of Psychiatry, University Medical Center Utrecht Brain Center, Utrecht University, Utrecht, The Netherlands |
| 5 | Social, Genetic and Developmental Psychiatry Centre; Institute of Psychiatry, Psychology and Neuroscience, King's College London, London, UK |
| 6 | Department of Child and Adolescent Psychiatry and Psychotherapy, Central Institute of Mental Health, Mannheim, Medical Faculty Mannheim / Heidelberg University, Mannheim, Germany |
| 7 | Adulthood ADHD Outpatient Program (ProDAH), Clinical Research Center, Hospital de Clínicas de Porto Alegre, Porto Alegre, Brazil |

| nr | aff |
|---|---|
| 8 | Department of Genetics, Institute of Biosciences, Universidade Federal do Rio Grande do Sul, Porto Alegre, Brazil |
| 9 | Research Institute of Pediatrics and child health of Central clinical hospital of the Russian Academy of Sciences of the Ministry of Science and Higher Education of the Russian Federation, Moscow, Russia |
| 10 | Developmental Psychiatry Program, Experimental Research Center, Hospital de Clínicas de Porto Alegre, Porto Alegre, Brazil |
| 11 | Department of Biological Psychology, Clinical Psychology and Psychotherapy, University of Würzburg, Würzburg, Germany |
| 12 | Turner Institute for Brain and Mental Health and School of Psychological Sciences, Monash University, Melbourne, Australia |
| 13 | Clinical and Research Programs in Pediatric Psychopharmacology and Adult ADHD |
| 14 | Department of Psychiatry, Massachusetts General Hospital, Harvard Medical School, USA |
| 15 | Department of Child and Adolescent Psychiatry and Psychotherapy, Psychiatric Hospital, University of Zurich, Zurich, Switzerland |
| 16 | The Neuroscience Center Zurich, University of Zurich and ETH Zurich, Zurich, Switzerland |
| 17 | Department of Child and Adolescent Psychiatry and Psychotherapy, Psychiatric Hospital, University of Zurich, Zurich, Switzerland |
| 18 | Department of Cognitive Neuroscience, Donders Institute for Brain, Cognition and Behaviour, Radboudumc, Nijmegen, The Netherlands |
| 19 | Karakter child and adolescent psychiatry University Center, Nijmegen, The Netherlands |
| 20 | Laboratory of Psychiatric Neuroimaging (LIM-21), Department and Institute of Psychiatry, Hospital das Clinicas HCFMUSP, Faculty of Medicine, University of São Paulo, Sao Paulo, Sao Paulo, Brazil |
| 21 | Department of Child and Adolescent Psychiatry, NYU Grossman School of Medicine, New York, NY, USA |
| 22 | Nathan Kline Institute for Psychiatric Research, Orangeburg, NY, USA |
| 23 | Department of Neuroscience, Brighton and Sussex Medical School, Falmer, Brighton, UK |
| 24 | Department of Child and Adolescent Psychiatry, Institute of Psychiatry, Psychology and Neuroscience, King's College London, London, UK |
| 25 | School of Psychology and Clinical Language Sciences, Centre for Integrative Neuroscience and Neurodynamics, University of Reading, Reading, UK |
| 26 | Departments of Paediatrics and Psychiatry, University of Melbourne, Melbourne, Australia |
| 27 | Murdoch Children's Research Institute, Melbourne, Australia |
| 28 | Department of Child and Adolescent Psychiatry, Psychosomatics and Psychotherapy, University Hospital of Tübingen, Germany |
| 29 | PFH – Private University of Applied Sciences, Department of Psychology (Clinical Psychology II), Göttingen, Germany |
| 30 | NICHE Lab, Department of Psychiatry, Brain Center Rudolf Magnus, University Medical Center Utrecht, The Netherlands |
| 31 | Center for Genomic Medicine, Massachusetts General Hospital, Harvard Medical School, USA |
| 32 | Department of Behavioral Neuroscience, Oregon Health & Science University, Portland OR, USA |
| 33 | Division of Behavioral Medicine and Clinical Psychology, Cincinnati Children’s Hospital Medical Center, Cincinnati, OH, USA |
| 34 | Department of Pediatrics, University of Cincinnati College of Medicine, Cincinnati, OH |
| 35 | Clinic for Psychiatry/Psychotherapy Tübingen / Department for Biomedical Magnetic Resonance, Tübingen |
| 36 | Department of Psychiatry and Psychotherapy, University Hospital of Tuebingen, Tuebingen, Germany |
| 37 | LEAD Graduate School, University of Tuebingen, Germany |
| 38 | Departments of Psychiatry and of Neuroscience and Physiology, SUNY Upstate Medical University, Syracuse, New York |
| 39 | Department of Psychiatry and Psychotherapy, Otto von Guericke University Magdeburg, Germany |
| 40 | Department of Psychiatry, Trinity College Dublin, Ireland |
| 41 | National Medical Research Center for Children's Health, Laboratory of Neurology and Cognitive Health, Moscow, Russia |
| 42 | K.G. Jebsen Centre for Neuropsychiatric Disorders, Department of Biomedicine, University of Bergen, Bergen, Norway |
| 43 | Division of Psychiatry, Haukeland University Hospital, Bergen, Norway |
| 44 | Sussex Partnership NHS Foundation Trust, Swandean, East Sussex, UK |
| 45 | University of Groningen, University Medical Center Groningen, Department of Psychiatry, Interdisciplinary Center Psychopathology and Emotion Regulation (ICPE), Groningen, The Netherlands |

| nr | aff |
|---|---|
| 46 | Faculty of Behavioural and Movement Sciences, Vrije Universiteit Amsterdam, Amsterdam, The Netherlands |
| 47 | University of Groningen, University Medical Center Groningen, Department of Child and Adolescent Psychiatry |
| 48 | Department of Clinical Medicine, University of Bergen, Bergen, Norway |
| 49 | Center for Human Development, UC San Diego, USA |
| 50 | Department of Biomedical Magnetic Resonance, University of Tuebingen, Tuebingen, Germany |
| 51 | School of Psychology and Department of Psychiatry at the School of Medicine, Trinity College Dublin, Ireland |
| 52 | Trinity College Institute of Neuroscience, Trinity College Dublin, Ireland |
| 53 | Child Neuropsychology Section, Department of Child and Adolescent Psychiatry, Psychosomatics, and Psychotherapy, University Hospital RWTH Aachen, Germany |
| 54 | JARA Institute Molecular Neuroscience and Neuroimaging (INM-11), Institute for Neuroscience and Medicine, Research Center Jülich, Germany |
| 55 | Institut d'Investigacions Biomèdiques August Pi i Sunyer (IDIBAPS), Barcelona, Spain; Biomedical Network Research Center on Mental Health (CIBERSAM), Barcelona, Spain |
| 56 | Department of Medicine, University of Barcelona, Spain |
| 57 | Department of Child and Adolescent Psychiatry and Psychology, Institute of Neurosciencies, Hospital Clínic, Barcelona |
| 58 | Division of Molecular Psychiatry, Center of Mental Health, University of Würzburg, Würzburg, Germany |
| 59 | Laboratory of Psychiatric Neurobiology, Institute of Molecular Medicine, I.M. Sechenov First Moscow State Medical University, Moscow, Russia |
| 60 | Department of Psychiatry and Neuropsychology, School for Mental Health and Neuroscience (MHeNS), Maastricht University, Maastricht, The Netherlands |
| 61 | Institute of Psychiatry, Faculty of Medicine, University of São Paulo, São Paulo, Brazil |
| 62 | Department of Biological and Medical Psychology, University of Bergen, Bergen, Norway |
| 63 | Developmental Imaging Group, Murdoch Children's Research Institute, Melbourne, Australia |
| 64 | Clinical Outcomes Research Unit (CORe), Department of Medicine, Royal Melbourne Hospital, The University of Melbourne, Melbourne, Australia |
| 65 | D'Or Institute for Research and Education, Rio de Janeiro, Brazil |
| 66 | Federal University of Rio de Janeiro |
| 67 | Centre of Advanced Medical Imaging, St James's Hospital, Dublin, Ireland |
| 68 | Russian National Research Medical University Ministry of Health of the Russian Federation, Moscow, Russia |
| 69 | Department of Child and Adolescent Psychiatry and Psychology, Institut of Neurosciencies, Hospital Clínic, Barcelona, Spain |
| 70 | Department of Psychiatry, Oregon Health & Science University, Portland OR, USA |
| 71 | Olin Neuropsychiatry Research Center, Hartford Hospital, Hartford, CT, USA |
| 72 | Translational Neuroscience, Child and Adolescent Psychiatry, University Hospital RWTH Aachen, Aachen, Germany |
| 73 | Cognitive Neuroscience (INM-3), Institute for Neuroscience and Medicine, Research Center Jülich |
| 74 | Center for MR Research, University Children's Hospital, Zurich, Switzerland |
| 75 | Zurich Center for Integrative Human Physiology (ZIHP) |
| 76 | Clinical Neuropsychology Section, Vrije Universiteit Amsterdam, Amsterdam, the Netherlands |
| 77 | Emma Children's Hospital Amsterdam University Medical Centers, University of Amsterdam, Emma Neuroscience Group, department of Pediatrics, Amsterdam Reproduction & Development, Amsterdam, The Netherlands |
| 78 | Department of Neuroimaging, Institute of Psychiatry, Psychology and Neuroscience, King's College London, London, UK |
| 79 | Department of Psychology (Biological Psychology, Clinical Psychology and Psychotherapy) and Center of Mental Health, University of Würzburg, Würzburg, Germany |
| 80 | Child and Adolescent Mental Health Centre, Capital Region Copenhagen, Denmark |
| 81 | Division of Child and Adolescent Psychiatry, Department of Psychiatry, University Hospital Lausanne, Switzerland |
| 82 | Department of Psychiatry, Hospital Universitari Vall d'Hebron, Barcelona, Catalonia, Spain |
| 83 | Group of Psychiatry, Mental Health and Addictions, Vall d'Hebron Research Institute (VHIR), Barcelona, Catalonia, Spain |
| 84 | Biomedical Network Research Centre on Mental Health (CIBERSAM), Barcelona, Catalonia, Spain |

| nr | aff |
|---|---|
| 85 | Department of Psychiatry and Legal Medicine, Universitat Autònoma de Barcelona, Barcelona, Catalonia, Spain |
| 86 | Department of Psychiatry, Psychosomatic Medicine and Psychotherapy, University Hospital Frankfurt, Frankfurt, Germany |
| 87 | Amsterdam University Medical Center, Academic Medical Center, Amsterdam, the Netherlands |
| 88 | Department of Radiology and Nuclear Medicine, Amsterdam University Medical Centers, Amsterdam; the Netherlands |
| 89 | Child and Adolescent Psychiatry, University Hospital RWTH Aachen, Aachen, Germany |
| 90 | National Human Genome Research Institute and National Institute of Mental health, Bethesda, MD, USA |
| 91 | Deakin University, School of Psychology, Geelong, Australia |
| 92 | Murdoch Children's Research Institute, Developmental Imaging, Melbourne, Australia |
| 93 | Centre for child and adolescent mental health, NTNU, Norway |
| 94 | Institute of Mental Health, Norwegian University of Science and Technology |
| 95 | Department of Psychiatry and Forensic Medicine, Universitat Autonoma de Barcelona, Spain |
| 96 | Department of Psychiatry, Yale University School of Medicine, USA |
| 97 | National Human Genome Research Institute, Bethesda, MD, USA |
| 98 | Department of Pediatrics, Cincinnati Children's Hospital Medical Center, USA |
| 99 | College of Medicine, University of Cincinnati, USA |
| 100 | Morphological Sciences Program, Federal University of Rio de Janeiro, Rio de Janeiro |
| 101 | Clinical Translational Neuroscience Laboratory, Department of Psychiatry and Human Behavior, University of California Irvine, 5251 California Ave, Irvine, CA, 92617, USA |
| 102 | Center for the Neurobiology of Learning and Memory, University of California Irvine, 309 Qureshey Research Lab, Irvine, CA, 92697, USA |
| 103 | Department of Paediatrics, University of Melbourne, Australia |
| 104 | Hospital del Mar Medical Research Institute (IMIM), Barcelona, Spain |
| 105 | Instituto ITACA, Universitat Politècnica de València, València, Spain |
| 106 | Brain and Behavior (INM-7), Institute for Neuroscience and Medicine, Research Center Jülich, Germany |
| 107 | Department of Child and Adolescent Psychiatry, NYU Child Study Center, Hassenfeld Children's Hospital at NYU Langone |
| 108 | Department of Psychiatry, Faculty of Medicine, University of São Paulo, São Paulo, Brazil |
| 109 | Hospital Sírio-Libanês, São Paulo Brazil |
| 110 | Department of Psychiatry, Boston Children's Hospital and Harvard Medical School, Boston, MA 02115-5724, USA |
| 111 | Imaging Genetics Center, Stevens Neuroimaging and Informatics Institute, Keck School of Medicine of USC, Marina del Rey, CA, 90292 |
| 112 | Psychiatric Genetics, QIMR Berghofer Medical Research Institute, Brisbane, Australia |
| 113 | Imaging Genetics Center, Stevens Institute for Neuroimaging & Informatics, Keck School of Medicine, University of Southern California, Los Angeles, CA, USA |
| 114 | Department of Psychiatry, Radboud university medical center, Nijmegen, Netherlands |

#### Acknowledgments

Data were made available for this study by participants of the ENIGMA-ADHD working group (http://enigma.ini.usc.edu/ongoing/enigma-adhd-working-group/). This research was funded by the Max Planck Society (Germany), and many others: ENIGMA: received funding from the National Institutes of Health (NIH) Consortium grant U54 EB020403, supported by a cross-NIH alliance that funds Big Data to Knowledge Centers of Excellence (BD2K). We also are supported by the European College for Neuropsychopharmacology (ECNP) by a grant for the ECNP Network ADHD across the lifespan. SEM was supported by NHMRC grants APP110362, APP1172917 and APP1158127. ADHD-WUE: Data collection and analysis was supported by the Deutsche Forschungsgemeinschaft (KFO 125, TRR 58/A1 and A5, SFB-TRR 58/B01, B06 and Z02, RE1632/5-1) and the research leading to these results also received funding from the 5-100 Russian Academic Excellence Project, European Union's Seventh Framework Programme for research, technological development and demonstration under grant agreement no 602805 (Aggressotype) and the Horizon 2020 research and innovation programme under Grant no. 728018 (Eat2beNICE). ADHD-DUB1 and DUB2: The ADHD-DUB1 and DUB2 studies received funding from the Health Research Board Ireland. ADHD-Mattos: Ivanei Bramati, Paulo Mattos and Fernanda Tovar-Moll were supported by an IDOR intramural grant. ADHD200-KKI: We would like to acknowledge Lindsey Koenig, Michelle Talley, Jessica Foster, Deana Crocetti, Lindsey MacNeil, Andrew Gaddis, Marin Ranta, Anita Barber, Mary Beth Nebel, John Muschelli, Suresh Joel, Brian Caffo, Jim Pekar, Stacy Suskauer. Research was made possible due to the following funding sources: The Autism Speaks Foundation and NIH (R01 NS048527, R01MH078160 and R01MH085328), Johns Hopkins General Clinical Research Center (M01 RR00052), National Center for Resource (P41 RR15241), Intellectual and Developmental Disabilities Research Center (HD-24061) ADHD200-NYU: We would like to acknowledge Amy Roy, Andrea McLaughlin, Ariel Schvarcz, Camille Chabernaud, Chiara Fontani, Christine Cox, Daniel Margulies, David Anderson, David Gutman, Devika Jutagir, Douglas Slaughter, Dylan Gee, Emily Brady, Jessica Raithel, Jessica Sunshine, Jonathan Adelstein, Kristin Gotimer, Leila Sadeghi, Lucina Uddin, Maki Koyama, Natan Potler, Nicoletta Adamo, Rebecca Grzadzinski, Rebecca Lange, Samantha Adelsberg, Samuele Cortese, Saroja Bangaru, Xinian Zuo, Zarrar Shehzad and Zoe Hyde. Data collection was made possible thanks to funding from NIMH (R01MH083246), Autism Speaks, The Stavros Niarchos Foundation, The Leon Levy Foundation, and an endowment provided by Phyllis Green and Randolph Cōwen. ADHD200-Peking: we would like to acknowledge Jue-jing Ren, De-yi Wang, Su-fang Li, Zu-lai Peng, Peng Wang, Yun-yun Zhu, Zhao Qing. Research was made possible due to the following funding sources: The Commonwealth Sciences Foundation, Ministry of Health, China (200802073), The National Foundation, Ministry of Science and Technology, China (2007BAI17B03), The National Natural Sciences Foundation, China (30970802), The Funds for International Cooperation of the National Natural Science Foundation of China (81020108022), The National Natural Science Foundation of China (8100059), Open Research Fund of the State Key Laboratory of Cognitive Neuroscience and Learning Page 4 of 55 ADHD200-OHSU: We would like to acknowledge the Advanced Imaging Research Center, Bill Rooney, Kathryn L. Mills, Taciana G. Costa Dias, Michelle C. Fenesy, Bria L. Thurlow, Corrine A. Stevens, Samuel D. Carpenter, Michael S. Blythe, Colleen F. Schmitt. Research was made possible due to the following funding resources: K99/R00 MH091238 (Fair), R01 MH086654 (Nigg), Oregon Clinical and Translational Research Institute (Fair), Medical Research Foundation (Fair), UNCF/Merck

(Fair), Ford Foundation (Fair) ADHD-UKA: KFO-112 and IRTG1328 was supported by the German Research Foundation (DFG). DAT-London: This work was supported in part by UK Medical Research, Council Grant G03001896 to J Kuntsi and NIH grants, R01MH62873 and R01MH081803 to SV Faraone. IMpACT: The IMpACT study was supported by a grant from the Brain & Cognition Excellence Program and a personal Vici grant (to Barbara Franke) of the Netherlands Organization for Scientific Research (NWO, grant numbers 433-09-229 and 016-130-669) and in part by the Netherlands Brain Foundation (grant number, 15F07[2]27) and the BBMRI-NL (grant CP2010-33). Funding was also provided by a pilot grant of the Dutch National Research Agenda for the NeuroLabNL project (grant 400 17 602). The research leading to these results also received funding from the European Community's Seventh Framework Programme (FP7/2007–2013) under grant agreement no. 602805 (Aggressotype), no. 278948 (TACTICS), and no. 602450 (IMAGEMEND). In addition, the project received funding from the European Union's Horizon 2020 research and innovation programme under the Marie Sklodowska-Curie grant agreement no. 643051 (MiND), under grant agreement no. 667302 (CoCA) and the grant agreement no. 728018 (Eat2beNICE). Niche: The structural neuroimaging studies of NICHE were supported by VIDI and VICI grants from the Netherlands Organization for Scientific Research (Nederlandse Organisatie voor Wetenschappelijk Onderzoek, NWO) to Sarah Durston (grant numbers Vidi-91.776.384 and Vici-453-10-005). NYU ADHD: NYU data collection and sharing was supported by NIH grants T32MH67763, R01MH083246, K23MH087770, R01MH094639, and U01MH099059 and a grant from the Stavros S. Niarchos Foundation. UAB-ADHD: The study and its contributors received funding from the Ministerio de Economía y Competitividad under research grant SAF2012-32362 and: PI12/01139 from the Department of Health of the Government of Catalonia. Additional funding was obtained from the Generalitat de Catalunya. ZI-CAPS: The Neurofeedback study was partly funded by the project D8 of the Deutsche Forschungsgesellschaft collaborative research center 636. ADHD-Rubia: The study was funded by the UK Department of Health via the National Institutes of Health Research Centre (BRC) for Mental Health South London and the Maudsley NHS Foundation Trust and the Institute of Psychiatry, King's College London. CAPS_UZH: The data contributed to this study were collected in two projects on ADHD and OCD in children and adolescents, supported by the Swiss National Science Foundation (projects No. 136249 Sinergia and No. 320030_130237) and the Hartmann Müller Foundation (No. 1460). Page 5 of 55 NeuroIMAGE: This work was supported by NIH Grant R01MH62873, NWO Large Investment Grant 1750102007010 and grants from Radboud University Medical Center, University Medical Center Groningen and Accare, and VU University Amsterdam. This work was also supported by grants from NWO Brain & Cognition (433-09-242 and 056-13-015) and from ZonMW (60-60600-97-193). Further support was received from the European Union FP7 programmes TACTICS (278948), IMAGEMEND (602450), Horizon2020 programmes CANDY (847818) and IMI programme EU-AIMS (115300) and AIMS-2-TRIALS (777394). MTA: Data collection and sharing for this project was funded by the NIDA MTA Neuroimaging Study (National Institute on Drug Abuse Grant Contract #: HHSN271200800009C). NIH: studies were supported by intramural grants at the National Institute of Mental Health and National Human Genome Research Institute. OHSU: The OHSU work was supported by NIMH grants R01MH86654, MH099064, and MH115357. UCHZ: This work was supported by the University Research Priority Program "Integrative Human Physiology" at the University of Zurich. ACPU: This research was conducted within the Academic Child Psychiatry Unit, University of Melbourne, Royal Children's Hospital and the Developmental Imaging research group, Murdoch Children's Research Institute, Melbourne,

Victoria. National Health and Medical Research Council of Australia (NHMRC) project grants 384419 and 569533 provided funds for the data collection. It was also supported by the Murdoch Children's Research Institute, the Royal Children's Hospital and the Children's MRI Centre, The Royal Children's Hospital Foundation, and the RCH Mental Health Service, Department of Paediatrics The University of Melbourne and the Victorian Government's Operational Infrastructure Support Program. Tim Silk was supported by an NHMRC Career Development Award. NICAP: The Neuroimaging of the Children's Attention Project was funded by the National Medical Health and Research Council of Australia (NHMRC; project grant #1065895). Earlier funding of the Children's Attention Project was funded by an NHMRC project grant #1008522 and a grant from the Collier Foundation. This research was conducted within the Developmental Imaging research group, Murdoch Children's Research Institute and the Children's MRI Centre, The Royal Children's Hospital, Melbourne, Victoria. It was supported by the Murdoch Children's Research Institute, The Royal Children's Hospital, The Royal Children's Hospital Foundation, Department of Paediatrics at The University of Melbourne and the Victorian Government's Operational Infrastructure Support Program. Tübingen: The recruitment of the Tübingen sample was funded by the Deutsche Forschungsgemeinschaft (DFG grant: ET 112/5-1). Dundee: This work was supported by a grant from TENOVUS SCOTLAND and was conducted in collaboration with the Dundee site of the ADHD Drugs Use Chronic Effects (ADDUCE) study (EU FP7 agreement No. 260576). ePOD: The neuroimaging studies of the ePOD-MPH trial (NTR3103) were supported by faculty resources of the Academic Medical Center, University of Amsterdam, and by grant 11.32050.26 from the European Page 6 of 55 Research Area Network Priority Medicines for Children (Sixth Framework Programme) to Liesbeth Reneman. Sao Paulo: The present investigation was supported by grants from the Brain & Behavior Research Foundation (2010 NARSAD Independent Investigator Award granted to Geraldo F. Busatto), FAPESP-Brazil (2013/03905-4) and CNPq-Brazil (#478466/2009 & 480370/2009). Sussex: This study was supported by funding from Brighton and Sussex Medical School and the Dr. Mortimer and Dame Theresa Sackler Foundation. Clinic Barcelona: This work has received financial support from two grants, Fundació la Marató de TV3- 2009 (project number: 091810) and Fondo de Investigaciones Sanitarias, of the Spanish Ministry of Health (project number: PI11/01419). Generation R: Supercomputing resources were supported by the NWO Physical Sciences Division (Exacte Wetenschappen) and SURFsara (Cartesius compute cluster, www.surfsara.nl). The neuroimaging and neuroimaging infrastructure was supported by the Netherlands Organization for Health Research and Development (ZonMw) TOP project number 91211021 to TW. The Generation R Study is conducted by the Erasmus Medical Center in close collaboration with Faculty of Social Sciences of the Erasmus University Rotterdam, the Municipal Health Service Rotterdam area, Rotterdam, and the Stichting Trombosedienst & Artsenlaboratorium Rijnmond (STAR-MDC), Rotterdam. We gratefully acknowledge the contribution of children and parents, general practitioners, hospitals, midwives and pharmacies in Rotterdam. The general design of Generation R Study is made possible by financial support from the Erasmus Medical Center, Rotterdam, the Erasmus University Rotterdam, ZonMw, the Netherlands Organisation for Scientific Research (NWO), and the Ministry of Health, Welfare and Sport. Martine Hoogman: supported by a personal Veni grant from of the Netherlands Organization for Scientific Research (NWO, grant number 91619115) Maarten Mennes: supported by a Marie Curie International Incoming Fellowship within the 7th European Community Framework Programme, grant agreement n° 327340. Ryan Muetzel: supported by Friends of Sophia Foundation project S18-20. Jan Haavik: K.G. Jebsen Centre for Research on

Neuropsychiatric Disorders, University of Bergen, Bergen, Norway ZI-CAPS: We would like to acknowledge Isabella Wolf, Nathalie Holz and Regina Boecker-Schlier. CAPS_UZH: We would like to acknowledge Tobias Hauser, Anthony Schläpfer and Reto Iannaccone. MTA: The Multimodal Treatment Study of Children with ADHD (MTA) was a National Institute of Mental Health (NIMH) cooperative agreement randomized clinical trial, continued under an NIMH contract as a follow-up study and finally under a National Institute on Drug Abuse (NIDA) contract. Collaborators from NIMH: Benedetto Vitiello, M.D. (Child & Adolescent Treatment and Preventive Interventions Research Branch), Joanne B. Severe, M.S. (Clinical Trials Operations and Biostatistics Unit, Division of Services and Intervention Research), Peter S. Jensen, M.D. (currently at REACH Institute and Mayo Clinic), L. Eugene Arnold, M.D., M.Ed. (currently at Ohio State University), Kimberly Hoagwood, Ph.D. (currently at New York University); previous contributors from NIMH to the early phases: John Richters, Ph.D. (currently at National Institute of Nursing Research); Donald Vereen, M.D. (currently at NIDA). Principal investigators and co-investigators from the sites are: University of California, Berkeley/San Francisco: Stephen P. Hinshaw, Ph.D. (Berkeley), Glen R. Elliott, Ph.D., M.D. (San Francisco); Duke University Medical Center: Karen C. Wells, Ph.D., Jeffery N. Epstein, Ph.D. (currently at Cincinnati Children's Hospital Medical Center), Desiree W. Murray, Ph.D.; previous Duke contributors to early phases: C. Keith Conners, Ph.D. (former PI); John March, M.D., M.P.H.; University of California, Irvine: James Swanson, Ph.D., Timothy Wigal, Ph.D.; previous contributor from UCLA to the early phases: Dennis P. Cantwell, M.D. (deceased); New York University: Howard B. Abikoff, Ph.D.; Montreal Children's Hospital/ McGill University: Lily Hechtman, M.D.; New York State Psychiatric Institute/Columbia University/Mount Sinai Medical Center: Laurence L. Greenhill, M.D. (Columbia), Jeffrey H. Newcorn, M.D. (Mount Sinai School of Medicine). University of Pittsburgh: Brooke Molina, Ph.D., Betsy Hoza, Ph.D. (currently at University of Vermont), William E. Pelham, Ph.D. (PI for early phases, currently at Florida International University). Follow-up phase statistical collaborators: Robert D. Gibbons, Ph.D. (University of Illinois, Chicago); Sue Marcus, Ph.D. (Mt. Sinai College of Medicine); Kwan Hur, Ph.D. (University of Illinois, Chicago). Original study statistical and design consultant: Helena C. Kraemer, Ph.D. (Stanford University). Collaborator from the Office of Special Education Programs/US Department of Education: Thomas Hanley, Ed.D. Collaborator from Office of Juvenile Justice and Delinquency Prevention/Department of Justice: Karen Stern, Ph.D. Additional investigators for Neuroimaging Substudy: Leanne Tamm, Ph.D., PI (Cincinnati Children's Hospital Medical Center), James Bjork, Ph.D. (Division of Clinical Neuroscience and Behavioral Research, NIDA; currently at Virginia Commonwealth University), Daniel Mathalon, M.D., Ph.D. (UC San Francisco), Allen Song, Ph.D. (Duke), Bradley Peterson, M.D. (currently USC), Steven Potkin, M.D. & Claudia Buss, Ph.D. (UC Irvine), Katerina Velanova, Ph.D. (Pittsburgh), Neuroimaging Consultants: Susan Tapert, Ph.D. & Joshua Kuperman, Ph.D. (UC San Diego), BJ Casey, Ph.D. & Leah Sommerville, Ph.D. (Sackler Institute, Cornell, currently at Yale and Harvard, respectively), Krista Lisdahl, Ph.D. (University of WisconsinMilwaukee). Neuroimaging Analysis and Interpretation: Terry Jernigan, Ph.D. & Anders Dale, Ph.D. (UC San Diego), F. Xavier Castellanos, M.D. & Clare Kelly, Ph.D. (New York University). UCHZ: We would like to acknowledge Carmen Ghisleni, Steffen Bollmann, Lars Michels, Peter Klaver, Simon Shlomo Poil, Stefanie Kübel, Juliane Ball, Dominique Eich-Höchli, and Ernst Martin. ADHD_Russia: We would like to acknowledge Vladimir Zelman, Boris Gutman, Anait Gevorkyan, Vladimir Smirnov NICAP: We would like to acknowledge other investigators: Emma Sciberras, Daryl Efron, Vicki Anderson, Jan M. Nicholson, Philip Hazell, and all the staff and students of the Children's Attention Project, as well as the RCH Medical Imaging staff for their assistance and expertise in the collection of the MRI data included in this study. We would also like to thank Ivanei Bramati, PhD, Anna Calvo, MSc, Anders Dale, PhD, and Lena Schwarz, MD, for their contributions to the study, as well as all of the many families and schools for their participation in this study.

#### Disclosures

Mr. Earl is co-inventor of the Oregon Health and Science University Technology #2198 (co-owned with Washington University in St. Louis), FIRMM: Real time monitoring and prediction of motion in MRI scans, exclusively licensed to Nous, Inc.) and any related research. Any potential conflict of interest has been reviewed and managed by OHSU. Dr. Biederman has received research support from AACAP, Alcobra, the Feinstein Institute for Medical Research, the Forest Research Institute, Genentech, Headspace, Ironshore, Lundbeck AS, Magceutics, Merck, Neurocentria, NIDA, NIH, PamLab, Pfizer, Roche TCRC, Shire, SPRITES, Sunovion, the U.S. Department of Defense, the U.S. Food and Drug Administration, and Vaya Pharma/Enzymotec; he has served as a consultant or on scientific advisory boards for Aevi Genomics, Akili, Alcobra, Arbor Pharmaceuticals, Guidepoint, Ironshore, Jazz Pharma, Medgenics, Piper Jaffray, and Shire; he has received honoraria from Alcobra, the American Professional Society of ADHD and Related Disorders, and the MGH Psychiatry Academy for tuition-funded CME courses; he has a financial interest in Avekshan, a company that develops treatments for ADHD; he has a U.S. patent application pending (Provisional Number #61/233,686) through MGH corporate licensing, on a method to prevent stimulant abuse; and his program has received royalties from a copyrighted rating scale used for ADHD diagnoses, paid to the Department of Psychiatry at Massachusetts General Hospital by Ingenix, Prophase, Shire, Bracket Global, Sunovion, and Theravance. Dr. Van Erp has served as consultant for Roche Pharmaceuticals and has a contract with Otsuka Pharmaceutical, Ltd. Dr. Gabel has received funding from the Motor Neurone Disease Association. Dr. Asherson has served as a consultant and as a speaker at sponsored events for Eli Lilly, Novartis, and Shire, and he has received educational/research awards from Eli Lilly, GW Pharma, Novartis, QbTech, Shire, and Vifor Pharma. Dr. Brandeis has served as an unpaid scientific consultant for an EU-funded neurofeedback trial. Dr. Karkashadze has received payment for article authorship and speaking fees from Sanofi and from Pikfarma. Dr. Mattos has served on speakers' bureau and/or as a consultant for Janssen-Cilag, Novartis, and Shire and has received travel awards from those companies to participate in scientific meetings; the ADHD outpatient program (Grupo de Estudos do Déficit de Atenção/Institute of Psychiatry) chaired by Dr. Mattos also received research support from Novartis and Shire. Dr. Banaschewski served in an advisory or consultancy role for Lundbeck, Medice, Neurim Pharmaceuticals, Oberberg GmbH, Shire, and Infectopharm. He received conference support or speaker's fee from Lilly, Medice, and Shire. He received royalties from Hogrefe, Kohlhammer, CIP Medien, Oxford University Press; the present work is unrelated to these relationships.Dr. Paloyelis has received an unrestricted research grant from PARI GmbH. Dr. Coghill has served in an advisory or consultancy role for Eli Lilly, Medice, Novartis, Oxford Outcomes, Shire, and Viforpharma; he has received conference support or speaking fees from Eli Lilly, Janssen McNeil, Medice, Novartis, Shire, and Sunovion; and he has been involved in clinical trials conducted by Eli Lilly and Shire. Dr. Kuntsi has received speaking honoraria and advisory panel payments for participation at educational events sponsored by Medice; all funds are received by King's College London and used for studies of ADHD. Dr. Mehta has received research funding from Lundbeck, Shire, and Takeda and has served on advisory boards for Lundbeck and Autifony. Dr. Harrison has received research funding from Janssen Pharmaceuticals. Dr. Bellgrove has received speaking fees and travel support from Shire. Dr. Rubia has received a grants from Eli Lilly/Takeda pharmaceuticals for another project. Dr. Walitza has received lecture honoraria from Eli Lilly and Opopharma, support from the Hartmann Müller, Olga Mayenfisch, and Gertrud Thalmann foundations, and royalties from Beltz, Hogrefe, Kohlhammer, Springer, and Thieme. Dr. Haavik has received speaking fees from Biocodex, Eli Lilly, HB Pharma, Janssen-Cilag, Medice, Novartis, and Shire. Dr. Lesch has served as a speaker for Eli Lilly and has received research support from Medice and travel support from Shire. Dr. Reif has received honoraria for serving as speaking or on advisory boards for Janssen, Medice, Neuraxpharm, Servier and Shire. Dr. Konrad has received speaking fees from Eli Lilly, Medice, and Shire. Dr. Hoekstra served on the advisory board for Shire. Dr. Ramos-Quiroga has served on the speakers bureaus and/or as a consultant for Almirall, Braingaze, Eli Lilly, Janssen-Cilag, Lundbeck, Medice, Novartis, Shire, Takeda, Shionogui, Bial, Sincrolab, and Rubió; he has received travel awards for taking part in psychiatric meetings from Eli Lilly, Janssen-Cilag, Medice, Rubió, and Shire, Takeda, Bial, Shionogui; and the Department of Psychiatry chaired by him has received unrestricted educational and research support from Actelion, Eli Lilly, Ferrer, Janssen-Cilag, Lundbeck, Oryzon, Psious, Roche, Rubió, and Shire. Dr. Fair is a founder of Nous Imaging, Inc.; any potential conflicts of interest are being reviewed and managed by OHSU. Dr. Thompson has received funding support from Biogen. Dr. Buitelaar has served as a consultant, advisory board member, and/or speaker for Eli Lilly, JanssenCilag, Medice, Roche, Takeda/Shire, Angelini, and Servier. Dr. Faraone has received income, potential income, travel expenses, continuing education support, and/or research support from Akili Interactive Labs, Alcobra, Arbor, Enzymotec, Genomind, Ironshore, Janssen, KemPharm, McNeil, Neurolifesciences, Neurovance, Novartis, Otsuka, Pfizer, Rhodes, Shire/Takeda, Sunovion, Supernus, Tris, and Medice; he receives royalties from Elsevier, Guilford Press, and Oxford University Press; he is principal investigator of www.adhdinadults.com; and, with his institution, he holds U.S. patent US20130217707 A1 for the use of sodium-hydrogen exchange inhibitors in the treatment of ADHD. Dr. Franke has received educational speaking fees from Shire and Medice.

### Supplementary Methods

#### ENIGMA imaging processing protocols

Visual inspection of both internal and external Freesurfer segmentations was done per site. All sites followed the standardized ENIGMA protocols that are publicly available on http://enigma.ini.usc.edu/protocols/imaging-protocols. In short, outliers were determined by calculating the interquartile range (IQR) for each of the values per cohort and per diagnostic group (ADHD and Controls). Values that were above or below 1.5 times the IQR were identified as an outlier, and were visually inspected (3D) by the researcher. When a segmentation failure was identified, all values from the affected regions were excluded from further analyses. Additionally, cortical segmentations were overlayed on T1 images of the subjects. Webpages were generated with snapshots from internal slices, and also with external views of the segmentation from different angles. All sites were provided with the manual on how to judge these images, including the most common segmentation errors.

#### Cohen's d calculation

The t-statistic for the factor 'diagnosis' in each linear mixed effects model was derived and used to calculate Cohen's $d$, with

$$d = \frac{t * (n1+n2)}{\sqrt{(n1*n2)} * \sqrt{df}}$$

where $n1$ and $n2$ are the number of cases and controls, and $df$ the degrees of freedom. The latter was derived from the lme summary table in R, but can also be calculated using $df = obs - (x1 + x2)$, wherein $obs$ equals the number of observations, $x1$ the number of groups and $x2$ the number of factors in the model.

The 95% confidence intervals for Cohen's $d$ were calculated using 95% $CI = d \pm 1.96 * SE$, with the standard error (SE) around Cohen's d calculated according to:

$$SE = \sqrt{\frac{n1+n2}{n1*n2} + \frac{d^2}{2*(n1+n2-2)}}$$

#### IQ, comorbidity, ADHD severity and psychostimulant medication

IQ was assessed differently per dataset, but most frequently using an age-appropriate version of the Wechsler intelligence scales (**Supplementary Table S1**). Similarly, datasets used different instruments to assess comorbidity, but this was most often assessed by means of the Structural Clinical Interview for DSM-IV Axis I Disorders (SCID) (2), or using the Schedule for Affective Disorders and Schizophrenia for School-Age Children Present and Lifetime Version (KSADS-PL) (3) (**Supplementary Table S1**). ADHD severity was assessed based on the Conners questionnaires (4), and included both hyperactivity/impulsivity and inattention scores, which were tested for associations with brain asymmetries in separate models. The analysis of psychostimulant medication was also done with two separate analyses, firstly testing current use ('currently using stimulants' versus 'not currently using stimulants') and secondly testing lifetime use ('ever used stimulant' versus 'never used stimulants').

**Table S1.** Characteristics of the different datasets used in the mega-analysis

| Sample name | N total | N cases (M/F) | N controls (M/F) | median age (range) | F | FS | Diagnostic instrument | IQ instrument | Comorbidity instrument | Instrument for symptom rating |
|---|---|---|---|---|---|---|---|---|---|---|
| ACPU[2] | 67 | 39/0 | 28/0 | 13(9, 18) | 5.3 | 3 T | DSM-IV | WISC subtests and full | DISC-IV | Conners parent long version |
| Amsterdam Neuroimage | 173 | 68/23 | 54/28 | 17(11, 26) | 5.3 | 1.5 T | DSM-IV | Vocabulary and block design subtest of WAIS/WIC | K-SADS-PL | NA |
| BergenADHD | 81 | 21/17 | 16/27 | 29(21, 48) | 5.3 | 3 T | ICD10 or DSM-IV | WASI | NA | NA |
| CAPSUZH scan1 | 57 | 15/6 | 21/15 | 11(8, 18) | 5.3 | 3 T | ICD10 and DSM-IV | WISC subtests block design, similarities, digit span | K-SADS-PL | NA |
| DATlondon | 56 | 27/0 | 29/0 | 16(12, 21) | 5.3 | 3 T | DSM-IV | Vocabulary and block design subtests of WAIS | NA | NA |
| Dublin1 | 80 | 30/9 | 32/9 | 20.5(18, 49) | 5.3 | 3 T | DSM-IV | Verbal comprehension, perceptual reasoning, working memory and processing speed subtests of WAIS-IV | SCID-I | Conners Adult ADHD rating scale observer |
| Dundee | 45 | 16/6 | 10/13 | 13(10, 18) | 5.3 | 3 T | DSM-IV | British Picture Vocabulary Scale standardised Score (proxy for verbal IQ) mean 100 SD 15 | K-SADS-PL, SNAP IV | NA |
| IMpACT-NL | 274 | 57/80 | 55/82 | 33(18, 63) | 5.3 | 1.5 T | DSM-IV | Vocabulary and block design subtests of WAIS | SCID-I and SCID-II | NA |
| ADHD200KKI[1] | 85 | 14/6 | 39/26 | 10(8, 12) | 5.3 | 1.5 T | DSM-IV | WISC-IV | NA | Conners Parent Rating Scale Revised Long Version |
| Clinic Barcelona | 73 | 52/0 | 21/0 | 11(8, 16) | 5.3 | 3 T | DSM-IV | Cognitive General Index (CGI) from WISC-IV | K-SADS | Conners Parents' Rating Scales |
| MGH | 144 | 41/36 | 28/39 | 35(18, 59) | 5.1 | 1.5 T | DSM-IV | Vocabulary and block design of WAIS | SCID-I | NA |
| MTA scan1 | 18 | 6/4 | 5/3 | 25(22, 27) | 5.3 | 3 T | DSM-IV | WISC-III full version /subtests of WISC-II (administered at baseline in childhood) | NA | NA |
| MTA scan2 | 25 | 12/4 | 7/2 | 25(21, 28) | 5.3 | 3 T | DSM-IV | WISC-III full version /subtests of WISC-II (administered at baseline in childhood) | NA | NA |
| MTA scan3 | 16 | 9/1 | 6/0 | 24(22, 26) | 5.3 | 3 T | DSM-IV | WISC-III full version /subtests of WISC-II | NA | NA |

| Sample name | N total | N cases (M/F) | N controls (M/F) | median age (range) | F | FS | Diagnostic instrument | IQ instrument | Comorbidity instrument | Instrument for symptom rating |
|---|---|---|---|---|---|---|---|---|---|---|
| | | | | | | | | (administered at baseline in childhood) | | |
| MTA scan4 | 22 | 13/3 | 6/0 | 24(22, 26) | 5.3 | 3 T | DSM-IV | WISC-III full version /subtests of WISC-II (administered at baseline in childhood) | NA | NA |
| MTA scan5 | 24 | 16/2 | 2/4 | 24.5(22, 27) | 5.3 | 3 T | DSM-IV | WISC-III full version /subtests of WISC-II (administered at baseline in childhood) | NA | NA |
| MTA scan6 | 24 | 17/1 | 5/1 | 25(22, 30) | 5.3 | 3 T | DSM-IV | WISC-III full version /subtests of WISC-II (administered at baseline in childhood) | NA | NA |
| NICAP | 146 | 53/12 | 47/34 | 10(9, 11) | 5.3 | 3 T | DSM-IV | WASI: vocabulary, matrix reasoning | DISC-IV | NA |
| Niche scan1 | 108 | 49/6 | 44/9 | 11(7, 16) | 5.1 | 1.5 T | DSM-IV | Vocabulary and block design WISC-III | DISC-IV | NA |
| Niche scan2 | 47 | 17/4 | 22/4 | 9(7, 16) | 5.1 | 1.5 T | DSM-IV | Vocabulary and block design WISC-III | DISC-IV | NA |
| NIH | 331 | 111/55 | 110/55 | 11(4, 18) | 5.3 | 1.5 T | DSM-IV | Subtests of WISC | NA | NA |
| Nijmegen Neuroimage | 158 | 82/38 | 23/15 | 18(9, 24) | 5.3 | 1.5 T | DSM-IV | Vocabulary and block design substest of WAIS/WIC | K-SADS-PL | NA |
| NYU | 80 | 22/18 | 22/18 | 30(18, 53) | 5.3 | 3 T | DSM-IV | WASI | SCID-I | NA |
| NYUADHD200[1] | 228 | 94/35 | 48/51 | 11(7, 18) | 5.3 | 3 T | DSM-IV | WASI | NA | Conners Parent Rating Scale Revised Long Version |
| OHSUADHD200[1] | 89 | 19/7 | 28/35 | 9(7, 13) | 5.3 | 3 T | DSM-IV | Block Design, Vocabulary and Information subtests of WISC-IV | NA | Parent/Teacher Conners rating scale 3rd edition |
| OHSU | 229 | 81/39 | 59/50 | 9(7, 13) | 5.3 | 3 T | DSM-IV and DSM-V | WISC subtests: block design, vocabulary, and information | NA | NA |
| Olin Neuropsychiatry Research Centre[2] | 181 | 59/14 | 58/50 | 15(12, 19) | 5.3 | 3 T | DSM-IV | WASI Full Scale | KSADS-PL | NA |
| PekingADHD200 scan1[1] | 32 | 14/0 | 18/0 | 13.5(12, 16) | 5.3 | 3 T | DSM-IV | WISCC-R | NA | NA |
| PekingADHD200 scan2[1] | 64 | 32/0 | 31/1 | 12(9, 16) | 5.3 | 3 T | DSM-IV | WISCC-R | NA | NA |
| PekingADHD200 scan3[1] | 133 | 34/11 | 30/58 | 11(8, 17) | 5.3 | 3 T | DSM-IV | WISCC-R | NA | NA |
| Rubia ADHD | 71 | 41/0 | 30/0 | 14(10, 18) | 5.3 | 3 T | DSM-IV | WASI | Comorbid disorders | NA |

| Sample name | N total | N cases (M/F) | N controls (M/F) | median age (range) | F | FS | Diagnostic instrument | IQ instrument | Comorbidity instrument | Instrument for symptom rating |
|---|---|---|---|---|---|---|---|---|---|---|
| | | | | | | | | | were exclusion criteria | |
| SãoPaulo1 – Estado | 147 | 57/24 | 44/22 | 27(17, 50) | 5.3 | 3 T | DSM-IV | WASI | SCID | NA |
| Sussex[2] | 60 | 19/11 | 19/11 | 31(19, 59) | 5.3 | 1.5 T | DSM-IV | NART | NA | NA |
| SVG Bergen | 51 | 19/4 | 20/8 | 10(8, 12) | 5.3 | 3 T | DSM-IV | WISC-IV | K-SADS-PL | NA |
| UAB | 198 | 82/21 | 64/31 | 27(6, 52) | 5.3 | 3 T | DSM-IV | WISC | NA | NA |
| UCHZ | 78 | 20/19 | 21/18 | 14.5(9, 61) | 5.3 | 3 T | DSM-IV | HAWIK | K-SADS-PL | Adults; Adult Conners; Children Conners -3d |
| UKA scan4 | 59 | 21/7 | 13/18 | 9(4, 15) | 5.3 | 3 T | ICD10 or DSM-IV | CPM (N=30) or WASI (N=49) or WISC-IV (N=14) | K-SADS and German K-Dips | NA |
| Wurzburg ADHD | 107 | 30/25 | 24/28 | 43(18, 62) | 5.3 | 1.5 T | DSM-IV | MWT-B | SCID-I | NA |
| ZiCAPS | 34 | 17/4 | 7/6 | 13(9, 15) | 5.3 | 3 T | DSM-IV | Subscales of WICS-IV | NA | NA |
| **Total** | **4180** | **2246** | **1934** | | | | | | | |

[1] Included in the mega-analysis by Douglas *et al*. (2018), mentioned in the introduction. [2] Only cortical data available for these datasets. F=FreeSurfer version; FS= Field Strength.

**Table S2.** Full linear model results for the subcortical volume AIs in children

| Subcortical volume AI | | beta-coefficient | | | Standard Error | | | t-value | | | p-value | | | Cohen's d (95% CI) |
|---|---|---|---|---|---|---|---|---|---|---|---|---|---|---|
| | N cases/ controls | diag | sex | age | diag | sex | age | diag | sex | age | diag | sex | age | diag |
| Accumbens | 842/928 | -0.0042 | -0.0004 | -0.0007 | 0.0035 | 0.0038 | 0.0009 | -1.22 | -0.10 | -0.78 | 0.224 | 0.924 | 0.435 | -0.06 (-0.15,0.04) |
| Amygdala | 841/928 | -0.0001 | 0.0040 | 0.0010 | 0.0025 | 0.0028 | 0.0007 | -0.04 | 1.42 | 1.47 | 0.964 | 0.156 | 0.141 | -0.002 (-0.1,0.09) |
| Caudate Nucleus | 843/928 | 0.0006 | -0.0033 | 0.0002 | 0.0014 | 0.0016 | 0.0004 | 0.45 | -2.11 | 0.44 | 0.655 | **0.035** | 0.660 | 0.02 (-0.07,0.11) |
| Globus Pallidus | 843/926 | 0.0003 | -0.0005 | -0.0018 | 0.0026 | 0.0029 | 0.0007 | 0.11 | -0.18 | -2.58 | 0.913 | 0.861 | **0.010** | 0.01 (-0.09,0.1) |
| Hippocampus | 841/926 | -0.0007 | 0.0000 | 0.0008 | 0.0020 | 0.0022 | 0.0005 | -0.37 | -0.01 | 1.56 | 0.710 | 0.991 | 0.119 | -0.02 (-0.11,0.08) |
| Putamen | 843/925 | -0.0012 | 0.0025 | 0.0009 | 0.0014 | 0.0015 | 0.0004 | -0.90 | 1.69 | 2.54 | 0.369 | 0.092 | **0.011** | -0.04 (-0.14,0.05) |
| Thalamus[1] | 747/819 | 0.0010 | 0.0011 | 0.0039 | 0.0014 | 0.0016 | 0.0004 | 0.72 | 0.72 | 9.38 | 0.473 | 0.472 | **$2.2\times10^{-20}$** | 0.04 (-0.06,0.14) |

P-values in **bold** are significant at the uncorrected level ($P < 0.05$). [1]Thalamus volume was not available from the NIH dataset.

**Table S3.** Full linear model results for the cortical surface area AIs in children

| Cortical surface area AI | | beta-coefficient | | | Standard Error | | | t-value | | | p-value | | | Cohen's d (95% CI) |
|---|---|---|---|---|---|---|---|---|---|---|---|---|---|---|
| | N cases / controls | diag | sex | age | diag | sex | age | diag | sex | age | diag | sex | age | diag |
| Banks of superior temporal sulcus | 883/962 | -0.0017 | 0.0008 | 0.0032 | 0.0040 | 0.0044 | 0.0011 | -0.42 | 0.19 | 2.91 | 0.677 | 0.849 | **0.004** | -0.02 (-0.11,0.07) |
| Caudal anterior cingulate cortex | 951/1029 | -0.0024 | 0.0062 | -0.0001 | 0.0054 | 0.0059 | 0.0014 | -0.45 | 1.06 | -0.07 | 0.651 | 0.291 | 0.941 | -0.02 (-0.11,0.07) |
| Caudal middle frontal cortex | 953/1034 | 0.0031 | -0.0005 | -0.0010 | 0.0035 | 0.0039 | 0.0009 | 0.90 | -0.13 | -1.10 | 0.368 | 0.899 | 0.273 | 0.04 (-0.05,0.13) |
| Cuneus | 947/1034 | 0.0051 | 0.0036 | 0.0015 | 0.0031 | 0.0034 | 0.0008 | 1.64 | 1.06 | 1.83 | 0.101 | 0.291 | 0.067 | 0.07 (-0.01,0.16) |
| Entorhinal cortex | 910/1001 | -0.0012 | -0.0071 | 0.0037 | 0.0057 | 0.0062 | 0.0015 | -0.21 | -1.14 | 2.56 | 0.835 | 0.254 | **0.011** | -0.01 (-0.1,0.08) |
| Frontal pole | 953/1035 | -0.0075 | 0.0021 | -0.0012 | 0.0042 | 0.0046 | 0.0010 | -1.79 | 0.45 | -1.22 | 0.074 | 0.654 | 0.224 | -0.08 (-0.17,0.01) |
| Fusiform gyrus | 947/1031 | -0.0035 | 0.0022 | 0.0004 | 0.0024 | 0.0026 | 0.0006 | -1.47 | 0.83 | 0.73 | 0.141 | 0.405 | 0.463 | -0.07 (-0.15,0.02) |
| Inferior parietal cortex | 950/1029 | 0.0035 | 0.0116 | -0.0005 | 0.0025 | 0.0028 | 0.0007 | 1.42 | 4.19 | -0.80 | 0.155 | **$2.9\times10^{-5}$** | 0.424 | 0.06 (-0.02,0.15) |
| Inferior temporal gyrus | 937/1029 | 0.0022 | -0.0006 | -0.0007 | 0.0026 | 0.0029 | 0.0007 | 0.84 | -0.19 | -1.05 | 0.399 | 0.846 | 0.293 | 0.04 (-0.05,0.13) |
| Insula | 950/1030 | 0.0039 | 0.0035 | -0.0014 | 0.0022 | 0.0024 | 0.0006 | 1.80 | 1.43 | -2.30 | 0.072 | 0.152 | **0.022** | 0.08 (-0.01,0.17) |
| Isthmus cingulate cortex | 951/1028 | 0.0000 | -0.0068 | -0.0007 | 0.0035 | 0.0039 | 0.0009 | 0.00 | -1.76 | -0.76 | 0.999 | 0.078 | 0.445 | $5.2\times10^{-5}$ (-0.09,0.09) |
| Lateral occipital cortex | 952/1035 | -0.0021 | -0.0025 | 0.0001 | 0.0023 | 0.0025 | 0.0006 | -0.88 | -0.99 | 0.13 | 0.377 | 0.324 | 0.893 | -0.04 (-0.13,0.05) |
| Lateral orbitofrontal cortex | 953/1035 | -0.0021 | 0.0058 | -0.0002 | 0.0019 | 0.0022 | 0.0005 | -1.10 | 2.68 | -0.29 | 0.273 | **0.008** | 0.774 | -0.05 (-0.14,0.04) |
| Lingual gyrus | 953/1035 | 0.0005 | 0.0019 | -0.0004 | 0.0024 | 0.0026 | 0.0006 | 0.19 | 0.73 | -0.57 | 0.848 | 0.465 | 0.569 | 0.01 (-0.08,0.1) |
| Medial orbitofrontal cortex | 946/1027 | 0.0074 | -0.0030 | -0.0002 | 0.0027 | 0.0030 | 0.0008 | 2.70 | -0.97 | -0.27 | **0.007** | 0.330 | 0.788 | 0.12 (0.03,0.21) |
| Middle temporal gyrus | 901/989 | 0.0030 | 0.0013 | 0.0009 | 0.0022 | 0.0024 | 0.0006 | 1.41 | 0.53 | 1.55 | 0.159 | 0.595 | 0.120 | 0.07 (-0.02,0.16) |
| Paracentral lobule | 953/1035 | -0.0056 | 0.0039 | 0.0005 | 0.0030 | 0.0033 | 0.0007 | -1.85 | 1.20 | 0.72 | 0.065 | 0.232 | 0.473 | -0.08 (-0.17,0) |
| Parahippocampal gyrus | 947/1028 | 0.0037 | 0.0115 | 0.0010 | 0.0035 | 0.0038 | 0.0009 | 1.05 | 3.00 | 1.14 | 0.292 | **0.003** | 0.255 | 0.05 (-0.04,0.14) |
| Pars opercularis of inferior frontal gyrus | 947/1032 | -0.0006 | 0.0030 | 0.0003 | 0.0038 | 0.0041 | 0.0009 | -0.15 | 0.73 | 0.32 | 0.878 | 0.464 | 0.751 | -0.01 (-0.1,0.08) |
| Pars orbitalis of inferior frontal gyrus | 953/1034 | 0.0022 | 0.0029 | -0.0012 | 0.0028 | 0.0031 | 0.0007 | 0.78 | 0.95 | -1.66 | 0.438 | 0.342 | 0.098 | 0.04 (-0.05,0.12) |
| Pars triangularis of inferior frontal gyrus | 946/1036 | 0.0036 | -0.0006 | -0.0011 | 0.0034 | 0.0038 | 0.0009 | 1.03 | -0.16 | -1.29 | 0.302 | 0.875 | 0.196 | 0.05 (-0.04,0.13) |
| Pericalcarine cortex | 951/1034 | 0.0033 | -0.0056 | -0.0007 | 0.0026 | 0.0029 | 0.0007 | 1.24 | -1.94 | -1.03 | 0.215 | 0.053 | 0.301 | 0.06 (-0.03,0.14) |
| Postcentral gyrus | 939/1020 | 0.0016 | -0.0046 | -0.0001 | 0.0021 | 0.0023 | 0.0006 | 0.75 | -1.98 | -0.22 | 0.454 | **0.048** | 0.823 | 0.03 (-0.05,0.12) |
| Posterior cingulate cortex | 950/1030 | -0.0001 | 0.0044 | 0.0009 | 0.0035 | 0.0038 | 0.0009 | -0.03 | 1.14 | 1.05 | 0.979 | 0.253 | 0.295 | -0.001 (-0.09,0.09) |
| Precentral gyrus | 942/1029 | 0.0005 | 0.0000 | -0.0005 | 0.0018 | 0.0020 | 0.0005 | 0.26 | 0.00 | -0.96 | 0.797 | 0.996 | 0.339 | 0.01 (-0.08,0.1) |
| Precuneus | 953/1032 | 0.0017 | 0.0058 | 0.0006 | 0.0019 | 0.0021 | 0.0005 | 0.90 | 2.78 | 1.29 | 0.371 | **0.005** | 0.196 | 0.04 (-0.05,0.13) |
| Rostral anterior cingulate cortex | 943/1029 | -0.0006 | 0.0058 | -0.0013 | 0.0046 | 0.0050 | 0.0012 | -0.14 | 1.15 | -1.09 | 0.892 | 0.249 | 0.275 | -0.01 (-0.09,0.08) |
| Rostral middle frontal gyrus | 952/1032 | -0.0024 | -0.0010 | 0.0004 | 0.0019 | 0.0021 | 0.0005 | -1.28 | -0.50 | 0.86 | 0.199 | 0.617 | 0.389 | -0.06 (-0.15,0.03) |
| Superior frontal gyrus | 950/1032 | 0.0017 | -0.0034 | 0.0008 | 0.0016 | 0.0017 | 0.0004 | 1.07 | -1.93 | 2.11 | 0.287 | 0.054 | **0.035** | 0.05 (-0.04,0.14) |
| Superior parietal cortex | 950/1033 | 0.0029 | 0.0021 | -0.0004 | 0.0020 | 0.0022 | 0.0005 | 1.44 | 0.94 | -0.75 | 0.149 | 0.347 | 0.453 | 0.07 (-0.02,0.15) |
| Superior temporal gyrus | 873/975 | 0.0017 | -0.0094 | 0.0002 | 0.0019 | 0.0021 | 0.0005 | 0.92 | -4.52 | 0.49 | 0.358 | **$6.7\times10^{-6}$** | 0.623 | 0.04 (-0.05,0.13) |
| Supramarginal gyrus | 938/1024 | 0.0008 | -0.0050 | 0.0011 | 0.0029 | 0.0033 | 0.0008 | 0.27 | -1.52 | 1.39 | 0.786 | 0.128 | 0.165 | 0.01 (-0.08,0.1) |
| Temporal pole | 947/1031 | 0.0033 | -0.0070 | 0.0013 | 0.0039 | 0.0042 | 0.0010 | 0.84 | -1.66 | 1.38 | 0.399 | 0.097 | 0.168 | 0.04 (-0.05,0.13) |
| Transverse temporal gyrus | 950/1034 | -0.0021 | 0.0016 | 0.0000 | 0.0036 | 0.0040 | 0.0009 | -0.58 | 0.41 | 0.01 | 0.559 | 0.684 | 0.991 | -0.03 (-0.11,0.06) |
| Total surface area | 953/1036 | 0.0008 | 0.0004 | 0.0001 | 0.0004 | 0.0004 | 0.0001 | 2.35 | 0.91 | 0.81 | **0.019** | 0.363 | 0.416 | 0.11 (0.02,0.19) |

P-values in **bold** are significant at the uncorrected level ($P < 0.05$).

**Table S4.** Full linear model results for the cortical thickness AIs in children

| Cortical thickness AI | | beta-coefficient | | | Standard Error | | | t-value | | | p-value | | | Cohen's d (95% CI) |
|---|---|---|---|---|---|---|---|---|---|---|---|---|---|---|
| | N cases / controls | diag | sex | age | diag | sex | age | diag | sex | age | diag | sex | age | diag |
| Banks of superior temporal sulcus | 884/962 | -0.0037 | 0.0021 | -0.0003 | 0.0018 | 0.0020 | 0.0005 | -2.05 | 1.06 | -0.57 | **0.040** | 0.291 | 0.570 | -0.1 (-0.19,0) |
| Caudal anterior cingulate cortex | 951/1029 | 0.0024 | -0.0003 | 0.0000 | 0.0024 | 0.0026 | 0.0007 | 1.00 | -0.13 | -0.02 | 0.316 | 0.900 | 0.982 | 0.05 (-0.04,0.13) |
| Caudal middle frontal cortex | 952/1035 | 0.0024 | -0.0011 | -0.0006 | 0.0012 | 0.0013 | 0.0003 | 2.07 | -0.83 | -1.94 | **0.038** | 0.408 | 0.052 | 0.09 (0.01,0.18) |
| Cuneus | 948/1035 | 0.0008 | 0.0016 | -0.0001 | 0.0016 | 0.0018 | 0.0004 | 0.48 | 0.93 | -0.26 | 0.634 | 0.355 | 0.791 | 0.02 (-0.07,0.11) |
| Entorhinal cortex | 910/1002 | -0.0041 | -0.0073 | -0.0006 | 0.0028 | 0.0031 | 0.0007 | -1.47 | -2.38 | -0.82 | 0.142 | **0.017** | 0.412 | -0.07 (-0.16,0.02) |
| Frontal pole | 952/1035 | 0.0037 | -0.0003 | -0.0005 | 0.0031 | 0.0034 | 0.0008 | 1.22 | -0.10 | -0.59 | 0.224 | 0.922 | 0.558 | 0.05 (-0.03,0.14) |
| Fusiform gyrus | 949/1032 | -0.0008 | 0.0004 | -0.0003 | 0.0010 | 0.0011 | 0.0003 | -0.78 | 0.34 | -1.16 | 0.435 | 0.737 | 0.246 | -0.04 (-0.12,0.05) |
| Inferior parietal cortex | 951/1031 | -0.0002 | -0.0008 | -0.0006 | 0.0009 | 0.0010 | 0.0002 | -0.24 | -0.75 | -2.50 | 0.808 | 0.451 | **0.013** | -0.01 (-0.1,0.08) |
| Inferior temporal gyrus | 937/1028 | -0.0013 | 0.0013 | 0.0005 | 0.0012 | 0.0014 | 0.0003 | -1.03 | 0.92 | 1.38 | 0.304 | 0.358 | 0.166 | -0.05 (-0.14,0.04) |
| Insula | 951/1031 | -0.0025 | 0.0007 | 0.0001 | 0.0012 | 0.0013 | 0.0003 | -2.07 | 0.50 | 0.30 | **0.038** | 0.615 | 0.767 | -0.09 (-0.18,-0.01) |
| Isthmus cingulate cortex | 950/1029 | -0.0012 | 0.0008 | 0.0012 | 0.0018 | 0.0020 | 0.0005 | -0.64 | 0.39 | 2.69 | 0.525 | 0.699 | **0.007** | -0.03 (-0.12,0.06) |
| Lateral occipital cortex | 952/1036 | 0.0006 | -0.0009 | 0.0004 | 0.0011 | 0.0012 | 0.0003 | 0.60 | -0.76 | 1.25 | 0.551 | 0.448 | 0.211 | 0.03 (-0.06,0.11) |
| Lateral orbitofrontal cortex | 953/1035 | -0.0005 | -0.0034 | -0.0002 | 0.0013 | 0.0015 | 0.0004 | -0.35 | -2.31 | -0.53 | 0.726 | **0.021** | 0.596 | -0.02 (-0.1,0.07) |
| Lingual gyrus | 953/1034 | -0.0010 | 0.0001 | 0.0008 | 0.0012 | 0.0013 | 0.0003 | -0.83 | 0.07 | 2.57 | 0.405 | 0.941 | **0.010** | -0.04 (-0.13,0.05) |
| Medial orbitofrontal cortex | 946/1028 | -0.0030 | -0.0059 | -0.0002 | 0.0017 | 0.0019 | 0.0005 | -1.73 | -3.10 | -0.42 | 0.084 | **0.002** | 0.671 | -0.08 (-0.17,0.01) |
| Middle temporal gyrus | 902/989 | -0.0003 | 0.0008 | 0.0004 | 0.0012 | 0.0013 | 0.0003 | -0.21 | 0.62 | 1.26 | 0.831 | 0.537 | 0.209 | -0.01 (-0.1,0.08) |
| Paracentral lobule | 953/1035 | -0.0019 | -0.0013 | -0.0008 | 0.0012 | 0.0013 | 0.0003 | -1.59 | -0.95 | -2.40 | 0.111 | 0.344 | **0.016** | -0.07 (-0.16,0.02) |
| Parahippocampal gyrus | 948/1029 | -0.0031 | -0.0056 | 0.0008 | 0.0021 | 0.0023 | 0.0005 | -1.50 | -2.46 | 1.55 | 0.135 | **0.014** | 0.122 | -0.07 (-0.16,0.02) |
| Pars opercularis of inferior frontal gyrus | 947/1032 | 0.0002 | 0.0015 | -0.0008 | 0.0013 | 0.0014 | 0.0004 | 0.18 | 1.01 | -2.21 | 0.855 | 0.314 | **0.027** | 0.01 (-0.08,0.1) |
| Pars orbitalis of inferior frontal gyrus | 953/1034 | 0.0024 | 0.0010 | 0.0004 | 0.0021 | 0.0024 | 0.0006 | 1.11 | 0.44 | 0.69 | 0.267 | 0.658 | 0.489 | 0.05 (-0.04,0.14) |
| Pars triangularis of inferior frontal gyrus | 946/1036 | -0.0003 | -0.0014 | -0.0001 | 0.0014 | 0.0016 | 0.0004 | -0.20 | -0.87 | -0.38 | 0.838 | 0.386 | 0.706 | -0.01 (-0.1,0.08) |
| Pericalcarine cortex | 949/1033 | 0.0003 | 0.0005 | -0.0008 | 0.0018 | 0.0021 | 0.0005 | 0.14 | 0.26 | -1.52 | 0.889 | 0.795 | 0.128 | 0.01 (-0.08,0.09) |
| Postcentral gyrus | 938/1022 | -0.0003 | -0.0013 | 0.0000 | 0.0010 | 0.0011 | 0.0003 | -0.26 | -1.16 | 0.08 | 0.791 | 0.245 | 0.938 | -0.01 (-0.1,0.08) |
| Posterior cingulate cortex | 949/1033 | -0.0007 | -0.0010 | 0.0002 | 0.0015 | 0.0017 | 0.0004 | -0.47 | -0.58 | 0.60 | 0.636 | 0.563 | 0.547 | -0.02 (-0.11,0.07) |
| Precentral gyrus | 942/1028 | 0.0023 | -0.0003 | -0.0001 | 0.0009 | 0.0010 | 0.0002 | 2.64 | -0.35 | -0.63 | **0.008** | 0.724 | 0.527 | 0.12 (0.03,0.21) |
| Precuneus | 953/1032 | 0.0002 | -0.0004 | 0.0000 | 0.0009 | 0.0010 | 0.0002 | 0.25 | -0.37 | -0.05 | 0.801 | 0.711 | 0.962 | 0.01 (-0.08,0.1) |
| Rostral anterior cingulate cortex | 943/1027 | -0.0005 | 0.0021 | -0.0001 | 0.0022 | 0.0024 | 0.0006 | -0.22 | 0.87 | -0.25 | 0.823 | 0.387 | 0.806 | -0.01 (-0.1,0.08) |
| Rostral middle frontal gyrus | 952/1033 | 0.0003 | 0.0001 | -0.0005 | 0.0011 | 0.0012 | 0.0003 | 0.29 | 0.09 | -1.81 | 0.774 | 0.926 | 0.070 | 0.01 (-0.08,0.1) |
| Superior frontal gyrus | 950/1032 | 0.0004 | -0.0009 | 0.0002 | 0.0008 | 0.0008 | 0.0002 | 0.47 | -1.09 | 0.94 | 0.637 | 0.277 | 0.346 | 0.02 (-0.07,0.11) |
| Superior parietal cortex | 950/1033 | 0.0002 | -0.0009 | -0.0002 | 0.0008 | 0.0009 | 0.0002 | 0.19 | -0.95 | -1.08 | 0.845 | 0.342 | 0.281 | 0.01 (-0.08,0.1) |
| Superior temporal gyrus | 874/978 | 0.0020 | 0.0022 | 0.0009 | 0.0011 | 0.0012 | 0.0003 | 1.78 | 1.80 | 2.86 | 0.075 | 0.073 | **0.004** | 0.08 (-0.01,0.17) |
| Supramarginal gyrus | 938/1027 | -0.0015 | 0.0011 | -0.0004 | 0.0011 | 0.0012 | 0.0003 | -1.40 | 0.96 | -1.57 | 0.161 | 0.335 | 0.116 | -0.06 (-0.15,0.02) |
| Temporal pole | 947/1030 | 0.0017 | -0.0002 | 0.0000 | 0.0028 | 0.0030 | 0.0007 | 0.63 | -0.07 | 0.02 | 0.530 | 0.945 | 0.984 | 0.03 (-0.06,0.12) |
| Transverse temporal gyrus | 950/1034 | 0.0013 | 0.0038 | 0.0006 | 0.0022 | 0.0024 | 0.0006 | 0.60 | 1.56 | 1.07 | 0.549 | 0.118 | 0.286 | 0.03 (-0.06,0.12) |
| Total average thickness | 953/1036 | 0.0000 | -0.0002 | 0.0000 | 0.0004 | 0.0004 | 0.0001 | -0.10 | -0.51 | 0.31 | 0.923 | 0.609 | 0.754 | -0.004 (-0.09,0.08) |

P-values in **bold** are significant at the uncorrected level ($P < 0.05$).

**Table S5.** Full linear model results for the subcortical volume AIs in adolescents

| Subcortical volume AI | | beta-coefficient | | | Standard Error | | | t-value | | | p-value | | | Cohen's d (95% CI) |
|---|---|---|---|---|---|---|---|---|---|---|---|---|---|---|
| | N cases/ controls | diag | sex | age | diag | sex | age | diag | sex | age | diag | sex | age | diag |
| Accumbens | 330/234 | -0.0054 | 0.0094 | 0.0016 | 0.0064 | 0.0074 | 0.0019 | -0.84 | 1.26 | 0.82 | 0.401 | 0.209 | 0.411 | -0.07 (-0.24,0.09) |
| Amygdala | 330/233 | 0.0021 | 0.0029 | -0.0013 | 0.0044 | 0.0050 | 0.0012 | 0.47 | 0.57 | -1.02 | 0.640 | 0.566 | 0.309 | 0.04 (-0.13,0.21) |
| Caudate Nucleus | 329/234 | 0.0002 | -0.0032 | 0.0009 | 0.0025 | 0.0029 | 0.0007 | 0.09 | -1.11 | 1.17 | 0.926 | 0.266 | 0.244 | 0.01 (-0.16,0.18) |
| Globus Pallidus | 330/233 | -0.0036 | -0.0050 | -0.0019 | 0.0047 | 0.0054 | 0.0014 | -0.77 | -0.92 | -1.34 | 0.444 | 0.357 | 0.181 | -0.07 (-0.24,0.1) |
| Hippocampus | 330/233 | 0.0040 | 0.0015 | -0.0001 | 0.0033 | 0.0037 | 0.0008 | 1.21 | 0.42 | -0.11 | 0.225 | 0.677 | 0.913 | 0.11 (-0.06,0.27) |
| Putamen | 329/227 | 0.0028 | -0.0020 | 0.0006 | 0.0026 | 0.0031 | 0.0008 | 1.07 | -0.65 | 0.72 | 0.284 | 0.514 | 0.469 | -0.03 (-0.2,0.14) |
| Thalamus[1] | 309/227 | -0.0054 | 0.0094 | 0.0016 | 0.0064 | 0.0074 | 0.0019 | -0.84 | 1.26 | 0.82 | 0.401 | 0.209 | 0.411 | 0.1 (-0.08,0.27) |

P-values in **bold** are significant at the uncorrected level (P < 0.05). [1]Thalamus volume was not available from the NIH dataset.

**Table S6.** Full linear model results for the cortical surface area AIs in adolescents

| Cortical surface area AI | | beta-coefficient | | | Standard Error | | | t-value | | | p-value | | | Cohen's d (95% CI) |
|---|---|---|---|---|---|---|---|---|---|---|---|---|---|---|
| | N cases / controls | diag | sex | age | diag | sex | age | diag | sex | age | diag | sex | age | diag |
| Banks of superior temporal sulcus | 384/323 | -0.0074 | 0.0011 | 0.0019 | 0.0066 | 0.0072 | 0.0017 | -1.13 | 0.16 | 1.08 | 0.259 | 0.877 | 0.280 | -0.09 (-0.24,0.06) |
| Caudal anterior cingulate cortex | 412/342 | -0.0095 | 0.0047 | 0.0015 | 0.0085 | 0.0094 | 0.0022 | -1.11 | 0.50 | 0.70 | 0.266 | 0.618 | 0.481 | -0.08 (-0.23,0.06) |
| Caudal middle frontal cortex | 412/340 | -0.0012 | -0.0023 | 0.0019 | 0.0054 | 0.0060 | 0.0014 | -0.22 | -0.39 | 1.37 | 0.824 | 0.697 | 0.172 | -0.02 (-0.16,0.13) |
| Cuneus | 412/341 | 0.0006 | 0.0029 | 0.0003 | 0.0045 | 0.0050 | 0.0012 | 0.13 | 0.58 | 0.27 | 0.894 | 0.560 | 0.790 | 0.01 (-0.13,0.15) |
| Entorhinal cortex | 402/326 | -0.0037 | -0.0158 | 0.0011 | 0.0083 | 0.0092 | 0.0021 | -0.44 | -1.72 | 0.52 | 0.659 | 0.087 | 0.603 | -0.03 (-0.18,0.11) |
| Frontal pole | 412/342 | 0.0054 | 0.0067 | 0.0035 | 0.0067 | 0.0074 | 0.0018 | 0.81 | 0.90 | 1.94 | 0.419 | 0.368 | 0.053 | 0.06 (-0.08,0.2) |
| Fusiform gyrus | 408/340 | 0.0035 | 0.0064 | -0.0008 | 0.0036 | 0.0039 | 0.0009 | 0.99 | 1.61 | -0.90 | 0.323 | 0.107 | 0.368 | 0.07 (-0.07,0.22) |
| Inferior parietal cortex | 410/339 | -0.0003 | 0.0016 | 0.0007 | 0.0037 | 0.0041 | 0.0009 | -0.07 | 0.38 | 0.72 | 0.940 | 0.704 | 0.472 | -0.01 (-0.15,0.14) |
| Inferior temporal gyrus | 388/331 | -0.0002 | -0.0096 | 0.0013 | 0.0040 | 0.0045 | 0.0011 | -0.06 | -2.15 | 1.21 | 0.954 | **0.032** | 0.227 | -0.004 (-0.15,0.14) |
| Insula | 408/339 | 0.0011 | 0.0054 | -0.0014 | 0.0033 | 0.0037 | 0.0010 | 0.35 | 1.48 | -1.40 | 0.730 | 0.139 | 0.161 | 0.03 (-0.12,0.17) |
| Isthmus cingulate cortex | 412/342 | -0.0095 | -0.0112 | 0.0022 | 0.0054 | 0.0060 | 0.0015 | -1.74 | -1.88 | 1.50 | 0.083 | 0.061 | 0.133 | -0.13 (-0.27,0.01) |
| Lateral occipital cortex | 412/342 | 0.0008 | 0.0097 | -0.0012 | 0.0034 | 0.0037 | 0.0009 | 0.23 | 2.62 | -1.46 | 0.822 | **0.009** | 0.145 | 0.02 (-0.13,0.16) |
| Lateral orbitofrontal cortex | 411/342 | -0.0022 | 0.0022 | 0.0006 | 0.0030 | 0.0033 | 0.0009 | -0.75 | 0.66 | 0.71 | 0.455 | 0.510 | 0.480 | -0.06 (-0.2,0.09) |
| Lingual gyrus | 409/339 | -0.0052 | -0.0031 | 0.0003 | 0.0038 | 0.0042 | 0.0011 | -1.39 | -0.75 | 0.23 | 0.165 | 0.456 | 0.815 | -0.1 (-0.25,0.04) |
| Medial orbitofrontal cortex | 410/341 | 0.0039 | -0.0052 | -0.0004 | 0.0041 | 0.0046 | 0.0012 | 0.94 | -1.13 | -0.36 | 0.347 | 0.258 | 0.716 | 0.07 (-0.07,0.21) |
| Middle temporal gyrus | 370/316 | -0.0017 | -0.0019 | 0.0011 | 0.0035 | 0.0039 | 0.0009 | -0.48 | -0.50 | 1.20 | 0.633 | 0.615 | 0.229 | -0.04 (-0.19,0.11) |
| Paracentral lobule | 411/342 | 0.0001 | 0.0097 | -0.0003 | 0.0046 | 0.0051 | 0.0012 | 0.03 | 1.91 | -0.23 | 0.977 | 0.057 | 0.817 | 0.002 (-0.14,0.15) |
| Parahippocampal gyrus | 409/340 | -0.0041 | 0.0115 | -0.0032 | 0.0051 | 0.0057 | 0.0015 | -0.80 | 2.02 | -2.21 | 0.425 | **0.044** | **0.027** | -0.06 (-0.2,0.08) |
| Pars opercularis of inferior frontal gyrus | 409/341 | 0.0085 | 0.0021 | -0.0020 | 0.0061 | 0.0067 | 0.0016 | 1.40 | 0.31 | -1.20 | 0.161 | 0.756 | 0.231 | 0.11 (-0.04,0.25) |
| Pars orbitalis of inferior frontal gyrus | 412/341 | 0.0108 | 0.0024 | -0.0013 | 0.0046 | 0.0050 | 0.0012 | 2.37 | 0.47 | -1.10 | **0.018** | 0.638 | 0.270 | 0.18 (0.03,0.32) |
| Pars triangularis of inferior frontal gyrus | 412/342 | 0.0083 | -0.0048 | -0.0002 | 0.0055 | 0.0060 | 0.0014 | 1.52 | -0.81 | -0.11 | 0.129 | 0.421 | 0.912 | 0.11 (-0.03,0.26) |
| Pericalcarine cortex | 411/342 | -0.0066 | -0.0058 | -0.0006 | 0.0042 | 0.0046 | 0.0011 | -1.59 | -1.26 | -0.54 | 0.112 | 0.207 | 0.587 | -0.12 (-0.26,0.02) |
| Postcentral gyrus | 408/340 | 0.0026 | -0.0017 | 0.0001 | 0.0033 | 0.0036 | 0.0008 | 0.80 | -0.49 | 0.13 | 0.424 | 0.627 | 0.900 | 0.06 (-0.08,0.2) |
| Posterior cingulate cortex | 412/342 | -0.0041 | 0.0035 | -0.0001 | 0.0058 | 0.0063 | 0.0015 | -0.71 | 0.56 | -0.05 | 0.477 | 0.578 | 0.958 | -0.05 (-0.2,0.09) |
| Precentral gyrus | 407/340 | -0.0053 | 0.0018 | -0.0005 | 0.0029 | 0.0032 | 0.0008 | -1.83 | 0.55 | -0.67 | 0.068 | 0.580 | 0.501 | -0.14 (-0.28,0.01) |
| Precuneus | 411/342 | -0.0030 | 0.0084 | 0.0005 | 0.0029 | 0.0031 | 0.0008 | -1.06 | 2.67 | 0.61 | 0.291 | **0.008** | 0.541 | -0.08 (-0.22,0.06) |
| Rostral anterior cingulate cortex | 410/341 | -0.0006 | -0.0005 | 0.0005 | 0.0071 | 0.0079 | 0.0018 | -0.09 | -0.06 | 0.26 | 0.931 | 0.951 | 0.796 | -0.01 (-0.15,0.14) |
| Rostral middle frontal gyrus | 412/342 | -0.0004 | -0.0024 | -0.0006 | 0.0030 | 0.0033 | 0.0008 | -0.15 | -0.72 | -0.75 | 0.885 | 0.474 | 0.456 | -0.01 (-0.15,0.13) |
| Superior frontal gyrus | 411/342 | 0.0041 | 0.0043 | -0.0009 | 0.0023 | 0.0025 | 0.0006 | 1.79 | 1.72 | -1.51 | 0.074 | 0.086 | 0.133 | 0.13 (-0.01,0.28) |
| Superior parietal cortex | 410/342 | 0.0023 | 0.0064 | -0.0010 | 0.0032 | 0.0035 | 0.0008 | 0.72 | 1.83 | -1.14 | 0.473 | 0.068 | 0.253 | 0.05 (-0.09,0.2) |
| Superior temporal gyrus | 361/314 | 0.0003 | -0.0135 | 0.0005 | 0.0029 | 0.0032 | 0.0008 | 0.11 | -4.19 | 0.67 | 0.911 | **0.00003** | 0.501 | 0.01 (-0.14,0.16) |
| Supramarginal gyrus | 409/338 | -0.0056 | -0.0082 | 0.0005 | 0.0042 | 0.0046 | 0.0011 | -1.34 | -1.78 | 0.45 | 0.180 | 0.076 | 0.652 | -0.1 (-0.25,0.04) |
| Temporal pole | 409/340 | 0.0045 | -0.0029 | -0.0009 | 0.0061 | 0.0067 | 0.0015 | 0.74 | -0.43 | -0.58 | 0.457 | 0.664 | 0.561 | 0.06 (-0.09,0.2) |
| Transverse temporal gyrus | 409/340 | 0.0051 | -0.0026 | 0.0019 | 0.0059 | 0.0066 | 0.0017 | 0.87 | -0.40 | 1.13 | 0.385 | 0.691 | 0.259 | 0.07 (-0.08,0.21) |
| Total surface area | 412/342 | 0.0001 | 0.0008 | -0.0001 | 0.0005 | 0.0006 | 0.0001 | 0.13 | 1.46 | -0.59 | 0.896 | 0.145 | 0.557 | 0.01 (-0.13,0.15) |

P-values in **bold** are significant at the uncorrected level ($P < 0.05$).

**Table S7**. Full linear model results for the cortical thickness AIs in adolescents.

| Cortical thickness AI | | beta-coefficient | | | Standard Error | | | t-value | | | p-value | | | Cohen's d (95% CI) |
|---|---|---|---|---|---|---|---|---|---|---|---|---|---|---|
| | N cases / controls | diag | sex | age | diag | sex | age | diag | sex | age | diag | sex | age | diag |
| Banks of superior temporal sulcus | 382/323 | -0.0022 | 0.0040 | -0.0007 | 0.0030 | 0.0033 | 0.0008 | -0.71 | 1.20 | -0.84 | 0.476 | 0.232 | 0.403 | -0.06 (-0.2,0.09) |
| Caudal anterior cingulate cortex | 411/342 | -0.0018 | -0.0038 | 0.0002 | 0.0041 | 0.0045 | 0.0012 | -0.45 | -0.84 | 0.17 | 0.651 | 0.400 | 0.864 | -0.03 (-0.18,0.11) |
| Caudal middle frontal cortex | 411/340 | 0.0034 | -0.0022 | -0.0007 | 0.0018 | 0.0020 | 0.0005 | 1.86 | -1.11 | -1.54 | 0.064 | 0.267 | 0.123 | 0.14 (0,0.28) |
| Cuneus | 411/341 | -0.0048 | -0.0043 | -0.0001 | 0.0024 | 0.0026 | 0.0007 | -2.01 | -1.62 | -0.11 | **0.045** | 0.106 | 0.913 | -0.15 (-0.29,-0.01) |
| Entorhinal cortex | 402/326 | 0.0008 | -0.0098 | -0.0013 | 0.0043 | 0.0049 | 0.0013 | 0.18 | -2.01 | -1.05 | 0.857 | **0.045** | 0.293 | 0.01 (-0.13,0.16) |
| Frontal pole | 411/342 | -0.0062 | -0.0035 | -0.0009 | 0.0047 | 0.0052 | 0.0012 | -1.32 | -0.67 | -0.75 | 0.188 | 0.503 | 0.451 | -0.1 (-0.24,0.05) |
| Fusiform gyrus | 407/340 | 0.0002 | -0.0015 | -0.0002 | 0.0017 | 0.0019 | 0.0005 | 0.11 | -0.80 | -0.48 | 0.911 | 0.421 | 0.632 | 0.01 (-0.14,0.15) |
| Inferior parietal cortex | 409/340 | -0.0010 | -0.0008 | -0.0003 | 0.0015 | 0.0017 | 0.0005 | -0.64 | -0.49 | -0.66 | 0.521 | 0.623 | 0.511 | -0.05 (-0.19,0.1) |
| Inferior temporal gyrus | 387/331 | 0.0012 | -0.0048 | 0.0007 | 0.0022 | 0.0025 | 0.0007 | 0.55 | -1.91 | 0.97 | 0.584 | 0.057 | 0.331 | 0.04 (-0.1,0.19) |
| Insula | 407/339 | -0.0023 | 0.0002 | -0.0005 | 0.0018 | 0.0020 | 0.0005 | -1.32 | 0.13 | -1.09 | 0.189 | 0.899 | 0.277 | -0.1 (-0.24,0.05) |
| Isthmus cingulate cortex | 410/341 | 0.0038 | -0.0027 | 0.0008 | 0.0028 | 0.0031 | 0.0007 | 1.35 | -0.88 | 1.06 | 0.177 | 0.381 | 0.291 | 0.1 (-0.04,0.24) |
| Lateral occipital cortex | 411/342 | -0.0020 | 0.0012 | 0.0005 | 0.0017 | 0.0019 | 0.0005 | -1.22 | 0.63 | 0.91 | 0.223 | 0.531 | 0.361 | -0.09 (-0.23,0.05) |
| Lateral orbitofrontal cortex | 410/342 | 0.0014 | -0.0031 | -0.0007 | 0.0021 | 0.0023 | 0.0006 | 0.67 | -1.36 | -1.08 | 0.503 | 0.173 | 0.279 | 0.05 (-0.09,0.19) |
| Lingual gyrus | 408/339 | 0.0001 | -0.0025 | -0.0001 | 0.0019 | 0.0021 | 0.0005 | 0.03 | -1.18 | -0.15 | 0.973 | 0.237 | 0.884 | 0.003 (-0.14,0.15) |
| Medial orbitofrontal cortex | 410/341 | 0.0032 | -0.0015 | 0.0002 | 0.0027 | 0.0030 | 0.0008 | 1.21 | -0.51 | 0.30 | 0.227 | 0.609 | 0.766 | 0.09 (-0.05,0.23) |
| Middle temporal gyrus | 369/318 | -0.0007 | -0.0055 | 0.0001 | 0.0020 | 0.0022 | 0.0006 | -0.34 | -2.43 | 0.21 | 0.737 | **0.015** | 0.831 | -0.03 (-0.18,0.12) |
| Paracentral lobule | 410/342 | 0.0026 | 0.0036 | 0.0008 | 0.0019 | 0.0021 | 0.0006 | 1.34 | 1.69 | 1.47 | 0.182 | 0.092 | 0.142 | 0.1 (-0.04,0.24) |
| Parahippocampal gyrus | 408/340 | -0.0069 | -0.0062 | -0.0012 | 0.0036 | 0.0040 | 0.0009 | -1.92 | -1.55 | -1.26 | 0.055 | 0.122 | 0.208 | -0.14 (-0.29,0) |
| Pars opercularis of inferior frontal gyrus | 408/341 | 0.0023 | -0.0031 | -0.0007 | 0.0021 | 0.0023 | 0.0005 | 1.13 | -1.36 | -1.25 | 0.258 | 0.176 | 0.213 | 0.08 (-0.06,0.23) |
| Pars orbitalis of inferior frontal gyrus | 411/341 | -0.0004 | 0.0019 | -0.0003 | 0.0039 | 0.0044 | 0.0012 | -0.09 | 0.44 | -0.28 | 0.928 | 0.658 | 0.779 | -0.01 (-0.15,0.14) |
| Pars triangularis of inferior frontal gyrus | 410/342 | 0.0019 | 0.0008 | -0.0002 | 0.0023 | 0.0025 | 0.0006 | 0.85 | 0.32 | -0.37 | 0.395 | 0.749 | 0.715 | 0.06 (-0.08,0.21) |
| Pericalcarine cortex | 410/342 | -0.0002 | -0.0031 | -0.0005 | 0.0029 | 0.0032 | 0.0008 | -0.06 | -0.95 | -0.58 | 0.954 | 0.342 | 0.560 | -0.004 (-0.15,0.14) |
| Postcentral gyrus | 406/340 | -0.0004 | -0.0025 | -0.0002 | 0.0017 | 0.0019 | 0.0005 | -0.27 | -1.36 | -0.39 | 0.791 | 0.174 | 0.700 | -0.02 (-0.16,0.12) |
| Posterior cingulate cortex | 411/342 | -0.0015 | 0.0002 | 0.0001 | 0.0024 | 0.0027 | 0.0007 | -0.61 | 0.07 | 0.08 | 0.541 | 0.946 | 0.934 | -0.05 (-0.19,0.1) |
| Precentral gyrus | 405/339 | -0.0012 | -0.0027 | -0.0007 | 0.0014 | 0.0015 | 0.0003 | -0.85 | -1.78 | -1.97 | 0.394 | 0.075 | **0.049** | -0.06 (-0.21,0.08) |
| Precuneus | 410/342 | 0.0020 | 0.0005 | 0.0004 | 0.0015 | 0.0017 | 0.0005 | 1.37 | 0.30 | 0.92 | 0.170 | 0.763 | 0.359 | 0.1 (-0.04,0.25) |
| Rostral anterior cingulate cortex | 409/341 | 0.0062 | -0.0018 | 0.0005 | 0.0036 | 0.0040 | 0.0011 | 1.71 | -0.45 | 0.45 | 0.087 | 0.652 | 0.650 | 0.13 (-0.02,0.27) |
| Rostral middle frontal gyrus | 411/342 | -0.0004 | -0.0015 | -0.0005 | 0.0016 | 0.0018 | 0.0005 | -0.27 | -0.82 | -0.93 | 0.784 | 0.412 | 0.351 | -0.02 (-0.16,0.12) |
| Superior frontal gyrus | 410/342 | 0.0020 | 0.0010 | -0.0006 | 0.0011 | 0.0013 | 0.0004 | 1.76 | 0.76 | -1.50 | 0.079 | 0.450 | 0.135 | 0.13 (-0.01,0.28) |
| Superior parietal cortex | 409/342 | -0.0010 | -0.0008 | -0.0007 | 0.0014 | 0.0015 | 0.0004 | -0.71 | -0.55 | -1.65 | 0.480 | 0.581 | 0.100 | -0.05 (-0.2,0.09) |
| Superior temporal gyrus | 360/314 | 0.0015 | 0.0001 | -0.0001 | 0.0018 | 0.0020 | 0.0005 | 0.84 | 0.03 | -0.27 | 0.403 | 0.977 | 0.786 | 0.07 (-0.09,0.22) |
| Supramarginal gyrus | 408/338 | -0.0014 | 0.0005 | -0.0007 | 0.0017 | 0.0019 | 0.0005 | -0.83 | 0.28 | -1.41 | 0.406 | 0.780 | 0.158 | -0.06 (-0.21,0.08) |
| Temporal pole | 407/340 | 0.0017 | 0.0026 | -0.0004 | 0.0040 | 0.0044 | 0.0011 | 0.44 | 0.60 | -0.37 | 0.661 | 0.548 | 0.713 | 0.03 (-0.11,0.18) |
| Transverse temporal gyrus | 408/340 | 0.0014 | 0.0024 | -0.0006 | 0.0040 | 0.0044 | 0.0010 | 0.36 | 0.55 | -0.55 | 0.718 | 0.581 | 0.583 | 0.03 (-0.12,0.17) |
| Total average thickness | 411/342 | 0.0002 | -0.0012 | -0.0002 | 0.0006 | 0.0006 | 0.0002 | 0.32 | -1.83 | -1.12 | 0.752 | 0.067 | 0.264 | 0.02 (-0.12,0.17) |

P-values in **bold** are significant at the uncorrected level ($P < 0.05$).

**Table S8.** Full linear model results for the subcortical volume AIs in adults

| Subcortical volume AI | | beta-coefficient | | | Standard Error | | | t-value | | | p-value | | | Cohen's d (95% CI) |
|---|---|---|---|---|---|---|---|---|---|---|---|---|---|---|
| | N cases/ controls | diag | sex | age | diag | sex | age | diag | sex | age | diag | sex | age | diag |
| Accumbens | 562/492 | 0.0004 | 0.0069 | 0.0000 | 0.0050 | 0.0052 | 0.0003 | 0.08 | 1.33 | -0.12 | 0.937 | 0.184 | 0.907 | 0.005 (-0.12,0.13) |
| Amygdala | 562/492 | -0.0013 | 0.0047 | 0.0000 | 0.0039 | 0.0040 | 0.0002 | -0.33 | 1.17 | -0.02 | 0.739 | 0.241 | 0.981 | -0.02 (-0.14,0.1) |
| Caudate Nucleus | 562/492 | 0.0014 | -0.0015 | 0.0001 | 0.0017 | 0.0017 | 0.0001 | 0.84 | -0.90 | 1.46 | 0.400 | 0.369 | 0.144 | 0.05 (-0.07,0.17) |
| Globus Pallidus | 563/491 | -0.0103 | -0.0049 | -0.0003 | 0.0035 | 0.0036 | 0.0002 | -2.94 | -1.34 | -1.37 | ***0.003*** | 0.180 | 0.171 | -0.18 (-0.3,-0.06) |
| Hippocampus | 562/491 | 0.0019 | 0.0001 | -0.0002 | 0.0026 | 0.0026 | 0.0001 | 0.73 | 0.04 | -1.83 | 0.463 | 0.970 | 0.067 | 0.05 (-0.08,0.17) |
| Putamen | 563/492 | 0.0012 | -0.0011 | 0.0002 | 0.0018 | 0.0019 | 0.0001 | 0.70 | -0.60 | 1.93 | 0.486 | 0.550 | 0.054 | -0.04 (-0.16,0.08) |
| Thalamus[1] | 563/490 | 0.0004 | 0.0069 | 0.0000 | 0.0050 | 0.0052 | 0.0003 | 0.08 | 1.33 | -0.12 | 0.937 | 0.184 | 0.907 | 0.04 (-0.08,0.16) |

P-values in **bold** are significant at the uncorrected level (P < 0.05). [1]Thalamus volume was not available from the NIH dataset.

**Table S9.** Full linear model results for the cortical surface area AIs in adults

| Cortical surface area AI | | beta-coefficient | | | Standard Error | | | t-value | | | p-value | | | Cohen's d (95% CI) |
|---|---|---|---|---|---|---|---|---|---|---|---|---|---|---|
| | N cases / controls | diag | sex | age | diag | sex | age | diag | sex | age | diag | sex | age | diag |
| Banks of superior temporal sulcus | 595/514 | 0.0011 | -0.0033 | 0.0002 | 0.0050 | 0.0051 | 0.0003 | 0.23 | -0.64 | 0.60 | 0.822 | 0.522 | 0.551 | 0.01 (-0.1,0.13) |
| Caudal anterior cingulate cortex | 610/538 | 0.0028 | 0.0093 | -0.0001 | 0.0073 | 0.0075 | 0.0004 | 0.38 | 1.24 | -0.17 | 0.705 | 0.214 | 0.864 | 0.02 (-0.09,0.14) |
| Caudal middle frontal cortex | 613/539 | 0.0054 | 0.0006 | -0.0007 | 0.0044 | 0.0045 | 0.0002 | 1.23 | 0.14 | -3.18 | 0.218 | 0.892 | **0.002** | 0.07 (-0.04,0.19) |
| Cuneus | 612/539 | -0.0080 | -0.0034 | -0.0002 | 0.0044 | 0.0045 | 0.0002 | -1.84 | -0.77 | -0.69 | 0.066 | 0.444 | 0.493 | -0.11 (-0.23,0.01) |
| Entorhinal cortex | 550/478 | -0.0111 | -0.0009 | 0.0006 | 0.0068 | 0.0071 | 0.0004 | -1.64 | -0.13 | 1.54 | 0.101 | 0.896 | 0.123 | -0.1 (-0.23,0.02) |
| Frontal pole | 613/539 | -0.0061 | 0.0053 | 0.0001 | 0.0054 | 0.0055 | 0.0003 | -1.14 | 0.97 | 0.52 | 0.253 | 0.331 | 0.606 | -0.07 (-0.18,0.05) |
| Fusiform gyrus | 567/493 | -0.0048 | 0.0073 | 0.0002 | 0.0030 | 0.0031 | 0.0002 | -1.59 | 2.36 | 1.18 | 0.112 | **0.018** | 0.238 | -0.1 (-0.22,0.02) |
| Inferior parietal cortex | 610/538 | -0.0004 | 0.0018 | 0.0001 | 0.0030 | 0.0031 | 0.0002 | -0.13 | 0.59 | 0.49 | 0.894 | 0.552 | 0.628 | -0.01 (-0.12,0.11) |
| Inferior temporal gyrus | 563/493 | 0.0036 | 0.0008 | 0.0001 | 0.0032 | 0.0033 | 0.0002 | 1.13 | 0.26 | 0.39 | 0.257 | 0.796 | 0.699 | 0.07 (-0.05,0.19) |
| Insula | 605/532 | -0.0012 | 0.0052 | -0.0001 | 0.0025 | 0.0026 | 0.0001 | -0.49 | 2.00 | -0.56 | 0.623 | **0.046** | 0.578 | -0.03 (-0.15,0.09) |
| Isthmus cingulate cortex | 613/539 | 0.0031 | -0.0085 | 0.0003 | 0.0044 | 0.0045 | 0.0002 | 0.70 | -1.88 | 1.26 | 0.484 | 0.060 | 0.208 | 0.04 (-0.07,0.16) |
| Lateral occipital cortex | 610/539 | 0.0055 | 0.0008 | 0.0000 | 0.0028 | 0.0029 | 0.0001 | 1.97 | 0.28 | 0.26 | **0.049** | 0.777 | 0.797 | 0.12 (0,0.23) |
| Lateral orbitofrontal cortex | 613/539 | -0.0017 | 0.0032 | 0.0001 | 0.0021 | 0.0021 | 0.0001 | -0.84 | 1.49 | 0.70 | 0.399 | 0.136 | 0.481 | -0.05 (-0.17,0.07) |
| Lingual gyrus | 568/494 | 0.0024 | 0.0023 | -0.0001 | 0.0034 | 0.0036 | 0.0002 | 0.69 | 0.63 | -0.78 | 0.489 | 0.527 | 0.436 | 0.04 (-0.08,0.16) |
| Medial orbitofrontal cortex | 611/539 | -0.0010 | -0.0052 | 0.0003 | 0.0033 | 0.0034 | 0.0002 | -0.29 | -1.52 | 1.63 | 0.768 | 0.128 | 0.103 | -0.02 (-0.13,0.1) |
| Middle temporal gyrus | 555/477 | -0.0002 | -0.0015 | 0.0001 | 0.0027 | 0.0027 | 0.0001 | -0.09 | -0.56 | 0.79 | 0.931 | 0.577 | 0.430 | -0.01 (-0.13,0.12) |
| Paracentral lobule | 612/538 | -0.0041 | 0.0052 | -0.0001 | 0.0038 | 0.0039 | 0.0002 | -1.07 | 1.34 | -0.66 | 0.285 | 0.181 | 0.510 | -0.06 (-0.18,0.05) |
| Parahippocampal gyrus | 568/491 | -0.0060 | 0.0100 | 0.0001 | 0.0040 | 0.0041 | 0.0002 | -1.50 | 2.45 | 0.54 | 0.134 | **0.014** | 0.587 | -0.09 (-0.21,0.03) |
| Pars opercularis of inferior frontal gyrus | 611/539 | 0.0026 | 0.0081 | 0.0000 | 0.0047 | 0.0048 | 0.0002 | 0.56 | 1.69 | -0.11 | 0.574 | 0.092 | 0.910 | 0.03 (-0.08,0.15) |
| Pars orbitalis of inferior frontal gyrus | 612/539 | 0.0022 | 0.0049 | 0.0004 | 0.0036 | 0.0037 | 0.0002 | 0.61 | 1.34 | 2.38 | 0.541 | 0.179 | **0.017** | 0.04 (-0.08,0.15) |
| Pars triangularis of inferior frontal gyrus | 612/538 | -0.0026 | -0.0007 | 0.0002 | 0.0046 | 0.0047 | 0.0002 | -0.57 | -0.14 | 0.86 | 0.571 | 0.885 | 0.392 | -0.03 (-0.15,0.08) |
| Pericalcarine cortex | 612/539 | 0.0000 | -0.0027 | -0.0003 | 0.0039 | 0.0040 | 0.0002 | 0.01 | -0.68 | -1.43 | 0.996 | 0.494 | 0.154 | 0.0003 (-0.12,0.12) |
| Postcentral gyrus | 607/528 | -0.0001 | 0.0004 | -0.0001 | 0.0027 | 0.0028 | 0.0001 | -0.03 | 0.14 | -0.84 | 0.975 | 0.892 | 0.398 | -0.002 (-0.12,0.11) |
| Posterior cingulate cortex | 613/539 | 0.0010 | 0.0049 | 0.0004 | 0.0046 | 0.0047 | 0.0002 | 0.22 | 1.04 | 1.59 | 0.822 | 0.297 | 0.111 | 0.01 (-0.1,0.13) |
| Precentral gyrus | 609/537 | -0.0047 | 0.0024 | 0.0000 | 0.0024 | 0.0025 | 0.0001 | -1.92 | 0.96 | -0.22 | 0.055 | 0.338 | 0.822 | -0.11 (-0.23,0) |
| Precuneus | 612/539 | 0.0011 | 0.0055 | 0.0001 | 0.0025 | 0.0025 | 0.0001 | 0.45 | 2.17 | 0.76 | 0.654 | **0.030** | 0.450 | 0.03 (-0.09,0.14) |
| Rostral anterior cingulate cortex | 610/535 | -0.0055 | 0.0035 | 0.0005 | 0.0060 | 0.0062 | 0.0003 | -0.92 | 0.56 | 1.64 | 0.359 | 0.576 | 0.102 | -0.05 (-0.17,0.06) |
| Rostral middle frontal gyrus | 613/538 | -0.0012 | 0.0008 | 0.0001 | 0.0024 | 0.0024 | 0.0001 | -0.53 | 0.34 | 0.90 | 0.598 | 0.732 | 0.371 | -0.03 (-0.15,0.08) |
| Superior frontal gyrus | 610/535 | -0.0032 | -0.0008 | 0.0000 | 0.0021 | 0.0021 | 0.0001 | -1.57 | -0.40 | -0.41 | 0.116 | 0.692 | 0.685 | -0.09 (-0.21,0.02) |
| Superior parietal cortex | 610/539 | -0.0010 | -0.0001 | 0.0000 | 0.0025 | 0.0026 | 0.0001 | -0.41 | -0.06 | 0.31 | 0.683 | 0.954 | 0.760 | -0.02 (-0.14,0.09) |
| Superior temporal gyrus | 547/476 | -0.0034 | -0.0095 | -0.0002 | 0.0026 | 0.0027 | 0.0001 | -1.30 | -3.52 | -1.79 | 0.196 | **0.0004** | 0.073 | -0.08 (-0.2,0.04) |
| Supramarginal gyrus | 608/535 | -0.0045 | -0.0093 | -0.0002 | 0.0037 | 0.0038 | 0.0002 | -1.22 | -2.46 | -1.12 | 0.223 | **0.014** | 0.262 | -0.07 (-0.19,0.04) |
| Temporal pole | 568/494 | -0.0044 | -0.0125 | 0.0000 | 0.0048 | 0.0050 | 0.0003 | -0.92 | -2.51 | 0.16 | 0.358 | **0.012** | 0.876 | -0.06 (-0.18,0.06) |
| Transverse temporal gyrus | 568/494 | 0.0004 | 0.0040 | 0.0002 | 0.0051 | 0.0052 | 0.0003 | 0.07 | 0.78 | 0.91 | 0.941 | 0.438 | 0.361 | 0.005 (-0.12,0.13) |
| Total surface area | 613/539 | -0.0008 | 0.0005 | 0.0000 | 0.0006 | 0.0007 | 0.0000 | -1.17 | 0.71 | 0.23 | 0.240 | 0.478 | 0.820 | -0.07 (-0.19,0.05) |

P-values in **bold** are significant at the uncorrected level ($P < 0.05$).

**Table S10.** Full linear model results for the cortical thickness AIs in adults

| Cortical thickness AI | | beta-coefficient | | | Standard Error | | | t-value | | | p-value | | | Cohen's d (95% CI) |
|---|---|---|---|---|---|---|---|---|---|---|---|---|---|---|
| | N cases / controls | diag | sex | age | diag | sex | age | diag | sex | age | diag | sex | age | diag |
| Banks of superior temporal sulcus | 595/514 | -0.0011 | 0.0008 | -0.0001 | 0.0024 | 0.0025 | 0.0001 | -0.48 | 0.31 | -0.52 | 0.635 | 0.758 | 0.603 | -0.03 (-0.15,0.09) |
| Caudal anterior cingulate cortex | 610/538 | 0.0061 | 0.0042 | 0.0000 | 0.0033 | 0.0034 | 0.0002 | 1.87 | 1.26 | -0.19 | 0.062 | 0.210 | 0.846 | 0.11 (0,0.23) |
| Caudal middle frontal cortex | 613/539 | 0.0005 | -0.0019 | -0.0001 | 0.0015 | 0.0016 | 0.0001 | 0.30 | -1.20 | -1.04 | 0.766 | 0.231 | 0.300 | 0.02 (-0.1,0.13) |
| Cuneus | 612/539 | 0.0033 | 0.0001 | 0.0001 | 0.0018 | 0.0019 | 0.0001 | 1.82 | 0.08 | 1.20 | 0.069 | 0.940 | 0.232 | 0.11 (-0.01,0.22) |
| Entorhinal cortex | 550/478 | -0.0017 | 0.0018 | -0.0002 | 0.0036 | 0.0038 | 0.0002 | -0.48 | 0.47 | -1.09 | 0.633 | 0.640 | 0.276 | -0.03 (-0.15,0.09) |
| Frontal pole | 613/539 | 0.0044 | -0.0070 | -0.0002 | 0.0033 | 0.0034 | 0.0002 | 1.32 | -2.06 | -1.14 | 0.185 | **0.040** | 0.256 | 0.08 (-0.04,0.19) |
| Fusiform gyrus | 567/493 | 0.0004 | 0.0003 | -0.0001 | 0.0014 | 0.0015 | 0.0001 | 0.32 | 0.18 | -1.28 | 0.749 | 0.860 | 0.201 | 0.02 (-0.1,0.14) |
| Inferior parietal cortex | 610/538 | 0.0007 | 0.0028 | 0.0000 | 0.0012 | 0.0012 | 0.0001 | 0.61 | 2.29 | -0.53 | 0.542 | **0.022** | 0.599 | 0.04 (-0.08,0.15) |
| Inferior temporal gyrus | 563/493 | -0.0003 | 0.0011 | -0.0002 | 0.0015 | 0.0016 | 0.0001 | -0.17 | 0.65 | -1.98 | 0.865 | 0.515 | **0.048** | -0.01 (-0.13,0.11) |
| Insula | 605/532 | -0.0001 | -0.0006 | -0.0001 | 0.0014 | 0.0015 | 0.0001 | -0.09 | -0.39 | -1.26 | 0.932 | 0.696 | 0.208 | -0.01 (-0.12,0.11) |
| Isthmus cingulate cortex | 613/539 | -0.0021 | 0.0002 | 0.0000 | 0.0022 | 0.0023 | 0.0001 | -0.95 | 0.09 | -0.23 | 0.340 | 0.931 | 0.822 | -0.06 (-0.17,0.06) |
| Lateral occipital cortex | 610/539 | 0.0026 | 0.0011 | 0.0001 | 0.0012 | 0.0012 | 0.0001 | 2.22 | 0.94 | 1.08 | **0.026** | 0.347 | 0.280 | 0.13 (0.02,0.25) |
| Lateral orbitofrontal cortex | 613/539 | 0.0023 | -0.0035 | -0.0001 | 0.0015 | 0.0016 | 0.0001 | 1.47 | -2.19 | -1.20 | 0.143 | **0.028** | 0.229 | 0.09 (-0.03,0.2) |
| Lingual gyrus | 568/494 | -0.0008 | -0.0007 | -0.0001 | 0.0015 | 0.0015 | 0.0001 | -0.53 | -0.48 | -1.32 | 0.595 | 0.634 | 0.188 | -0.03 (-0.15,0.09) |
| Medial orbitofrontal cortex | 611/539 | 0.0041 | -0.0043 | -0.0001 | 0.0020 | 0.0020 | 0.0001 | 2.07 | -2.12 | -0.60 | **0.039** | **0.034** | 0.548 | 0.12 (0.01,0.24) |
| Middle temporal gyrus | 555/477 | -0.0038 | -0.0031 | 0.0001 | 0.0015 | 0.0015 | 0.0001 | -2.61 | -2.02 | 0.84 | **0.009** | **0.043** | 0.400 | -0.16 (-0.29,-0.04) |
| Paracentral lobule | 612/538 | -0.0005 | 0.0010 | 0.0000 | 0.0015 | 0.0016 | 0.0001 | -0.30 | 0.62 | 0.32 | 0.767 | 0.536 | 0.750 | -0.02 (-0.13,0.1) |
| Parahippocampal gyrus | 568/491 | 0.0026 | 0.0046 | 0.0000 | 0.0030 | 0.0031 | 0.0002 | 0.85 | 1.46 | -0.02 | 0.395 | 0.143 | 0.985 | 0.05 (-0.07,0.17) |
| Pars opercularis of inferior frontal gyrus | 611/539 | -0.0003 | 0.0020 | 0.0000 | 0.0017 | 0.0018 | 0.0001 | -0.17 | 1.09 | -0.07 | 0.864 | 0.277 | 0.947 | -0.01 (-0.13,0.11) |
| Pars orbitalis of inferior frontal gyrus | 612/539 | 0.0020 | -0.0003 | 0.0000 | 0.0023 | 0.0024 | 0.0001 | 0.86 | -0.10 | 0.15 | 0.390 | 0.917 | 0.877 | 0.05 (-0.06,0.17) |
| Pars triangularis of inferior frontal gyrus | 612/538 | -0.0002 | 0.0010 | 0.0000 | 0.0019 | 0.0019 | 0.0001 | -0.13 | 0.52 | -0.26 | 0.900 | 0.602 | 0.796 | -0.01 (-0.12,0.11) |
| Pericalcarine cortex | 612/539 | 0.0061 | -0.0002 | -0.0002 | 0.0021 | 0.0022 | 0.0001 | 2.88 | -0.10 | -1.57 | **0.004** | 0.922 | 0.117 | 0.17 (0.06,0.29) |
| Postcentral gyrus | 607/528 | -0.0034 | -0.0011 | 0.0000 | 0.0013 | 0.0014 | 0.0001 | -2.52 | -0.82 | 0.02 | **0.012** | 0.411 | 0.985 | -0.15 (-0.27,-0.03) |
| Posterior cingulate cortex | 613/539 | -0.0004 | 0.0005 | -0.0002 | 0.0018 | 0.0018 | 0.0001 | -0.21 | 0.29 | -1.77 | 0.836 | 0.769 | 0.076 | -0.01 (-0.13,0.1) |
| Precentral gyrus | 609/537 | 0.0018 | -0.0029 | -0.0001 | 0.0013 | 0.0013 | 0.0001 | 1.39 | -2.20 | -1.65 | 0.164 | **0.028** | 0.100 | 0.08 (-0.03,0.2) |
| Precuneus | 612/539 | 0.0004 | -0.0013 | -0.0001 | 0.0012 | 0.0012 | 0.0001 | 0.34 | -1.04 | -1.04 | 0.731 | 0.301 | 0.297 | 0.02 (-0.1,0.14) |
| Rostral anterior cingulate cortex | 610/535 | 0.0028 | 0.0015 | -0.0004 | 0.0030 | 0.0031 | 0.0002 | 0.94 | 0.47 | -2.16 | 0.345 | 0.636 | **0.031** | 0.06 (-0.06,0.17) |
| Rostral middle frontal gyrus | 613/538 | -0.0010 | -0.0006 | -0.0001 | 0.0012 | 0.0013 | 0.0001 | -0.86 | -0.46 | -1.10 | 0.391 | 0.646 | 0.270 | -0.05 (-0.17,0.06) |
| Superior frontal gyrus | 610/535 | 0.0005 | -0.0011 | -0.0001 | 0.0009 | 0.0009 | 0.0001 | 0.59 | -1.22 | -1.76 | 0.556 | 0.221 | 0.079 | 0.04 (-0.08,0.15) |
| Superior parietal cortex | 610/539 | 0.0002 | -0.0006 | 0.0000 | 0.0010 | 0.0010 | 0.0001 | 0.15 | -0.63 | 0.90 | 0.879 | 0.527 | 0.367 | 0.01 (-0.11,0.12) |
| Superior temporal gyrus | 547/476 | -0.0014 | 0.0009 | 0.0000 | 0.0015 | 0.0016 | 0.0001 | -0.91 | 0.57 | -0.28 | 0.365 | 0.572 | 0.777 | -0.06 (-0.18,0.07) |
| Supramarginal gyrus | 608/535 | -0.0001 | 0.0015 | -0.0001 | 0.0014 | 0.0015 | 0.0001 | -0.09 | 1.04 | -0.76 | 0.930 | 0.300 | 0.447 | -0.01 (-0.12,0.11) |
| Temporal pole | 568/494 | -0.0016 | 0.0019 | 0.0001 | 0.0033 | 0.0034 | 0.0002 | -0.48 | 0.54 | 0.72 | 0.633 | 0.586 | 0.474 | -0.03 (-0.15,0.09) |
| Transverse temporal gyrus | 568/494 | -0.0027 | 0.0031 | -0.0003 | 0.0029 | 0.0029 | 0.0001 | -0.95 | 1.06 | -1.79 | 0.344 | 0.289 | 0.073 | -0.06 (-0.18,0.06) |
| Total average thickness | 613/539 | 0.0001 | -0.0002 | -0.0001 | 0.0004 | 0.0004 | 0.0000 | 0.28 | -0.56 | -2.44 | 0.780 | 0.577 | **0.015** | 0.02 (-0.1,0.13) |

P-values in **bold** are significant at the uncorrected level ($P < 0.05$).

**Table S11.** Full linear model results for the subcortical volume AIs in all age groups combined.

| Subcortical volume AI | | beta-coefficient | | | Standard Error | | | t-value | | | p-value | | | Cohen's d (95% CI) |
|---|---|---|---|---|---|---|---|---|---|---|---|---|---|---|
| | N cases/ controls | diag | age | sex | diag | age | sex | diag | age | sex | diag | age | sex | diag |
| Accumbens | 1734/1654 | -0.0027 | -0.0002 | 0.0032 | 0.0026 | 0.0002 | 0.0028 | -1.06 | -0.97 | 1.15 | 0.289 | 0.333 | 0.251 | -0.037 (-0.1,0.03) |
| Amygdala | 1733/1653 | -0.0005 | -0.0002 | 0.0035 | 0.0019 | 0.0001 | 0.0021 | -0.27 | -1.35 | 1.64 | 0.788 | 0.176 | 0.102 | -0.009 (-0.08,0.06) |
| Caudate Nucleus | 1734/1654 | 0.0008 | 0.0002 | -0.0024 | 0.0010 | 0.0001 | 0.0011 | 0.77 | 2.14 | -2.29 | 0.441 | 0.032 | **0.022** | 0.027 (-0.04,0.09) |
| Globus Pallidus | 1736/1650 | -0.0040 | -0.0006 | -0.0027 | 0.0019 | 0.0001 | 0.0021 | -2.13 | -4.55 | -1.33 | **0.033** | **$5.6 \times 10^{-6}$** | 0.185 | -0.074 (-0.14,-0.01) |
| Hippocampus | 1733/1650 | 0.0005 | -0.0002 | 0.0002 | 0.0014 | 0.0001 | 0.0015 | 0.37 | -2.03 | 0.13 | 0.708 | **0.042** | 0.900 | 0.013 (-0.05,0.08) |
| Putamen | 1735/1644 | -0.0015 | 0.0003 | 0.0010 | 0.0011 | 0.0001 | 0.0012 | -1.40 | 4.54 | 0.83 | 0.162 | **$5.7 \times 10^{-6}$** | 0.406 | -0.048 (-0.12,0.02) |
| Thalamus[1] | 1619/1536 | 0.0013 | 0.0008 | -0.0008 | 0.0010 | 0.0001 | 0.0011 | 1.27 | 10.29 | -0.66 | 0.202 | **$2.0 \times 10^{-24}$** | 0.507 | 0.046 (-0.02,0.12) |

P-values in **bold** are significant at the uncorrected level ($P < 0.05$). **[1]**Thalamus volume was not available from the NIH dataset

**Table S12.** Full linear model results for the cortical surface area AI in all age groups combined.

| Surface area AI | | beta-coefficient | | | Standard Error | | | t-value | | | p-value | | | Cohen's d (95% CI) |
|---|---|---|---|---|---|---|---|---|---|---|---|---|---|---|
| | N cases / controls | diag | age | sex | diag | age | sex | diag | age | sex | diag | age | sex | diag |
| Banks of superior temporal sulcus | 1862/1799 | -0.0025 | 0.0002 | -0.0005 | 0.0028 | 0.0002 | 0.0031 | -0.88 | 1.02 | -0.17 | 0.378 | 0.308 | 0.866 | -0.029 (-0.09,0.04) |
| Caudal anterior cingulate cortex | 1973/1909 | -0.0021 | -0.0004 | 0.0066 | 0.0039 | 0.0002 | 0.0041 | -0.55 | -1.87 | 1.60 | 0.582 | 0.061 | 0.110 | -0.018 (-0.08,0.05) |
| Caudal middle frontal cortex | 1978/1913 | 0.0033 | -0.0005 | $-4.3\times10^{-5}$ | 0.0024 | 0.0002 | 0.0026 | 1.36 | -3.45 | -0.02 | 0.174 | **0.001** | 0.987 | 0.044 (-0.02,0.11) |
| Cuneus | 1971/1914 | -0.0004 | 0.0003 | 0.0006 | 0.0022 | 0.0001 | 0.0024 | -0.17 | 2.35 | 0.27 | 0.867 | **0.019** | 0.790 | -0.005 (-0.07,0.06) |
| Entorhinal cortex | 1862/1805 | -0.0043 | 0.0007 | -0.0079 | 0.0039 | 0.0002 | 0.0042 | -1.10 | 3.02 | -1.88 | 0.273 | **0.003** | 0.060 | -0.036 (-0.1,0.03) |
| Frontal pole | 1978/1916 | -0.0046 | 0.0001 | 0.0046 | 0.0030 | 0.0001 | 0.0031 | -1.57 | 0.45 | 1.45 | 0.117 | 0.649 | 0.148 | -0.051 (-0.11,0.01) |
| Fusiform gyrus | 1922/1864 | -0.0026 | $4.03\times10^{-5}$ | 0.0043 | 0.0017 | 0.0001 | 0.0018 | -1.56 | 0.50 | 2.40 | 0.120 | 0.614 | **0.016** | -0.051 (-0.11,0.01) |
| Inferior parietal cortex | 1970/1906 | 0.0017 | -0.0001 | 0.0068 | 0.0017 | 0.0001 | 0.0019 | 1.01 | -0.99 | 3.64 | 0.312 | 0.320 | **$2.8\times10^{-4}$** | 0.033 (-0.03,0.1) |
| Inferior temporal gyrus | 1888/1853 | 0.0023 | $4.6\times10^{-5}$ | -0.0014 | 0.0018 | 0.0001 | 0.0020 | 1.30 | 0.45 | -0.74 | 0.193 | 0.651 | 0.458 | 0.043 (-0.02,0.11) |
| Insula | 1963/1901 | 0.0018 | -0.0004 | 0.0047 | 0.0015 | 0.0001 | 0.0016 | 1.19 | -3.67 | 2.90 | 0.236 | **$2.45\times10^{-4}$** | **0.004** | 0.038 (-0.02,0.1) |
| Isthmus cingulate cortex | 1976/1909 | -0.0009 | -0.0001 | -0.0078 | 0.0024 | 0.0001 | 0.0026 | -0.36 | -0.76 | -2.96 | 0.718 | 0.448 | **0.003** | -0.012 (-0.07,0.05) |
| Lateral occipital cortex | 1974/1916 | 0.0007 | 0.0001 | 0.0006 | 0.0016 | 0.0001 | 0.0017 | 0.45 | 1.11 | 0.34 | 0.656 | 0.268 | 0.733 | 0.014 (-0.05,0.08) |
| Lateral orbitofrontal cortex | 1977/1916 | -0.0022 | 0.0001 | 0.0044 | 0.0013 | 0.0001 | 0.0014 | -1.72 | 1.27 | 3.12 | 0.085 | 0.204 | **0.002** | -0.056 (-0.12,0.01) |
| Lingual gyrus | 1930/1868 | 0.0002 | 0.0001 | 0.0013 | 0.0017 | 0.0001 | 0.0019 | 0.14 | 0.52 | 0.70 | 0.886 | 0.603 | 0.485 | 0.005 (-0.06,0.07) |
| Medial orbitofrontal cortex | 1967/1907 | 0.0040 | 0.0003 | -0.0038 | 0.0019 | 0.0001 | 0.0020 | 2.16 | 2.62 | -1.88 | **0.031** | **0.009** | 0.060 | 0.07 (0.01,0.13) |
| Middle temporal gyrus | 1826/1782 | 0.0012 | 0.0002 | 0.0003 | 0.0015 | 0.0001 | 0.0017 | 0.77 | 2.09 | 0.16 | 0.443 | **0.037** | 0.869 | 0.026 (-0.04,0.09) |
| Paracentral lobule | 1976/1915 | -0.0040 | -0.0002 | 0.0050 | 0.0021 | 0.0001 | 0.0023 | -1.89 | -1.56 | 2.20 | 0.059 | 0.119 | **0.028** | -0.061 (-0.12,0) |
| Parahippocampal gyrus | 1924/1859 | -0.0004 | -0.0001 | 0.0104 | 0.0024 | 0.0001 | 0.0026 | -0.19 | -0.80 | 4.07 | 0.852 | 0.423 | **$4.8\times10^{-5}$** | -0.006 (-0.07,0.06) |
| Pars opercularis of inferior frontal gyrus | 1967/1912 | 0.0020 | $2.7\times10^{-5}$ | 0.0041 | 0.0026 | 0.0001 | 0.0028 | 0.74 | 0.23 | 1.48 | 0.459 | 0.816 | 0.140 | 0.024 (-0.04,0.09) |
| Pars orbitalis of inferior frontal gyrus | 1977/1914 | 0.0039 | 0.0001 | 0.0040 | 0.0020 | 0.0001 | 0.0021 | 1.95 | 1.26 | 1.84 | 0.051 | 0.208 | 0.065 | 0.063 (0,0.13) |
| Pars triangularis of inferior frontal gyrus | 1970/1916 | 0.0030 | -0.0001 | -0.0010 | 0.0025 | 0.0001 | 0.0026 | 1.23 | -0.44 | -0.39 | 0.220 | 0.659 | 0.698 | 0.04 (-0.02,0.1) |
| Pericalcarine cortex | 1974/1915 | 0.0004 | 0.0001 | -0.0041 | 0.0019 | 0.0001 | 0.0021 | 0.21 | 1.29 | -1.95 | 0.836 | 0.197 | 0.051 | 0.007 (-0.06,0.07) |
| Postcentral gyrus | 1954/1888 | 0.0013 | 0.0001 | -0.0022 | 0.0015 | 0.0001 | 0.0016 | 0.89 | 0.68 | -1.40 | 0.374 | 0.493 | 0.160 | 0.029 (-0.03,0.09) |
| Posterior cingulate cortex | 1975/1911 | -0.0004 | 0.0001 | 0.0040 | 0.0025 | 0.0001 | 0.0027 | -0.16 | 0.91 | 1.50 | 0.876 | 0.361 | 0.134 | -0.005 (-0.07,0.06) |
| Precentral gyrus | 1958/1906 | -0.0022 | $-3.7\times10^{-6}$ | 0.0015 | 0.0013 | 0.0001 | 0.0014 | -1.68 | -0.05 | 1.10 | 0.092 | 0.956 | 0.272 | -0.054 (-0.12,0.01) |
| Precuneus | 1976/1913 | 0.0007 | $-7.8\times10^{-6}$ | 0.0060 | 0.0013 | 0.0001 | 0.0014 | 0.56 | -0.12 | 4.19 | 0.574 | 0.903 | **$2.8\times10^{-5}$** | 0.018 (-0.04,0.08) |
| Rostral anterior cingulate cortex | 1963/1905 | -0.0024 | -0.0001 | 0.0031 | 0.0032 | 0.0002 | 0.0035 | -0.74 | -0.58 | 0.88 | 0.462 | 0.565 | 0.380 | -0.024 (-0.09,0.04) |
| Rostral middle frontal gyrus | 1977/1912 | -0.0017 | -0.0001 | -0.0008 | 0.0013 | 0.0001 | 0.0014 | -1.25 | -1.57 | -0.55 | 0.210 | 0.117 | 0.582 | -0.04 (-0.1,0.02) |
| Superior frontal gyrus | 1971/1909 | 0.0007 | 0.0001 | -0.0015 | 0.0011 | 0.0001 | 0.0012 | 0.61 | 1.67 | -1.23 | 0.544 | 0.095 | 0.218 | 0.02 (-0.04,0.08) |
| Superior parietal cortex | 1970/1914 | 0.0014 | -0.0002 | 0.0021 | 0.0014 | 0.0001 | 0.0015 | 0.96 | -2.53 | 1.39 | 0.338 | **0.011** | 0.165 | 0.031 (-0.03,0.09) |
| Superior temporal gyrus | 1781/1765 | $9.6\times10^{-6}$ | 0.0000 | -0.0101 | 0.0014 | 0.0001 | 0.0015 | 0.01 | -0.03 | -6.85 | 0.994 | 0.979 | **$8.6\times10^{-12}$** | 0 (-0.07,0.07) |
| Supramarginal gyrus | 1955/1897 | -0.0022 | -0.0002 | -0.0072 | 0.0020 | 0.0001 | 0.0022 | -1.08 | -1.77 | -3.30 | 0.279 | 0.076 | **0.001** | -0.035 (-0.1,0.03) |
| Temporal pole | 1924/1865 | 0.0009 | 0.0001 | -0.0082 | 0.0027 | 0.0002 | 0.0029 | 0.35 | 0.98 | -2.80 | 0.727 | 0.326 | **0.005** | 0.011 (-0.05,0.08) |
| Transverse temporal gyrus | 1927/1868 | -0.0001 | 0.0003 | 0.0024 | 0.0026 | 0.0002 | 0.0029 | -0.02 | 1.61 | 0.82 | 0.980 | 0.108 | 0.409 | -0.001 (-0.06,0.06) |
| Total surface area | 1978/1917 | 0.0002 | $-1.7\times10^{-6}$ | 0.0005 | 0.0003 | $1.9\times10^{-5}$ | 0.0003 | 0.80 | -0.09 | 1.63 | 0.425 | 0.926 | 0.104 | 0.026 (-0.04,0.09) |

P-values in **bold** are significant at the uncorrected level ($P<0.05$).

**Table S13.** Full linear model results for the cortical thickness AIs in all age groups combined.

| Thickness AI | | beta-coefficient | | | Standard Error | | | t-value | | | p-value | | | Cohen's d (95% CI) |
|---|---|---|---|---|---|---|---|---|---|---|---|---|---|---|
| | N cases / controls | diag | age | sex | diag | age | sex | diag | age | sex | diag | age | sex | diag |
| Banks of superior temporal sulcus | 1861/1799 | -0.00261 | -0.00006 | 0.00221 | 0.0013 | 0.00008 | 0.0014 | -2.01 | -0.70 | 1.58 | **0.045** | 0.483 | 0.115 | -0.067 (-0.13,0) |
| Caudal anterior cingulate cortex | 1972/1909 | 0.00249 | -0.00006 | 0.00043 | 0.0017 | 0.00012 | 0.0018 | 1.44 | -0.47 | 0.23 | 0.150 | 0.640 | 0.820 | 0.046 (-0.02,0.11) |
| Caudal middle frontal cortex | 1976/1914 | 0.00187 | -0.00013 | -0.00137 | 0.0008 | 0.00006 | 0.0009 | 2.27 | -2.17 | -1.52 | **0.024** | **0.030** | 0.128 | 0.073 (0.01,0.14) |
| Cuneus | 1971/1915 | 0.00057 | 0.00006 | 0.00018 | 0.0010 | 0.00007 | 0.0011 | 0.53 | 0.90 | 0.16 | 0.593 | 0.368 | 0.877 | 0.017 (-0.05,0.08) |
| Entorhinal cortex | 1862/1806 | -0.00220 | -0.00030 | -0.00487 | 0.0019 | 0.00013 | 0.0021 | -1.11 | -2.40 | -2.26 | 0.265 | **0.016** | **0.024** | -0.037 (-0.1,0.03) |
| Frontal pole | 1976/1916 | 0.00207 | -0.00021 | -0.00328 | 0.0020 | 0.00013 | 0.0022 | 1.01 | -1.63 | -1.48 | 0.313 | 0.103 | 0.139 | 0.033 (-0.03,0.1) |
| Fusiform gyrus | 1923/1865 | -0.00014 | -0.00010 | 0.00007 | 0.0007 | 0.00005 | 0.0007 | -0.19 | -1.86 | 0.09 | 0.847 | 0.063 | 0.925 | -0.006 (-0.07,0.06) |
| Inferior parietal cortex | 1970/1909 | -0.00007 | -0.00001 | 0.00068 | 0.0006 | 0.00004 | 0.0007 | -0.10 | -0.17 | 0.96 | 0.920 | 0.868 | 0.339 | -0.003 (-0.07,0.06) |
| Inferior temporal gyrus | 1887/1852 | -0.00026 | -0.00015 | -0.00003 | 0.0008 | 0.00006 | 0.0009 | -0.30 | -2.33 | -0.03 | 0.766 | **0.020** | 0.977 | -0.01 (-0.07,0.05) |
| Insula | 1963/1902 | -0.00169 | -0.00014 | 0.00032 | 0.0008 | 0.00006 | 0.0009 | -2.05 | -2.49 | 0.36 | **0.040** | **0.013** | 0.719 | -0.066 (-0.13,0) |
| Isthmus cingulate cortex | 1973/1909 | -0.00044 | 0.00011 | -0.00049 | 0.0012 | 0.00006 | 0.0013 | -0.35 | 1.82 | -0.36 | 0.726 | 0.069 | 0.720 | -0.011 (-0.07,0.05) |
| Lateral occipital cortex | 1973/1917 | 0.00071 | 0.00009 | 0.00024 | 0.0007 | 0.00005 | 0.0007 | 0.98 | 1.84 | 0.30 | 0.328 | 0.066 | 0.764 | 0.032 (-0.03,0.09) |
| Lateral orbitofrontal cortex | 1976/1916 | 0.00072 | -0.00002 | -0.00332 | 0.0009 | 0.00006 | 0.0009 | 0.80 | -0.36 | -3.38 | 0.422 | 0.720 | **0.001** | 0.026 (-0.04,0.09) |
| Lingual gyrus | 1929/1867 | -0.00081 | -0.00013 | -0.00114 | 0.0008 | 0.00006 | 0.0009 | -0.95 | -2.16 | -1.23 | 0.340 | 0.031 | 0.220 | -0.031 (-0.09,0.03) |
| Medial orbitofrontal cortex | 1967/1908 | 0.00026 | -0.00003 | -0.00496 | 0.0011 | 0.00009 | 0.0012 | 0.23 | -0.32 | -3.89 | 0.821 | 0.749 | **0.0001** | 0.007 (-0.06,0.07) |
| Middle temporal gyrus | 1826/1784 | -0.00121 | -0.00011 | -0.00155 | 0.0008 | 0.00006 | 0.0009 | -1.45 | -1.92 | -1.70 | 0.147 | 0.056 | 0.090 | -0.049 (-0.11,0.02) |
| Paracentral lobule | 1975/1915 | -0.00074 | -0.00008 | 0.00043 | 0.0008 | 0.00006 | 0.0009 | -0.87 | -1.40 | 0.46 | 0.386 | 0.163 | 0.644 | -0.028 (-0.09,0.03) |
| Parahippocampal gyrus | 1924/1860 | -0.00208 | -0.00004 | -0.00302 | 0.0015 | 0.00009 | 0.0016 | -1.34 | -0.40 | -1.80 | 0.179 | 0.687 | 0.072 | -0.044 (-0.11,0.02) |
| Pars opercularis of inferior frontal gyrus | 1966/1912 | 0.00073 | -0.00022 | 0.00098 | 0.0009 | 0.00006 | 0.0010 | 0.78 | -3.46 | 0.96 | 0.437 | **0.001** | 0.336 | 0.025 (-0.04,0.09) |
| Pars orbitalis of inferior frontal gyrus | 1976/1914 | 0.00170 | 0.00003 | 0.00092 | 0.0014 | 0.00011 | 0.0016 | 1.15 | 0.25 | 0.57 | 0.250 | 0.799 | 0.567 | 0.037 (-0.03,0.1) |
| Pars triangularis of inferior frontal gyrus | 1968/1916 | 0.00037 | -0.00023 | -0.00005 | 0.0010 | 0.00007 | 0.0011 | 0.36 | -3.30 | -0.05 | 0.719 | **0.001** | 0.963 | 0.012 (-0.05,0.07) |
| Pericalcarine cortex | 1971/1914 | 0.00203 | -0.00042 | -0.00014 | 0.0012 | 0.00009 | 0.0013 | 1.61 | -4.88 | -0.10 | 0.108 | $\mathbf{1.1{\times}10^{-6}}$ | 0.917 | 0.052 (-0.01,0.11) |
| Postcentral gyrus | 1951/1890 | -0.00130 | 0.00000 | -0.00147 | 0.0007 | 0.00005 | 0.0007 | -1.77 | -0.08 | -1.86 | 0.077 | 0.937 | 0.063 | -0.057 (-0.12,0.01) |
| Posterior cingulate cortex | 1973/1914 | -0.00099 | -0.00005 | -0.00033 | 0.0010 | 0.00007 | 0.0011 | -0.96 | -0.75 | -0.29 | 0.338 | 0.453 | 0.769 | -0.031 (-0.09,0.03) |
| Precentral gyrus | 1956/1904 | 0.00137 | -0.00007 | -0.00124 | 0.0006 | 0.00004 | 0.0006 | 2.13 | -1.62 | -1.79 | **0.033** | 0.104 | 0.074 | 0.069 (0.01,0.13) |
| Precuneus | 1975/1913 | 0.00049 | 0.00002 | -0.00046 | 0.0006 | 0.00005 | 0.0007 | 0.76 | 0.49 | -0.66 | 0.446 | 0.623 | 0.512 | 0.025 (-0.04,0.09) |
| Rostral anterior cingulate cortex | 1962/1903 | 0.00179 | -0.00046 | 0.00083 | 0.0015 | 0.00011 | 0.0017 | 1.14 | -4.05 | 0.49 | 0.252 | **0.0001** | 0.627 | 0.037 (-0.03,0.1) |
| Rostral middle frontal gyrus | 1976/1913 | -0.00017 | -0.00021 | -0.00039 | 0.0007 | 0.00005 | 0.0007 | -0.23 | -3.92 | -0.50 | 0.817 | **0.0001** | 0.615 | -0.007 (-0.07,0.06) |
| Superior frontal gyrus | 1970/1909 | 0.00066 | -0.00015 | -0.00072 | 0.0005 | 0.00004 | 0.0005 | 1.28 | -3.97 | -1.29 | 0.199 | **0.0001** | 0.199 | 0.041 (-0.02,0.1) |
| Superior parietal cortex | 1969/1914 | -0.00013 | 0.00005 | -0.00067 | 0.0005 | 0.00004 | 0.0006 | -0.23 | 1.38 | -1.07 | 0.821 | 0.167 | 0.286 | -0.007 (-0.07,0.06) |
| Superior temporal gyrus | 1781/1768 | 0.00093 | -0.00008 | 0.00114 | 0.0008 | 0.00005 | 0.0008 | 1.17 | -1.46 | 1.31 | 0.243 | 0.144 | 0.191 | 0.039 (-0.03,0.11) |
| Supramarginal gyrus | 1954/1900 | -0.00108 | 0.00001 | 0.00130 | 0.0007 | 0.00005 | 0.0008 | -1.43 | 0.27 | 1.59 | 0.153 | 0.784 | 0.111 | -0.046 (-0.11,0.02) |
| Temporal pole | 1922/1864 | 0.00070 | -0.00030 | 0.00093 | 0.0018 | 0.00011 | 0.0020 | 0.37 | -2.65 | 0.45 | 0.711 | **0.008** | 0.650 | 0.012 (-0.05,0.08) |
| Transverse temporal gyrus | 1926/1868 | 0.00020 | -0.00011 | 0.00311 | 0.0015 | 0.00009 | 0.0017 | 0.13 | -1.26 | 1.81 | 0.900 | 0.207 | 0.070 | 0.004 (-0.06,0.07) |
| Total average thickness | 1977/1917 | 0.00007 | -0.00006 | -0.00039 | 0.0002 | 0.00002 | 0.0002 | 0.26 | -3.65 | -1.43 | 0.794 | **0.0003** | 0.154 | 0.008 (-0.05,0.07) |

P-values in **bold** are significant at the uncorrected level ($P < 0.05$)

**Table S14** Directions of asymmetry changes in ADHD individuals versus controls for those AIs that had shown nominally significant (P<0.05) associations with diagnosis in any of the main analyses.

| AI region | Mean AI ± SD in controls | Mean AI ± SD in ADHD | Cohen's *d* (95% CI) Left hemisphere | Cohen's *d* (95% CI) Right hemisphere | Controls | ADHD |
|---|---|---|---|---|---|---|
| **Children:** | | | | | | |
| Medial orbitofrontal cortex surface area | -0.0124 ± 0.06 | -0.0035 ± 0.06 | -0.185 (-0.27,-0.1) | -0.326 (-0.42,-0.24) | rightward | decreased |
| Total average surface area | -0.0025 ± 0.01 | -0.0017 ± 0.01 | -0.323 (-0.41,-0.23) | -0.344 (-0.43,-0.26) | rightward | decreased |
| Banks of superior temporal sulcus thickness | -0.0139 ± 0.04 | -0.0176 ± 0.04 | -0.109 (-0.2,-0.02) | 0.008 (-0.08,0.1) | rightward | increased |
| Caudal middle frontal cortex thickness | 0.0028 ± 0.02 | 0.0061 ± 0.03 | -0.022 (-0.11,0.07) | -0.098 (-0.19,-0.01) | leftward | increased |
| Insula thickness | 0.0029 ± 0.03 | 1e-04 ± 0.03 | -0.133 (-0.22,-0.04) | -0.043 (-0.13,0.05) | leftward | decreased |
| Precentral gyrus thickness | 0.0062 ± 0.02 | 0.009 ± 0.02 | -0.116 (-0.2,-0.03) | -0.191 (-0.28,-0.1) | leftward | increased |
| **Adolescents:** | | | | | | |
| Pars orbitalis of inferior frontal gyrus surface area | -0.1113 ± 0.06 | -0.101 ± 0.06 | -0.048 (-0.19,0.1) | -0.223 (-0.37,-0.08) | rightward | decreased |
| Cuneus thickness | -0.0026 ± 0.03 | -0.0076 ± 0.03 | -0.081 (-0.22,0.06) | 0.032 (-0.11,0.18) | rightward | increased |
| **Adults:** | | | | | | |
| Globus Pallidus | 0.0085 ± 0.06 | 0.0031 ± 0.07 | -0.048 (-0.17,0.07) | 0.124 (0.002,0.24) | leftward | decreased |
| Lateral occipital cortex surface area | 0.0125 ± 0.05 | 0.0182 ± 0.05 | 0.021 (-0.09,0.14) | -0.068 (-0.18,0.05) | leftward | increased |
| Lateral occipital cortex thickness | -0.0141 ± 0.02 | -0.012 ± 0.02 | 0.171 (0.05,0.29) | 0.078 (-0.04,0.19) | rightward | decreased |
| Medial orbitofrontal cortex thickness | 0.0128 ± 0.04 | 0.0174 ± 0.04 | -0.019 (-0.14,0.1) | -0.132 (-0.25,-0.02) | leftward | increased |
| Middle temporal gyrus thickness | -0.0035 ± 0.02 | -0.007 ± 0.02 | -0.039 (-0.16,0.08) | 0.05 (-0.07,0.17) | rightward | increased |
| Pericalcarine cortex thickness | -0.0087 ± 0.04 | -0.0025 ± 0.04 | 0.119 (0,0.23) | -0.036 (-0.15,0.08) | rightward | decreased |
| Postcentral gyrus thickness | 0.0065 ± 0.02 | 0.0031 ± 0.02 | -0.008 (-0.12,0.11) | 0.099 (-0.02,0.22) | leftward | decreased |
| **Total:** | | | | | | |
| Globus Pallidus | 0.0231 ± 0.06 | 0.0193 ± 0.06 | -0.127 (-0.19,-0.06) | -0.077 (-0.14,-0.01) | leftward | decreased |
| Medial orbitofrontal cortex surface area | -0.0073 ± 0.06 | -0.0017 ± 0.06 | -0.127 (-0.19,-0.06) | -0.212 (-0.27,-0.15) | rightward | decreased |
| Banks of superior temporal sulcus thickness | -0.0153 ± 0.04 | -0.0183 ± 0.04 | -0.031 (-0.1,0.03) | 0.034 (-0.03,0.1) | rightward | increased |
| Caudal middle frontal cortex thickness | 0.0026 ± 0.03 | 0.0052 ± 0.03 | -0.02 (-0.08,0.04) | -0.078 (-0.14,-0.02) | leftward | increased |
| Insula thickness | 0.0025 ± 0.03 | 4e-04 ± 0.03 | -0.079 (-0.14,-0.02) | -0.019 (-0.08,0.04) | leftward | decreased |
| Precentral gyrus thickness | 0.0057 ± 0.02 | 0.0074 ± 0.02 | -0.091 (-0.15,-0.03) | -0.132 (-0.2,-0.07) | leftward | increased |

The raw means and standard deviations are indicated, as well as the Cohen's *d* effect sizes for left and right hemispheric measures (i.e., when left or right hemispheric measures were analyzed separately as dependent variables). Additionally, the average direction of asymmetry in controls (derived from the raw mean AI) and its change in ADHD is shown. Positive AI values indicate leftward asymmetry, negative AI values indicate rightward asymmetry.

**Table S15.** Sensitivity analyses in  for the effects of diagnosis in all age groups combined, for subcortical volume AIs.

| | Main analysis | | Non-linear age | | Winsorized | |
|---|---|---|---|---|---|---|
| **Subcortical volume AI** | ***P*** | ***d*** | ***P*** | ***d*** | ***P*** | ***d*** |
| Accumbens | 0.289 | -0.037 | 0.290 | -0.037 | 0.279 | -0.037 |
| Amygdala | 0.788 | -0.009 | 0.797 | -0.009 | 0.875 | -0.005 |
| Caudate Nucleus | 0.441 | 0.027 | 0.442 | 0.027 | 0.445 | 0.026 |
| Globus Pallidus | **0.033** | -0.074 | **0.034** | -0.073 | **0.032** | -0.074 |
| Hippocampus | 0.708 | 0.013 | 0.706 | 0.013 | 0.614 | 0.017 |
| Putamen | 0.162 | -0.048 | 0.160 | -0.049 | 0.226 | -0.042 |
| Thalamus[1] | 0.202 | 0.046 | 0.194 | 0.047 | 0.191 | 0.047 |

P-values (P) and Cohen's *d* values (d) for the effects of diagnosis are indicated. P-values in **bold** are significant at the uncorrected level (P < 0.05). [1]Thalamus volume was not available from the NIH dataset

.

**Table S16.** Sensitivity analyses for the effects of diagnosis in all age groups combined, for cortical surface area AIs.

| | Main analysis | | Non-linear age | | Winsorized | |
|---|---|---|---|---|---|---|
| **Cortical surface area AI** | ***P*** | ***d*** | ***P*** | ***d*** | ***P*** | ***d*** |
| banks of superior temporal sulcus | 0.378 | -0.029 | 0.369 | -0.030 | 0.387 | -0.029 |
| caudal anterior cingulate cortex | 0.582 | -0.018 | 0.586 | -0.018 | 0.576 | -0.018 |
| caudal middle frontal cortex | 0.174 | 0.044 | 0.179 | 0.043 | 0.165 | 0.045 |
| cuneus | 0.867 | -0.005 | 0.832 | -0.007 | 0.900 | -0.004 |
| entorhinal cortex | 0.273 | -0.036 | 0.271 | -0.037 | 0.304 | -0.034 |
| frontal pole | 0.117 | -0.051 | 0.118 | -0.050 | 0.111 | -0.051 |
| fusiform gyrus | 0.120 | -0.051 | 0.122 | -0.051 | 0.142 | -0.048 |
| inferior parietal cortex | 0.312 | 0.033 | 0.295 | 0.034 | 0.313 | 0.033 |
| inferior temporal gyrus | 0.193 | 0.043 | 0.182 | 0.044 | 0.185 | 0.044 |
| insula | 0.236 | 0.038 | 0.220 | 0.040 | 0.263 | 0.036 |
| isthmus cingulate cortex | 0.718 | -0.012 | 0.756 | -0.010 | 0.820 | -0.007 |
| lateral occipital cortex | 0.656 | 0.014 | 0.649 | 0.015 | 0.647 | 0.015 |
| lateral orbitofrontal cortex | 0.085 | -0.056 | 0.084 | -0.056 | 0.084 | -0.056 |
| lingual gyrus | 0.886 | 0.005 | 0.898 | 0.004 | 0.705 | 0.012 |
| medial orbitofrontal cortex | **0.031** | 0.070 | **0.032** | 0.070 | **0.029** | 0.071 |
| middle temporal gyrus | 0.443 | 0.026 | 0.451 | 0.025 | 0.424 | 0.027 |
| paracentral lobule | 0.059 | -0.061 | 0.060 | -0.061 | 0.059 | -0.061 |
| parahippocampal gyrus | 0.852 | -0.006 | 0.859 | -0.006 | 0.835 | -0.007 |
| pars opercularis of inferior frontal gyrus | 0.459 | 0.024 | 0.452 | 0.024 | 0.456 | 0.024 |
| pars orbitalis of inferior frontal gyrus | 0.051 | 0.063 | **0.047** | 0.064 | **0.047** | 0.064 |
| pars triangularis of inferior frontal gyrus | 0.220 | 0.040 | 0.209 | 0.041 | 0.198 | 0.042 |
| pericalcarine cortex | 0.836 | 0.007 | 0.860 | 0.006 | 0.705 | 0.012 |
| postcentral gyrus | 0.374 | 0.029 | 0.385 | 0.028 | 0.356 | 0.030 |
| posterior cingulate cortex | 0.876 | -0.005 | 0.880 | -0.005 | 0.862 | -0.006 |
| precentral gyrus | 0.092 | -0.054 | 0.093 | -0.054 | 0.108 | -0.052 |
| precuneus | 0.574 | 0.018 | 0.568 | 0.018 | 0.492 | 0.022 |
| rostral anterior cingulate cortex | 0.462 | -0.024 | 0.487 | -0.022 | 0.456 | -0.024 |
| rostral middle frontal gyrus | 0.210 | -0.040 | 0.227 | -0.039 | 0.228 | -0.039 |
| superior frontal gyrus | 0.544 | 0.020 | 0.583 | 0.018 | 0.483 | 0.023 |
| superiorparietal | 0.338 | 0.031 | 0.314 | 0.033 | 0.286 | 0.034 |
| superior temporal gyrus | 0.994 | 0.000 | 0.978 | -0.001 | 0.810 | 0.008 |
| supramarginal gyrus | 0.279 | -0.035 | 0.275 | -0.035 | 0.313 | -0.033 |
| temporal pole | 0.727 | 0.011 | 0.752 | 0.010 | 0.723 | 0.012 |
| transverse temporal gyrus | 0.980 | -0.001 | 0.966 | -0.001 | 0.920 | 0.003 |
| total average surface area | 0.425 | 0.026 | 0.417 | 0.026 | 0.094 | 0.054 |

P-values (P) and Cohen's *d* values (d) for the effects of diagnosis are indicated. P-values in **bold** are significant at the uncorrected level (P < 0.05).

**Table S17.** Sensitivity analyses for the effects of diagnosis in all age groups combined, for cortical thickness AIs.

| | **Main analysis** | | **Non-linear age** | | **Winsorized** | |
|---|---|---|---|---|---|---|
| **Cortical thickness AI** | ***P*** | ***d*** | ***P*** | ***d*** | ***P*** | ***d*** |
| banks of superior temporal sulcus | **0.045** | -0.067 | **0.045** | -0.067 | **0.046** | -0.066 |
| caudal anterior cingulate cortex | 0.150 | 0.046 | 0.148 | 0.047 | 0.147 | 0.047 |
| caudal middle frontal cortex | **0.024** | 0.073 | **0.023** | 0.073 | **0.023** | 0.073 |
| cuneus | 0.593 | 0.017 | 0.589 | 0.017 | 0.596 | 0.017 |
| entorhinal cortex | 0.265 | -0.037 | 0.266 | -0.037 | 0.277 | -0.036 |
| frontal pole | 0.313 | 0.033 | 0.312 | 0.033 | 0.306 | 0.033 |
| fusiform gyrus | 0.847 | -0.006 | 0.842 | -0.007 | 0.880 | -0.005 |
| inferior parietal cortex | 0.920 | -0.003 | 0.923 | -0.003 | 0.917 | -0.003 |
| inferior temporal gyrus | 0.766 | -0.010 | 0.758 | -0.010 | 0.768 | -0.010 |
| insula | **0.040** | -0.066 | **0.041** | -0.066 | 0.050 | -0.063 |
| isthmus cingulate cortex | 0.726 | -0.011 | 0.682 | -0.013 | 0.730 | -0.011 |
| lateral occipital cortex | 0.328 | 0.032 | 0.335 | 0.031 | 0.327 | 0.032 |
| lateral orbitofrontal cortex | 0.422 | 0.026 | 0.419 | 0.026 | 0.434 | 0.025 |
| lingual gyrus | 0.340 | -0.031 | 0.332 | -0.032 | 0.360 | -0.030 |
| medial orbitofrontal cortex | 0.821 | 0.007 | 0.821 | 0.007 | 0.809 | 0.008 |
| middle temporal gyrus | 0.147 | -0.049 | 0.151 | -0.048 | 0.171 | -0.046 |
| paracentral lobule | 0.386 | -0.028 | 0.398 | -0.027 | 0.386 | -0.028 |
| parahippocampal gyrus | 0.179 | -0.044 | 0.173 | -0.045 | 0.178 | -0.044 |
| pars opercularis of inferior frontal gyrus | 0.437 | 0.025 | 0.420 | 0.026 | 0.414 | 0.026 |
| pars orbitalis of inferior frontal gyrus | 0.250 | 0.037 | 0.242 | 0.038 | 0.252 | 0.037 |
| pars triangularis of inferior frontal gyrus | 0.719 | 0.012 | 0.697 | 0.013 | 0.640 | 0.015 |
| pericalcarine cortex | 0.108 | 0.052 | 0.102 | 0.053 | 0.106 | 0.052 |
| postcentral gyrus | 0.077 | -0.057 | 0.075 | -0.058 | 0.078 | -0.057 |
| posterior cingulate cortex | 0.338 | -0.031 | 0.331 | -0.031 | 0.358 | -0.030 |
| precentral gyrus | **0.033** | 0.069 | **0.032** | 0.069 | **0.031** | 0.070 |
| precuneus | 0.446 | 0.025 | 0.454 | 0.024 | 0.452 | 0.024 |
| rostral anterior cingulate cortex | 0.252 | 0.037 | 0.248 | 0.037 | 0.260 | 0.036 |
| rostral middle frontal gyrus | 0.817 | -0.007 | 0.825 | -0.007 | 0.805 | -0.008 |
| superior frontal gyrus | 0.199 | 0.041 | 0.193 | 0.042 | 0.234 | 0.038 |
| superiorparietal | 0.821 | -0.007 | 0.823 | -0.007 | 0.833 | -0.007 |
| superior temporal gyrus | 0.243 | 0.039 | 0.243 | 0.039 | 0.197 | 0.044 |
| supramarginal gyrus | 0.153 | -0.046 | 0.153 | -0.046 | 0.173 | -0.044 |
| temporal pole | 0.711 | 0.012 | 0.683 | 0.013 | 0.691 | 0.013 |
| transverse temporal gyrus | 0.900 | 0.004 | 0.920 | 0.003 | 0.872 | 0.005 |
| total average thickness | 0.794 | 0.008 | 0.791 | 0.009 | 0.830 | 0.007 |

P-values (P) and Cohen's *d* values (d) for the effects of diagnosis are indicated. P-values in **bold** are significant at the uncorrected level ($P < 0.05$).

**Table S18**. Associations of subcortical volume AIs with IQ in all age groups combined.

| Region | IQ cases | | | IQ controls | | |
|---|---|---|---|---|---|---|
| | **N subjects** | **p-value** | **t-value** | **N subjects** | **p-value** | **t-value** |
| Accumbens | 1553 | **0.02** | 2.27 | 1474 | 0.83 | 0.22 |
| Amygdala | 1552 | 0.26 | 1.13 | 1473 | 0.10 | 1.63 |
| Caudate Nucleus | 1553 | 0.42 | 0.81 | 1474 | 0.77 | -0.29 |
| Globus Pallidus | 1555 | 0.63 | 0.48 | 1470 | **0.04** | -2.05 |
| Hippocampus | 1552 | **0.05** | -1.97 | 1470 | 0.51 | 0.66 |
| Putamen | 1554 | 0.33 | 0.98 | 1464 | 0.28 | 1.09 |
| Thalamus[1] | 1450 | 0.31 | -1.01 | 1361 | 0.51 | -0.66 |

P-values (P) and t-values (t) for the effects of IQ in cases and controls are indicated. P-values in **bold** are significant at the uncorrected level ($P < 0.05$). [1]Thalamus volume was not available from the NIH dataset.

**Table S19**. Associations of cortical surface area AIs with IQ in all age groups combined.

| Region | IQ cases | | | IQ controls | | |
|---|---|---|---|---|---|---|
| | N subjects | p-value | t-value | N subjects | p-value | t-value |
| banks of superior temporal sulcus | 1674 | 0.83 | 0.21 | 1614 | 0.35 | 0.94 |
| caudal anterior cingulate cortex | 1783 | 0.49 | 0.69 | 1719 | 1.00 | 0.01 |
| caudal middle frontal cortex | 1788 | 0.11 | 1.62 | 1723 | 0.05 | -1.93 |
| cuneus | 1781 | 0.69 | -0.41 | 1724 | 0.12 | 1.54 |
| entorhinal cortex | 1679 | 0.43 | 0.79 | 1618 | 0.52 | 0.64 |
| frontal pole | 1788 | 0.69 | 0.39 | 1726 | 0.72 | 0.36 |
| fusiform gyrus | 1733 | 0.96 | 0.05 | 1675 | 0.33 | -0.97 |
| inferior parietal cortex | 1780 | 0.38 | 0.88 | 1716 | 0.74 | -0.33 |
| inferior temporal gyrus | 1699 | 0.62 | -0.50 | 1663 | 0.60 | -0.53 |
| insula | 1777 | 0.99 | -0.02 | 1713 | 0.76 | -0.31 |
| isthmus cingulate cortex | 1786 | 0.52 | 0.64 | 1719 | 0.39 | 0.86 |
| lateral occipital cortex | 1784 | 0.05 | -1.93 | 1726 | 0.71 | -0.37 |
| lateral orbitofrontal cortex | 1787 | 0.19 | -1.33 | 1726 | 0.25 | -1.16 |
| lingual gyrus | 1740 | 0.66 | -0.43 | 1678 | 0.41 | 0.83 |
| medial orbitofrontal cortex | 1777 | 0.37 | 0.90 | 1717 | 0.50 | -0.68 |
| middle temporal gyrus | 1637 | 0.25 | 1.16 | 1594 | **0.02** | -2.43 |
| paracentral lobule | 1786 | 0.49 | -0.70 | 1725 | 0.48 | -0.71 |
| parahippocampal gyrus | 1735 | 0.19 | -1.30 | 1669 | 0.08 | -1.76 |
| pars opercularis of inferior frontal gyrus | 1778 | 0.63 | -0.48 | 1722 | 0.22 | 1.24 |
| pars orbitalis of inferior frontal gyrus | 1787 | 0.72 | -0.36 | 1724 | 0.65 | 0.46 |
| pars triangularis of inferior frontal gyrus | 1782 | 0.28 | 1.08 | 1726 | 0.07 | 1.84 |
| pericalcarine cortex | 1784 | 0.07 | -1.81 | 1725 | 0.48 | 0.71 |
| postcentral gyrus | 1766 | 0.76 | -0.30 | 1699 | 0.97 | 0.04 |
| posterior cingulate cortex | 1785 | 0.64 | 0.47 | 1721 | 0.79 | -0.27 |
| precentral gyrus | 1770 | 0.14 | 1.46 | 1716 | 0.80 | 0.26 |
| precuneus | 1786 | 0.55 | -0.60 | 1723 | 0.98 | -0.02 |
| rostral anterior cingulate cortex | 1773 | 0.88 | -0.15 | 1715 | 0.87 | 0.16 |
| rostral middle frontal gyrus | 1787 | 0.32 | -0.99 | 1722 | 0.76 | -0.30 |
| superior frontal gyrus | 1781 | 0.07 | -1.83 | 1719 | 0.74 | -0.33 |
| superiorparietal | 1780 | 0.62 | 0.50 | 1724 | 0.89 | -0.13 |
| superior temporal gyrus | 1594 | 0.53 | -0.63 | 1577 | 0.49 | 0.69 |
| supramarginal gyrus | 1765 | 0.19 | 1.30 | 1708 | 0.97 | 0.03 |
| temporal pole | 1735 | 0.99 | 0.02 | 1675 | 0.58 | -0.55 |
| transverse temporal gyrus | 1738 | 0.99 | -0.01 | 1678 | 0.53 | 0.63 |
| total average surface area | 1788 | 0.55 | -0.59 | 1727 | 0.48 | -0.70 |

P-values (P) and t-values (t) for the effects of IQ in cases and controls are indicated. P-values in **bold** are significant at the uncorrected level ($P < 0.05$).

**Table S20**. Associations of cortical thickness AIs with IQ in all age groups combined.

| Region | IQ cases | | | IQ controls | | |
|---|---|---|---|---|---|---|
| | **N subjects** | **p-value** | **t-value** | **N subjects** | **p-value** | **t-value** |
| banks of superior temporal sulcus | 1673 | 0.66 | 0.44 | 1614 | 0.45 | -0.76 |
| caudal anterior cingulate cortex | 1782 | 0.36 | 0.91 | 1719 | 0.06 | 1.90 |
| caudal middle frontal cortex | 1786 | 0.15 | -1.45 | 1724 | 0.67 | 0.43 |
| cuneus | 1781 | 0.89 | -0.13 | 1725 | 0.92 | 0.10 |
| entorhinal cortex | 1679 | 0.42 | 0.80 | 1619 | 0.40 | 0.84 |
| frontal pole | 1786 | 0.54 | 0.62 | 1726 | 0.14 | 1.46 |
| fusiform gyrus | 1734 | 0.49 | 0.69 | 1675 | 0.07 | 1.78 |
| inferior parietal cortex | 1780 | 0.72 | -0.35 | 1720 | 0.32 | -1.00 |
| inferior temporal gyrus | 1698 | 0.39 | -0.85 | 1662 | 0.13 | 1.52 |
| insula | 1776 | 0.04 | 2.04 | 1714 | 0.92 | -0.10 |
| isthmus cingulate cortex | 1783 | 0.66 | -0.44 | 1719 | 0.94 | 0.07 |
| lateral occipital cortex | 1783 | 0.17 | 1.37 | 1727 | 0.68 | -0.41 |
| lateral orbitofrontal cortex | 1786 | 0.96 | -0.05 | 1726 | 0.86 | -0.18 |
| lingual gyrus | 1739 | 0.51 | 0.66 | 1678 | 0.48 | 0.71 |
| medial orbitofrontal cortex | 1777 | 0.66 | 0.44 | 1718 | 0.25 | 1.15 |
| middle temporal gyrus | 1637 | 0.87 | 0.16 | 1596 | 0.10 | 1.64 |
| paracentral lobule | 1785 | 0.45 | -0.76 | 1725 | 0.23 | -1.21 |
| parahippocampal gyrus | 1735 | 0.92 | 0.10 | 1670 | 0.73 | 0.35 |
| pars opercularis of inferior frontal gyrus | 1777 | 0.46 | 0.73 | 1722 | 0.11 | 1.60 |
| pars orbitalis of inferior frontal gyrus | 1786 | 0.71 | 0.37 | 1724 | 0.37 | -0.89 |
| pars triangularis of inferior frontal gyrus | 1779 | 0.32 | 1.00 | 1726 | 0.64 | 0.47 |
| pericalcarine cortex | 1781 | 0.37 | 0.90 | 1724 | 0.83 | -0.22 |
| postcentral gyrus | 1763 | 0.19 | 1.31 | 1701 | 0.71 | 0.37 |
| posterior cingulate cortex | 1783 | 0.97 | 0.04 | 1724 | 0.57 | -0.57 |
| precentral gyrus | 1768 | 0.20 | -1.28 | 1714 | 0.67 | -0.43 |
| precuneus | 1785 | 0.73 | -0.34 | 1723 | 0.38 | -0.88 |
| rostral anterior cingulate cortex | 1772 | 0.39 | 0.87 | 1713 | **0.02** | 2.31 |
| rostral middle frontal gyrus | 1786 | 0.91 | -0.11 | 1723 | 0.54 | 0.61 |
| superior frontal gyrus | 1780 | 0.97 | 0.04 | 1719 | 0.89 | -0.13 |
| superiorparietal | 1779 | 0.18 | 1.35 | 1724 | 0.49 | 0.69 |
| superior temporal gyrus | 1594 | 0.24 | -1.18 | 1580 | 0.69 | -0.41 |
| supramarginal gyrus | 1764 | 0.68 | 0.42 | 1711 | **0.03** | -2.23 |
| temporal pole | 1734 | 0.25 | -1.16 | 1674 | 0.24 | 1.18 |
| transverse temporal gyrus | 1737 | 0.38 | 0.88 | 1678 | 0.72 | 0.36 |
| total average thickness | 1787 | 0.37 | 0.89 | 1727 | 0.60 | 0.53 |

P-values (P) and t-values (t) for the effects of IQ in cases and controls are indicated. P-values in **bold** are significant at the uncorrected level (P < 0.05).

**Table S21** Associations of subcortical volume AIs with comorbidities in ADHD individuals, all age groups combined.

| | Mood | | | | ODD | | | | Anxiety | | | | SUD | | | |
|---|---|---|---|---|---|---|---|---|---|---|---|---|---|---|---|---|
| | N subj (no/yes) | N dtst | p-value | t-value | N subj (no/yes) | N dtst | p-value | t-value | N subj (no/yes) | N dtst | p-value | t-value | N subj (no/yes) | N dtst | p-value | t-value |
| Accumbens | 268/164 | 6 | 0.58 | 0.55 | 81/39 | 4 | 0.25 | 1.16 | 388/66 | 7 | 0.66 | 0.43 | 252/67 | 5 | 0.49 | 0.69 |
| Amygdala | 266/164 | 6 | 0.75 | 0.32 | 81/39 | 4 | 0.32 | -1.01 | 386/66 | 7 | 0.28 | 1.09 | 252/67 | 5 | 0.33 | -0.99 |
| Caudate Nucleus | 268/164 | 6 | 0.48 | 0.71 | 81/39 | 4 | 0.25 | -1.16 | 388/66 | 7 | 0.17 | -1.36 | 252/65 | 5 | 0.11 | -1.62 |
| Globus Pallidus | 268/164 | 6 | 0.67 | 0.42 | 81/39 | 4 | 0.61 | 0.52 | 388/66 | 7 | 0.98 | -0.03 | 252/67 | 5 | 0.58 | 0.55 |
| Hippocampus | 267/164 | 6 | 0.94 | 0.08 | 81/39 | 4 | 0.31 | -1.03 | 387/66 | 7 | 0.92 | -0.10 | 252/67 | 5 | 0.41 | -0.82 |
| Putamen | 268/164 | 6 | 0.11 | -1.61 | 81/39 | 4 | 0.62 | -0.50 | 388/66 | 7 | 0.67 | -0.43 | 252/66 | 5 | 0.66 | -0.44 |
| Thalamus[1] | 266/164 | 6 | 0.81 | -0.24 | 81/39 | 4 | 0.42 | -0.81 | 386/66 | 7 | 0.49 | 0.70 | 251/65 | 5 | 0.55 | 0.60 |

P-values (P) and t-values (t) for the effects of ADHD comorbidities are indicated. P-values in **bold** are significant at the uncorrected level (P < 0.05). [1]Thalamus volume was not available from the NIH dataset. subj=subjects; dtst=dataset

**Table S22**. Associations of cortical surface area AIs with comorbidities in ADHD individuals, all age groups combined.

| | | | Mood | | | ODD | | | | Anxiety | | | SUD | | | |
|---|---|---|---|---|---|---|---|---|---|---|---|---|---|---|---|---|
| | N subj (no/yes) | N dtst | p-value | t-value | N subj (no/yes) | N dtst | p-value | t-value | N subj (no/yes) | N dtst | p-value | t-value | N subj (no/yes) | N dtst | p-value | t-value |
| banks of superior temporal sulcus | 378/174 | 8 | 0.42 | 0.80 | 145/78 | 6 | 0.41 | -0.82 | 493/81 | 9 | 0.79 | 0.27 | 331/75 | 6 | 0.11 | 1.60 |
| caudal anterior cingulate cortex | 383/178 | 8 | 0.15 | -1.43 | 151/78 | 6 | 0.30 | -1.03 | 501/82 | 9 | 0.30 | -1.04 | 333/77 | 6 | 0.80 | -0.26 |
| caudal middle frontal cortex | 384/179 | 8 | 0.81 | -0.24 | 151/80 | 6 | 0.45 | 0.75 | 503/82 | 9 | 0.08 | 1.76 | 335/77 | 6 | 0.65 | 0.46 |
| cuneus | 383/178 | 8 | 0.09 | 1.72 | 149/79 | 6 | 0.85 | 0.19 | 501/82 | 9 | 0.21 | 1.25 | 335/76 | 6 | 0.86 | 0.18 |
| entorhinal cortex | 344/157 | 8 | 0.13 | 1.53 | 131/76 | 6 | 0.53 | -0.63 | 443/77 | 9 | 0.12 | 1.58 | 295/69 | 6 | 0.34 | 0.97 |
| frontal pole | 384/179 | 8 | 0.94 | 0.08 | 151/80 | 6 | 0.47 | -0.73 | 503/82 | 9 | 0.26 | 1.14 | 335/77 | 6 | 0.79 | 0.26 |
| fusiform gyrus | 355/158 | 8 | 0.70 | -0.39 | 148/78 | 6 | 0.79 | -0.26 | 455/77 | 9 | 0.15 | 1.44 | 294/70 | 6 | 0.41 | -0.82 |
| inferior parietal cortex | 383/176 | 8 | 0.74 | -0.33 | 150/80 | 6 | 0.14 | -1.48 | 500/81 | 9 | 0.93 | -0.09 | 334/74 | 6 | 0.77 | -0.30 |
| inferior temporal gyrus | 357/159 | 8 | 0.05 | 2.00 | 150/77 | 6 | 0.07 | 1.79 | 458/77 | 9 | 0.48 | 0.70 | 295/70 | 6 | 0.88 | 0.15 |
| insula | 377/176 | 8 | 0.88 | 0.15 | 149/78 | 6 | 0.79 | -0.27 | 494/81 | 9 | 0.69 | -0.39 | 335/76 | 6 | 0.72 | 0.36 |
| isthmus cingulate cortex | 384/179 | 8 | 0.26 | -1.12 | 151/79 | 6 | 0.69 | -0.40 | 503/82 | 9 | 0.69 | 0.40 | 335/77 | 6 | 0.32 | 1.00 |
| lateral occipital cortex | 383/177 | 8 | 0.86 | -0.17 | 151/80 | 6 | 0.49 | -0.69 | 500/82 | 9 | 0.30 | -1.05 | 335/74 | 6 | 0.50 | 0.67 |
| lateral orbitofrontal cortex | 383/179 | 8 | 0.18 | 1.33 | 150/80 | 6 | 0.63 | 0.49 | 502/82 | 9 | 0.97 | 0.04 | 334/77 | 6 | 0.71 | 0.37 |
| lingual gyrus | 356/159 | 8 | 0.44 | -0.78 | 151/80 | 6 | 0.15 | -1.43 | 458/79 | 9 | 0.23 | -1.21 | 295/69 | 6 | 0.53 | 0.63 |
| medial orbitofrontal cortex | 382/179 | 8 | 0.56 | -0.58 | 149/79 | 6 | 0.78 | -0.28 | 501/82 | 9 | 0.93 | -0.08 | 335/76 | 6 | 0.13 | 1.53 |
| middle temporal gyrus | 353/155 | 8 | 0.71 | 0.37 | 147/74 | 6 | 0.75 | -0.32 | 451/76 | 9 | 0.73 | -0.35 | 290/68 | 6 | 0.99 | 0.01 |
| paracentral lobule | 383/179 | 8 | 0.33 | -0.98 | 151/80 | 6 | 0.07 | 1.82 | 503/81 | 9 | 0.99 | -0.01 | 334/77 | 6 | 0.06 | 1.88 |
| parahippocampal gyrus | 357/158 | 8 | 0.32 | -1.00 | 149/77 | 6 | 0.36 | 0.92 | 458/76 | 9 | 0.11 | 1.61 | 295/70 | 6 | 0.76 | -0.30 |
| pars opercularis of inferior frontal gyrus | 381/178 | 8 | 0.40 | 0.84 | 145/79 | 6 | 0.93 | -0.08 | 499/82 | 9 | 0.57 | 0.57 | 331/77 | 6 | 0.67 | -0.42 |
| pars orbitalis of inferior frontal gyrus | 383/179 | 8 | 0.85 | -0.20 | 151/80 | 6 | 0.84 | 0.20 | 502/82 | 9 | 0.18 | 1.35 | 334/77 | 6 | 0.45 | 0.76 |
| pars triangularis of inferior frontal gyrus | 384/179 | 8 | 0.83 | -0.22 | 147/79 | 6 | 0.15 | -1.44 | 503/82 | 9 | 0.85 | -0.19 | 335/77 | 6 | 0.32 | 0.99 |
| pericalcarine cortex | 382/179 | 8 | 0.19 | 1.30 | 148/80 | 6 | 0.48 | -0.71 | 501/82 | 9 | 0.96 | 0.05 | 334/76 | 6 | 0.94 | 0.07 |
| postcentral gyrus | 379/176 | 8 | 0.68 | 0.41 | 149/75 | 6 | 0.97 | -0.04 | 496/81 | 9 | 0.72 | -0.35 | 334/77 | 6 | **0.01** | 2.78 |
| posterior cingulate cortex | 384/179 | 8 | 0.36 | 0.92 | 149/80 | 6 | 0.32 | -0.99 | 503/82 | 9 | 0.17 | -1.39 | 335/77 | 6 | 0.27 | -1.10 |
| precentral gyrus | 379/177 | 8 | 0.50 | 0.68 | 146/78 | 6 | 0.81 | 0.24 | 497/81 | 9 | 0.36 | -0.92 | 331/77 | 6 | 0.95 | -0.07 |
| precuneus | 383/179 | 8 | 0.90 | 0.13 | 151/80 | 6 | 0.75 | 0.31 | 502/82 | 9 | 0.97 | -0.03 | 335/76 | 6 | 0.83 | -0.21 |

| | | | | | | | | | | | | | | | | |
|---|---|---|---|---|---|---|---|---|---|---|---|---|---|---|---|---|
| rostral anterior cingulate cortex | 381/177 | 8 | **0.03** | -2.19 | 148/78 | 6 | 0.51 | -0.66 | 500/80 | 9 | 0.39 | -0.86 | 332/75 | 6 | 0.63 | -0.49 |
| rostral middle frontal gyrus | 384/179 | 8 | 0.58 | 0.55 | 150/80 | 6 | 0.19 | -1.30 | 503/82 | 9 | 0.88 | 0.15 | 335/77 | 6 | 0.05 | -1.94 |
| superior frontal gyrus | 383/178 | 8 | 0.99 | -0.01 | 150/79 | 6 | 0.07 | 1.85 | 501/82 | 9 | 0.25 | -1.15 | 333/77 | 6 | 0.45 | 0.76 |
| superiorparietal | 382/177 | 8 | 0.87 | -0.17 | 150/79 | 6 | 0.58 | 0.55 | 500/81 | 9 | 0.78 | 0.27 | 334/74 | 6 | 0.44 | -0.77 |
| superior temporal gyrus | 352/151 | 8 | 0.20 | -1.30 | 147/76 | 6 | 0.52 | -0.64 | 446/76 | 9 | 0.68 | 0.41 | 288/67 | 6 | 0.37 | -0.89 |
| supramarginal gyrus | 382/176 | 8 | 0.62 | -0.50 | 151/79 | 6 | 0.28 | -1.08 | 498/82 | 9 | 0.17 | 1.39 | 334/76 | 6 | **0.03** | -2.14 |
| temporal pole | 357/159 | 8 | 0.98 | 0.02 | 150/77 | 6 | **0.04** | 2.10 | 458/77 | 9 | 0.82 | 0.23 | 295/70 | 6 | 0.21 | 1.27 |
| transverse temporal gyrus | 357/159 | 8 | 0.63 | -0.48 | 150/78 | 6 | 0.67 | -0.42 | 458/77 | 9 | 0.66 | 0.44 | 295/70 | 6 | 0.23 | -1.20 |
| total average surface area | 384/179 | 8 | 0.87 | 0.17 | 151/80 | 6 | 0.69 | -0.40 | 503/82 | 9 | 0.60 | 0.52 | 335/77 | 6 | 0.37 | 0.89 |

P-values (P) and t-values (t) for the effects of ADHD comorbidities are indicated. P-values in **bold** are significant at the uncorrected level (P < 0.05). subj=subjects; dtst=dataset

**Table S23**. Associations of cortical thickness AIs with comorbidities in ADHD individuals, all age groups combined.

| | Mood | | | | ODD | | | | Anxiety | | | | SUD | | | |
|---|---|---|---|---|---|---|---|---|---|---|---|---|---|---|---|---|
| | N subj (no/yes) | N dtst | p-value | t-value | N subj (no/yes) | N dtst | p-value | t-value | N subj (no/yes) | N dtst | p-value | t-value | N subj (no/yes) | N dtst | p-value | t-value |
| banks of superior temporal sulcus | 378/174 | 8 | 0.63 | -0.48 | 145/78 | 6 | 0.22 | 1.23 | 493/81 | 9 | 0.36 | -0.91 | 331/75 | 6 | 0.41 | 0.83 |
| caudal anterior cingulate cortex | 383/178 | 8 | 0.58 | -0.55 | 151/78 | 6 | 0.56 | 0.59 | 501/82 | 9 | 0.55 | 0.59 | 333/77 | 6 | 0.30 | -1.04 |
| caudal middle frontal cortex | 383/179 | 8 | 0.29 | 1.05 | 151/79 | 6 | 0.19 | 1.31 | 502/82 | 9 | 0.39 | 0.85 | 335/77 | 6 | 0.92 | -0.09 |
| cuneus | 383/179 | 8 | 0.98 | 0.03 | 149/79 | 6 | 0.74 | 0.33 | 502/82 | 9 | 0.02 | 2.36 | 335/76 | 6 | 0.01 | 2.70 |
| entorhinal cortex | 345/157 | 8 | **0.004** | -2.89 | 132/76 | 6 | 0.53 | -0.63 | 444/77 | 9 | 0.79 | -0.26 | 295/69 | 6 | 0.95 | -0.06 |
| frontal pole | 384/179 | 8 | 0.77 | -0.29 | 151/80 | 6 | 0.58 | 0.56 | 503/82 | 9 | 0.11 | -1.62 | 335/77 | 6 | 0.16 | 1.39 |
| fusiform gyrus | 357/158 | 8 | 0.55 | 0.59 | 148/78 | 6 | 0.21 | -1.26 | 457/77 | 9 | 0.24 | 1.17 | 294/70 | 6 | 0.69 | 0.40 |
| inferior parietal cortex | 383/176 | 8 | 0.20 | 1.30 | 150/80 | 6 | 0.96 | 0.05 | 500/81 | 9 | 0.70 | -0.38 | 334/74 | 6 | 0.51 | -0.65 |
| inferior temporal gyrus | 357/159 | 8 | 0.90 | 0.12 | 150/77 | 6 | 0.67 | -0.43 | 458/77 | 9 | 0.43 | -0.78 | 295/70 | 6 | 0.54 | -0.62 |
| insula | 377/176 | 8 | 0.49 | -0.70 | 149/78 | 6 | 0.21 | -1.26 | 494/81 | 9 | 0.99 | -0.01 | 335/76 | 6 | 0.44 | -0.78 |
| isthmus cingulate cortex | 384/179 | 8 | 0.93 | 0.09 | 151/79 | 6 | 0.74 | 0.34 | 503/82 | 9 | 0.61 | -0.52 | 335/77 | 6 | 0.75 | -0.32 |
| lateral occipital cortex | 383/177 | 8 | 0.44 | -0.77 | 151/80 | 6 | 0.66 | -0.44 | 500/82 | 9 | **0.02** | -2.41 | 335/74 | 6 | 0.31 | 1.02 |
| lateral orbitofrontal cortex | 383/179 | 8 | 0.76 | 0.31 | 150/80 | 6 | 0.29 | -1.05 | 502/82 | 9 | 0.48 | 0.70 | 334/77 | 6 | 0.23 | 1.19 |
| lingual gyrus | 356/159 | 8 | 0.30 | 1.05 | 151/80 | 6 | 0.17 | -1.38 | 458/79 | 9 | 0.25 | -1.15 | 295/69 | 6 | 0.73 | 0.34 |
| medial orbitofrontal cortex | 383/179 | 8 | 0.50 | 0.67 | 150/79 | 6 | 0.03 | -2.20 | 502/82 | 9 | 0.06 | -1.91 | 335/76 | 6 | 0.52 | -0.64 |
| middle temporal gyrus | 354/155 | 8 | 0.35 | 0.95 | 147/74 | 6 | 0.92 | -0.10 | 452/76 | 9 | 0.41 | -0.83 | 290/68 | 6 | 0.78 | -0.28 |
| paracentral lobule | 383/179 | 8 | 0.49 | -0.68 | 151/80 | 6 | 0.79 | 0.26 | 503/81 | 9 | 0.06 | 1.88 | 334/77 | 6 | 0.05 | 1.99 |
| parahippocampal gyrus | 356/159 | 8 | 0.19 | 1.31 | 149/78 | 6 | 0.55 | 0.60 | 458/77 | 9 | 0.16 | 1.41 | 295/70 | 6 | 0.26 | 1.13 |
| pars opercularis of inferior frontal gyrus | 381/178 | 8 | 0.81 | -0.25 | 145/79 | 6 | 0.81 | 0.24 | 499/82 | 9 | 0.96 | -0.04 | 331/77 | 6 | 0.13 | 1.51 |
| pars orbitalis of inferior frontal gyrus | 383/179 | 8 | 0.51 | 0.66 | 151/80 | 6 | 0.43 | 0.78 | 502/82 | 9 | 0.95 | -0.06 | 334/77 | 6 | 0.31 | 1.02 |
| pars triangularis of inferior frontal gyrus | 383/178 | 8 | **0.03** | 2.22 | 147/77 | 6 | 0.35 | 0.93 | 501/82 | 9 | 0.40 | 0.84 | 335/77 | 6 | 0.39 | -0.86 |
| pericalcarine cortex | 382/179 | 8 | 0.05 | 2.00 | 148/80 | 6 | 0.63 | -0.48 | 501/82 | 9 | 0.99 | 0.01 | 334/76 | 6 | 0.43 | -0.79 |
| postcentral gyrus | 379/176 | 8 | 0.41 | -0.83 | 149/75 | 6 | 0.23 | 1.19 | 496/81 | 9 | 0.76 | 0.30 | 334/77 | 6 | 0.05 | 1.94 |
| posterior cingulate cortex | 383/179 | 8 | 0.08 | 1.78 | 149/79 | 6 | 0.10 | 1.65 | 502/82 | 9 | 0.93 | 0.09 | 335/77 | 6 | 0.60 | 0.52 |
| precentral gyrus | 379/177 | 8 | 0.74 | 0.33 | 146/78 | 6 | 0.56 | 0.58 | 497/81 | 9 | 0.90 | 0.12 | 331/77 | 6 | 0.46 | -0.73 |
| precuneus | 383/179 | 8 | **0.02** | -2.26 | 151/80 | 6 | 0.62 | 0.50 | 502/82 | 9 | 0.24 | -1.19 | 335/76 | 6 | 0.09 | 1.70 |
| rostral anterior cingulate cortex | 381/177 | 8 | 0.05 | 1.96 | 148/78 | 6 | 0.63 | -0.49 | 500/80 | 9 | 0.48 | -0.71 | 332/75 | 6 | 0.12 | -1.57 |

| | | | | | | | | | | | | | | | | |
|---|---|---|---|---|---|---|---|---|---|---|---|---|---|---|---|---|
| rostral middle frontal gyrus | 384/179 | 8 | ***0.0002*** | 3.70 | 150/80 | 6 | 0.36 | 0.91 | 503/82 | 9 | 0.78 | 0.28 | 335/77 | 6 | 0.37 | 0.89 |
| superior frontal gyrus | 383/178 | 8 | 0.99 | -0.02 | 150/79 | 6 | 0.68 | -0.41 | 501/82 | 9 | 0.84 | 0.20 | 333/77 | 6 | 0.59 | -0.55 |
| superiorparietal | 382/177 | 8 | 0.58 | 0.56 | 150/79 | 6 | 0.08 | 1.74 | 500/81 | 9 | 0.73 | -0.35 | 334/74 | 6 | 0.38 | 0.88 |
| superior temporal gyrus | 352/151 | 8 | 0.85 | 0.19 | 147/76 | 6 | 0.33 | 0.98 | 446/76 | 9 | 0.18 | -1.33 | 288/67 | 6 | 0.63 | 0.49 |
| supramarginal gyrus | 382/176 | 8 | 0.68 | 0.41 | 151/79 | 6 | 0.23 | 1.20 | 498/82 | 9 | 0.12 | 1.57 | 334/76 | 6 | 0.40 | -0.84 |
| temporal pole | 357/159 | 8 | 0.92 | -0.10 | 150/77 | 6 | 0.26 | -1.12 | 458/77 | 9 | 0.20 | -1.29 | 295/70 | 6 | 0.71 | -0.37 |
| transverse temporal gyrus | 357/159 | 8 | **0.04** | 2.10 | 150/78 | 6 | 0.75 | 0.32 | 458/77 | 9 | 0.98 | -0.03 | 295/70 | 6 | 0.38 | 0.89 |
| total average thickness | 384/179 | 8 | 0.11 | 1.61 | 151/80 | 6 | 0.67 | 0.42 | 503/82 | 9 | 0.60 | -0.52 | 335/77 | 6 | 0.64 | 0.46 |

P-values (P) and t-values (t) for the effects of ADHD comorbidities are indicated. P-values in **bold** are significant at the uncorrected level (P < 0.05), and those in ***bold italic*** survived FDR < 0.05. subj=subjects; dtst=dataset.

**Table S24**. Associations of subcortical volume AIs with disorder severity in ADHD individuals, all age groups combined.

| Region | Hyperactivity/impulsivity | | | Inattention | | |
|---|---|---|---|---|---|---|
| | N subjects (no/yes) | p-value | t-value | N subjects (no/yes) | p-value | t-value |
| Accumbens | 281 | 0.54 | 0.61 | 281 | 0.36 | 0.91 |
| Amygdala | 282 | 0.83 | 0.21 | 282 | 0.93 | 0.09 |
| Caudate Nucleus | 281 | 0.31 | 1.03 | 281 | 0.83 | 0.22 |
| Globus Pallidus | 283 | 0.96 | 0.05 | 283 | 0.67 | 0.43 |
| Hippocampus | 281 | 0.43 | 0.79 | 281 | 0.56 | -0.58 |
| Putamen | 282 | 0.52 | 0.64 | 282 | 0.41 | 0.83 |
| Thalamus[1] | 280 | 0.13 | -1.54 | 280 | 0.40 | -0.85 |

P-values (P) and t-values (t) for the effects of ADHD severity, as measured by hyperactivity/impulsivity and inattention symptoms, are indicated. P-values in **bold** are significant at the uncorrected level ($P < 0.05$). [1]Thalamus volume was not available from the NIH dataset

**Table S25**. Associations of cortical surface area AIs with disorder severity in ADHD individuals, all age groups combined.

| Region | Hyperactivity/impulsivity | | | Inattention | | |
|---|---|---|---|---|---|---|
| | N subjects | p-value | t-value | N subjects | p-value | t-value |
| banks of superior temporal sulcus | 286 | 0.97 | -0.04 | 286 | 0.35 | 0.93 |
| caudal anterior cingulate cortex | 322 | 0.16 | 1.42 | 322 | 0.90 | 0.12 |
| caudal middle frontal cortex | 322 | 0.30 | -1.04 | 322 | 0.55 | -0.60 |
| cuneus | 322 | 0.11 | 1.60 | 322 | 0.86 | -0.17 |
| entorhinal cortex | 320 | **0.03** | 2.12 | 320 | 0.07 | 1.85 |
| frontal pole | 322 | 0.69 | -0.40 | 322 | 0.92 | -0.10 |
| fusiform gyrus | 322 | 0.60 | 0.52 | 322 | 0.28 | 1.08 |
| inferior parietal cortex | 321 | 0.31 | -1.02 | 321 | 0.76 | -0.31 |
| inferior temporal gyrus | 321 | 0.26 | -1.14 | 321 | 0.43 | 0.79 |
| insula | 322 | 0.90 | -0.12 | 322 | 0.16 | 1.41 |
| isthmus cingulate cortex | 321 | 0.69 | -0.39 | 321 | 0.09 | -1.72 |
| lateral occipital cortex | 321 | 0.65 | 0.46 | 321 | 0.72 | 0.36 |
| lateral orbitofrontal cortex | 322 | 0.18 | 1.34 | 322 | 0.17 | 1.38 |
| lingual gyrus | 322 | 0.44 | -0.77 | 322 | 0.84 | 0.20 |
| medial orbitofrontal cortex | 321 | 0.62 | 0.49 | 321 | 0.67 | 0.42 |
| middle temporal gyrus | 308 | 0.92 | 0.10 | 308 | 0.24 | -1.17 |
| paracentral lobule | 322 | 0.58 | -0.55 | 322 | 0.18 | 1.35 |
| parahippocampal gyrus | 322 | 0.65 | -0.46 | 322 | 0.21 | 1.27 |
| pars opercularis of inferior frontal gyrus | 322 | 0.47 | -0.72 | 322 | 0.16 | -1.39 |
| pars orbitalis of inferior frontal gyrus | 322 | 0.28 | 1.07 | 322 | 0.29 | 1.07 |
| pars triangularis of inferior frontal gyrus | 322 | 0.81 | -0.23 | 322 | 0.83 | -0.21 |
| pericalcarine cortex | 322 | 0.99 | 0.01 | 322 | 0.75 | -0.31 |
| postcentral gyrus | 321 | 0.77 | -0.29 | 321 | 0.33 | 0.97 |
| posterior cingulate cortex | 321 | 0.52 | -0.65 | 321 | 0.49 | -0.69 |
| precentral gyrus | 322 | 0.94 | -0.08 | 322 | 0.14 | 1.50 |
| precuneus | 321 | 0.55 | 0.60 | 321 | 0.38 | -0.88 |
| rostral anterior cingulate cortex | 321 | 0.33 | 0.98 | 321 | 0.55 | 0.60 |
| rostral middle frontal gyrus | 322 | 0.85 | -0.19 | 322 | 0.83 | 0.22 |
| superior frontal gyrus | 322 | 0.07 | -1.84 | 322 | 0.12 | 1.54 |
| superiorparietal | 322 | 0.62 | 0.49 | 322 | 0.12 | -1.56 |
| superior temporal gyrus | 291 | 0.18 | 1.34 | 291 | 0.51 | 0.66 |
| supramarginal gyrus | 314 | 0.41 | 0.82 | 314 | 0.81 | -0.25 |
| temporal pole | 321 | 0.61 | 0.50 | 321 | 0.38 | 0.87 |
| transverse temporal gyrus | 322 | 0.64 | -0.47 | 322 | 0.07 | -1.84 |
| total average surface area | 322 | 0.81 | -0.24 | 322 | 0.43 | 0.79 |

P-values (P) and t-values (t) for the effects of ADHD severity, as measured by hyperactivity/impulsivity and inattention symptoms, are indicated. P-values in **bold** are significant at the uncorrected level (P < 0.05).

**Table S26**. Associations of cortical thickness AIs with disorder severity in ADHD individuals, all age groups combined.

| Region | Hyperactivity/impulsivity | | | Inattention | | |
|---|---|---|---|---|---|---|
| | N subjects | p-value | t-value | N subjects | p-value | t-value |
| banks of superior temporal sulcus | 286 | 0.78 | -0.27 | 286 | 0.80 | 0.25 |
| caudal anterior cingulate cortex | 322 | **0.01** | 2.66 | 322 | 0.31 | 1.03 |
| caudal middle frontal cortex | 321 | 0.08 | 1.74 | 321 | 0.80 | -0.25 |
| cuneus | 322 | 0.74 | 0.33 | 322 | 0.71 | 0.38 |
| entorhinal cortex | 321 | 0.68 | -0.41 | 321 | 0.62 | 0.49 |
| frontal pole | 321 | 0.37 | 0.89 | 321 | 0.79 | 0.26 |
| fusiform gyrus | 322 | 0.45 | -0.76 | 322 | 0.85 | 0.19 |
| inferior parietal cortex | 321 | 0.82 | -0.22 | 321 | 0.30 | -1.04 |
| inferior temporal gyrus | 321 | 0.79 | 0.26 | 321 | 0.39 | 0.85 |
| insula | 322 | 0.76 | -0.30 | 322 | 0.08 | -1.74 |
| isthmus cingulate cortex | 321 | 0.99 | 0.01 | 321 | 0.47 | -0.73 |
| lateral occipital cortex | 321 | 0.64 | 0.46 | 321 | 0.63 | -0.48 |
| lateral orbitofrontal cortex | 322 | 0.74 | 0.33 | 322 | 0.60 | -0.53 |
| lingual gyrus | 322 | 0.58 | -0.55 | 322 | 0.58 | -0.56 |
| medial orbitofrontal cortex | 322 | 0.11 | 1.61 | 322 | 0.94 | 0.07 |
| middle temporal gyrus | 308 | 0.83 | -0.21 | 308 | 0.81 | -0.24 |
| paracentral lobule | 322 | 0.11 | 1.60 | 322 | 0.74 | 0.33 |
| parahippocampal gyrus | 322 | 0.97 | -0.04 | 322 | 0.91 | -0.12 |
| pars opercularis of inferior frontal gyrus | 322 | **0.03** | 2.12 | 322 | **0.04** | 2.04 |
| pars orbitalis of inferior frontal gyrus | 322 | 0.08 | -1.75 | 322 | 0.44 | -0.78 |
| pars triangularis of inferior frontal gyrus | 320 | 0.31 | 1.02 | 320 | 0.08 | 1.74 |
| pericalcarine cortex | 322 | **0.02** | 2.41 | 322 | 0.31 | 1.02 |
| postcentral gyrus | 321 | 0.49 | -0.69 | 321 | 0.14 | 1.49 |
| posterior cingulate cortex | 320 | 0.27 | 1.11 | 320 | 0.28 | 1.09 |
| precentral gyrus | 322 | 0.48 | 0.70 | 322 | 0.39 | -0.87 |
| precuneus | 321 | 0.21 | 1.26 | 321 | 0.49 | 0.69 |
| rostral anterior cingulate cortex | 321 | 0.77 | -0.30 | 321 | 0.48 | 0.70 |
| rostral middle frontal gyrus | 322 | 0.35 | 0.94 | 322 | 0.76 | 0.30 |
| superior frontal gyrus | 322 | 0.14 | 1.48 | 322 | 0.59 | -0.54 |
| superiorparietal | 322 | 0.87 | -0.16 | 322 | 0.45 | -0.76 |
| superior temporal gyrus | 291 | 0.78 | -0.28 | 291 | 0.35 | 0.93 |
| supramarginal gyrus | 314 | 0.46 | -0.75 | 314 | 0.81 | 0.25 |
| temporal pole | 321 | 0.28 | -1.08 | 321 | 0.98 | -0.03 |
| transverse temporal gyrus | 322 | 0.31 | 1.01 | 322 | 0.17 | 1.37 |
| total average thickness | 322 | 0.11 | 1.59 | 322 | 0.49 | 0.70 |

P-values (P) and t-values (t) for the effects of ADHD severity, as measured by hyperactivity/impulsivity and inattention symptoms, are indicated. P-values in **bold** are significant at the uncorrected level ($P < 0.05$).

**Table S27**. Associations of subcortical volume AIs with psychostimulant medication use in ADHD individuals, all age groups combined.

| Region | Lifetime medication use | | | | Current medication use | | | |
|---|---|---|---|---|---|---|---|---|
| | N subjects (no/yes) | N datasets | p-value | t-value | N subjects (no/yes) | N datasets | p-value | t-value |
| Accumbens | 183/298 | 8 | 0.38 | -0.88 | 320/318 | 13 | 0.95 | 0.06 |
| Amygdala | 182/298 | 8 | 0.51 | 0.66 | 319/318 | 13 | 0.53 | 0.62 |
| Caudate Nucleus | 183/297 | 8 | 0.90 | -0.13 | 320/318 | 13 | 0.79 | 0.26 |
| Globus Pallidus | 183/298 | 8 | 0.46 | -0.74 | 320/318 | 13 | 0.09 | 1.70 |
| Hippocampus | 182/297 | 8 | 0.89 | 0.14 | 320/317 | 13 | 0.71 | 0.37 |
| Putamen | 183/297 | 8 | 0.29 | -1.06 | 320/318 | 13 | 0.07 | 1.78 |
| Thalamus[1] | 182/295 | 8 | 0.34 | 0.96 | 290/235 | 12 | 0.98 | -0.02 |

P-values (P) and t-values (t) for the effects of current and lifetime psychostimulant medication use are indicated. P-values in **bold** are significant at the uncorrected level (P < 0.05). [1]Thalamus volume was not available from the NIH dataset

**Table S28**. Associations of cortical surface area AIs with psychostimulant medication use in ADHD individuals, all age groups combined.

| Region | Lifetime medication use | | | | Current medication use | | | |
|---|---|---|---|---|---|---|---|---|
| | N subjects (no/yes) | N datasets | p-value | t-value | N subjects (no/yes) | N datasets | p-value | t-value |
| banks of superior temporal sulcus | 177/310 | 9 | 0.15 | 1.45 | 349/337 | 15 | 0.31 | -1.02 |
| caudal anterior cingulate cortex | 188/335 | 9 | 0.13 | 1.50 | 377/359 | 15 | 0.29 | 1.05 |
| caudal middle frontal cortex | 188/337 | 9 | 0.84 | 0.21 | 377/361 | 15 | 0.17 | -1.38 |
| cuneus | 186/337 | 9 | 0.60 | 0.52 | 374/358 | 15 | 0.75 | 0.32 |
| entorhinal cortex | 176/335 | 9 | 0.48 | -0.70 | 367/337 | 15 | 0.74 | -0.33 |
| frontal pole | 188/337 | 9 | 0.67 | 0.42 | 377/361 | 15 | 0.87 | 0.16 |
| fusiform gyrus | 187/335 | 9 | 0.98 | -0.03 | 376/359 | 15 | 0.96 | -0.04 |
| inferior parietal cortex | 186/335 | 9 | 0.98 | -0.02 | 375/361 | 15 | 0.94 | -0.08 |
| inferior temporal gyrus | 187/301 | 9 | 0.84 | 0.21 | 361/339 | 15 | 0.07 | 1.79 |
| insula | 179/336 | 9 | **0.04** | -2.03 | 377/360 | 15 | 0.07 | -1.83 |
| isthmus cingulate cortex | 188/337 | 9 | 0.27 | 1.11 | 377/360 | 15 | 0.28 | -1.09 |
| lateral occipital cortex | 186/337 | 9 | 0.61 | -0.51 | 377/361 | 15 | 0.98 | -0.02 |
| lateral orbitofrontal cortex | 188/337 | 9 | 0.81 | -0.23 | 376/361 | 15 | 0.61 | -0.52 |
| lingual gyrus | 187/337 | 9 | 0.75 | 0.32 | 377/361 | 15 | 0.24 | -1.17 |
| medial orbitofrontal cortex | 187/337 | 9 | 0.42 | 0.82 | 374/357 | 15 | 0.08 | 1.73 |
| middle temporal gyrus | 183/285 | 9 | 0.60 | 0.53 | 344/327 | 15 | 0.10 | 1.67 |
| paracentral lobule | 188/336 | 9 | 0.49 | -0.69 | 377/361 | 15 | 0.58 | 0.55 |
| parahippocampal gyrus | 188/337 | 9 | 0.28 | 1.08 | 377/359 | 15 | 0.36 | 0.91 |
| pars opercularis of inferior frontal gyrus | 188/334 | 9 | 0.32 | 1.00 | 376/356 | 15 | 1.00 | 0.00 |
| pars orbitalis of inferior frontal gyrus | 187/337 | 9 | 0.54 | 0.61 | 377/361 | 15 | 0.06 | 1.87 |
| pars triangularis of inferior frontal gyrus | 188/337 | 9 | 0.10 | 1.66 | 377/358 | 15 | 0.94 | -0.07 |
| pericalcarine cortex | 187/336 | 9 | 1.00 | 0.00 | 376/359 | 15 | 0.95 | -0.06 |
| postcentral gyrus | 180/337 | 9 | 0.29 | -1.05 | 370/354 | 15 | 0.99 | -0.02 |
| posterior cingulate cortex | 188/337 | 9 | 0.87 | -0.17 | 377/360 | 15 | 0.76 | -0.30 |
| precentral gyrus | 183/335 | 9 | 0.34 | 0.95 | 373/351 | 15 | 0.28 | 1.07 |
| precuneus | 187/337 | 9 | 0.97 | -0.03 | 377/361 | 15 | **0.02** | -2.25 |
| rostral anterior cingulate cortex | 187/334 | 9 | **0.05** | 1.97 | 372/357 | 15 | 0.26 | 1.13 |
| rostral middle frontal gyrus | 188/337 | 9 | 0.83 | -0.21 | 377/360 | 15 | 0.08 | -1.73 |
| superior frontal gyrus | 188/335 | 9 | 0.62 | 0.50 | 376/358 | 15 | 0.72 | 0.35 |
| superiorparietal | 185/336 | 9 | 0.28 | 1.08 | 375/360 | 15 | 0.19 | -1.32 |
| superior temporal gyrus | 179/276 | 9 | 0.62 | 0.49 | 336/317 | 15 | 0.10 | 1.65 |
| supramarginal gyrus | 184/333 | 9 | **0.04** | -2.08 | 374/353 | 15 | 0.51 | -0.67 |
| temporal pole | 188/337 | 9 | 0.70 | -0.39 | 377/360 | 15 | 0.40 | -0.84 |
| transverse temporal gyrus | 188/337 | 9 | 0.18 | -1.34 | 377/361 | 15 | **0.02** | -2.34 |
| total average surface area | 188/337 | 9 | 0.92 | 0.10 | 377/361 | 15 | 0.14 | -1.48 |

P-values (P) and t-values (t) for the effects of current and lifetime psychostimulant medication use are indicated. P-values in **bold** are significant at the uncorrected level ($P < 0.05$).

**Table S29**. Associations of cortical thickness AIs with psychostimulant medication use in ADHD individuals, all age groups combined.

| Region | Lifetime medication use | | | | Current medication use | | | |
|---|---|---|---|---|---|---|---|---|
| | N subjects (no/yes) | N datasets | p-value | t-value | N subjects (no/yes) | N datasets | p-value | t-value |
| banks of superior temporal sulcus | 177/310 | 9 | 0.26 | -1.13 | 349/337 | 15 | 0.48 | 0.71 |
| caudal anterior cingulate cortex | 188/335 | 9 | 0.89 | -0.13 | 377/359 | 15 | 0.94 | -0.08 |
| caudal middle frontal cortex | 188/337 | 9 | 0.24 | -1.18 | 376/361 | 15 | 0.81 | 0.24 |
| cuneus | 187/337 | 9 | 0.87 | 0.16 | 375/358 | 15 | 0.77 | 0.29 |
| entorhinal cortex | 176/335 | 9 | 0.95 | 0.06 | 367/337 | 15 | 0.50 | -0.67 |
| frontal pole | 188/337 | 9 | 0.80 | 0.25 | 377/361 | 15 | 0.62 | 0.50 |
| fusiform gyrus | 188/336 | 9 | 0.63 | 0.49 | 377/360 | 15 | 0.26 | -1.13 |
| inferior parietal cortex | 186/335 | 9 | 0.38 | 0.88 | 375/361 | 15 | **0.02** | -2.33 |
| inferior temporal gyrus | 187/301 | 9 | 0.70 | -0.39 | 361/339 | 15 | 0.40 | -0.85 |
| insula | 179/336 | 9 | 0.72 | 0.35 | 377/360 | 15 | 0.84 | 0.20 |
| isthmus cingulate cortex | 188/337 | 9 | 1.00 | 0.00 | 377/360 | 15 | 0.14 | 1.48 |
| lateral occipital cortex | 186/337 | 9 | 0.42 | 0.80 | 377/361 | 15 | 0.32 | 1.00 |
| lateral orbitofrontal cortex | 188/337 | 9 | 0.28 | 1.08 | 376/361 | 15 | 0.94 | 0.07 |
| lingual gyrus | 187/337 | 9 | 0.93 | -0.09 | 377/361 | 15 | 0.62 | 0.50 |
| medial orbitofrontal cortex | 187/337 | 9 | 0.28 | -1.08 | 374/357 | 15 | 0.42 | 0.81 |
| middle temporal gyrus | 184/285 | 9 | 0.49 | 0.69 | 345/327 | 15 | 0.48 | 0.70 |
| paracentral lobule | 188/336 | 9 | **0.03** | 2.16 | 377/361 | 15 | 0.55 | 0.60 |
| parahippocampal gyrus | 188/336 | 9 | 0.38 | 0.87 | 377/358 | 15 | 0.73 | -0.35 |
| pars opercularis of inferior frontal gyrus | 188/334 | 9 | 0.25 | 1.15 | 376/356 | 15 | 0.38 | 0.87 |
| pars orbitalis of inferior frontal gyrus | 187/337 | 9 | 0.75 | -0.31 | 377/361 | 15 | 0.64 | -0.47 |
| pars triangularis of inferior frontal gyrus | 188/337 | 9 | 0.94 | -0.08 | 375/358 | 15 | 0.94 | -0.08 |
| pericalcarine cortex | 187/336 | 9 | 0.87 | 0.17 | 376/359 | 15 | 0.09 | 1.72 |
| postcentral gyrus | 181/336 | 9 | 0.59 | -0.54 | 371/353 | 15 | 0.99 | 0.01 |
| posterior cingulate cortex | 188/337 | 9 | 0.46 | -0.74 | 376/360 | 15 | 0.87 | 0.17 |
| precentral gyrus | 183/335 | 9 | 0.26 | -1.13 | 373/351 | 15 | **0.03** | -2.16 |
| precuneus | 187/337 | 9 | 0.47 | 0.73 | 377/361 | 15 | 0.61 | 0.52 |
| rostral anterior cingulate cortex | 187/334 | 9 | 0.66 | -0.45 | 372/357 | 15 | 0.51 | -0.66 |
| rostral middle frontal gyrus | 188/337 | 9 | 0.84 | -0.20 | 377/360 | 15 | 0.38 | 0.88 |
| superior frontal gyrus | 188/335 | 9 | 0.70 | -0.38 | 376/358 | 15 | 0.91 | -0.11 |
| superiorparietal | 185/336 | 9 | 0.76 | -0.30 | 375/360 | 15 | 0.74 | 0.33 |
| superior temporal gyrus | 179/276 | 9 | 0.29 | -1.07 | 336/317 | 15 | 0.17 | 1.37 |
| supramarginal gyrus | 184/333 | 9 | 0.79 | 0.27 | 374/353 | 15 | 0.50 | -0.67 |
| temporal pole | 188/337 | 9 | 0.86 | 0.18 | 377/360 | 15 | 0.90 | -0.12 |
| transverse temporal gyrus | 188/337 | 9 | 0.21 | 1.26 | 377/361 | 15 | 0.80 | 0.26 |
| total average thickness | 188/337 | 9 | 0.68 | 0.42 | 377/361 | 15 | 0.80 | 0.25 |

P-values (P) and t-values (t) for the effects of current and lifetime psychostimulant medication use are indicated. P-values in **bold** are significant at the uncorrected level ($P < 0.05$).

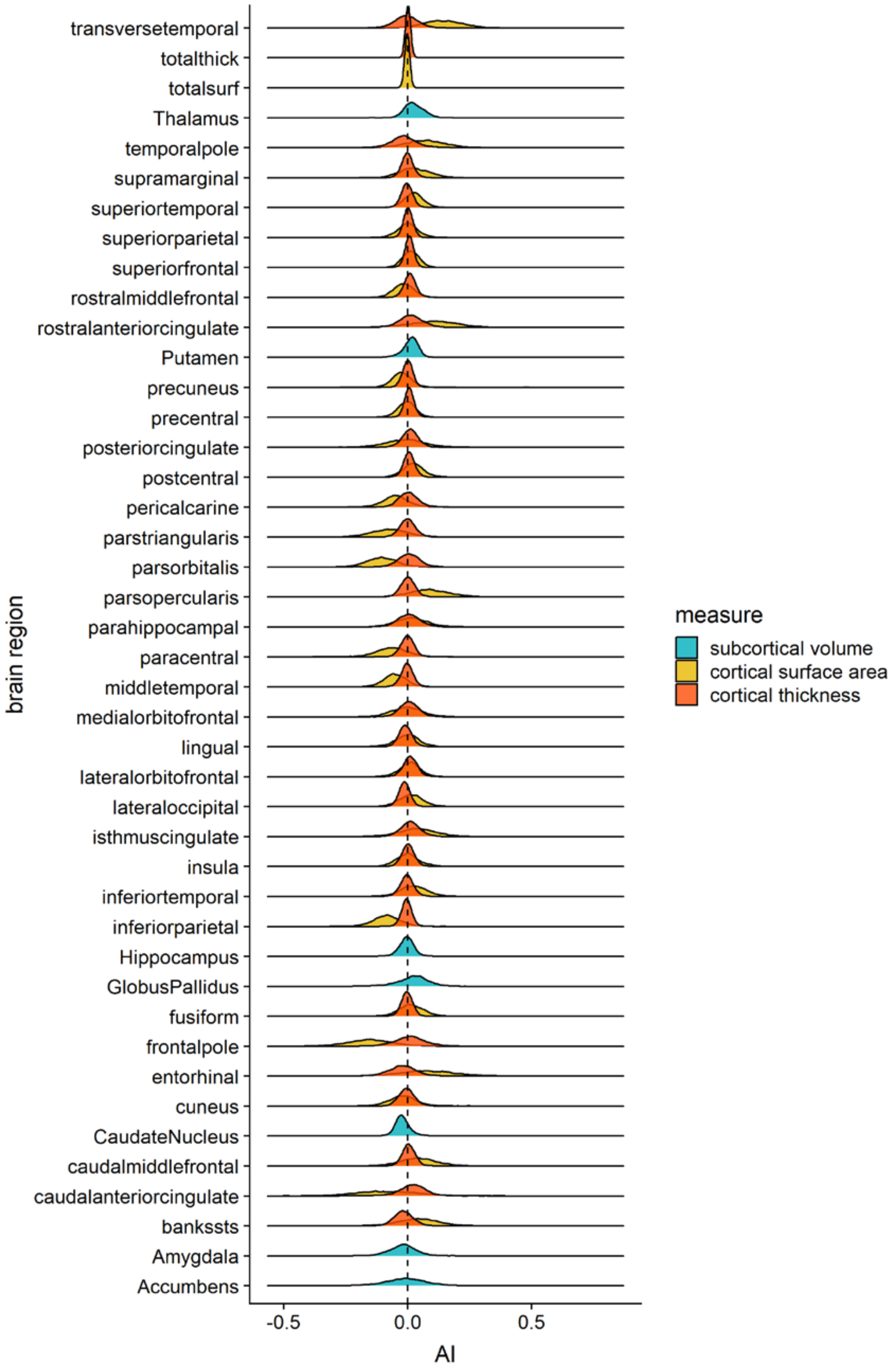

**Figure S1**. Joyplot of the distributions of AIs in the total study sample (without winsorization). Shown for subcortical volumes (*cyan*), cortical surface areas (*orange*), and cortical thicknesses (*red*).

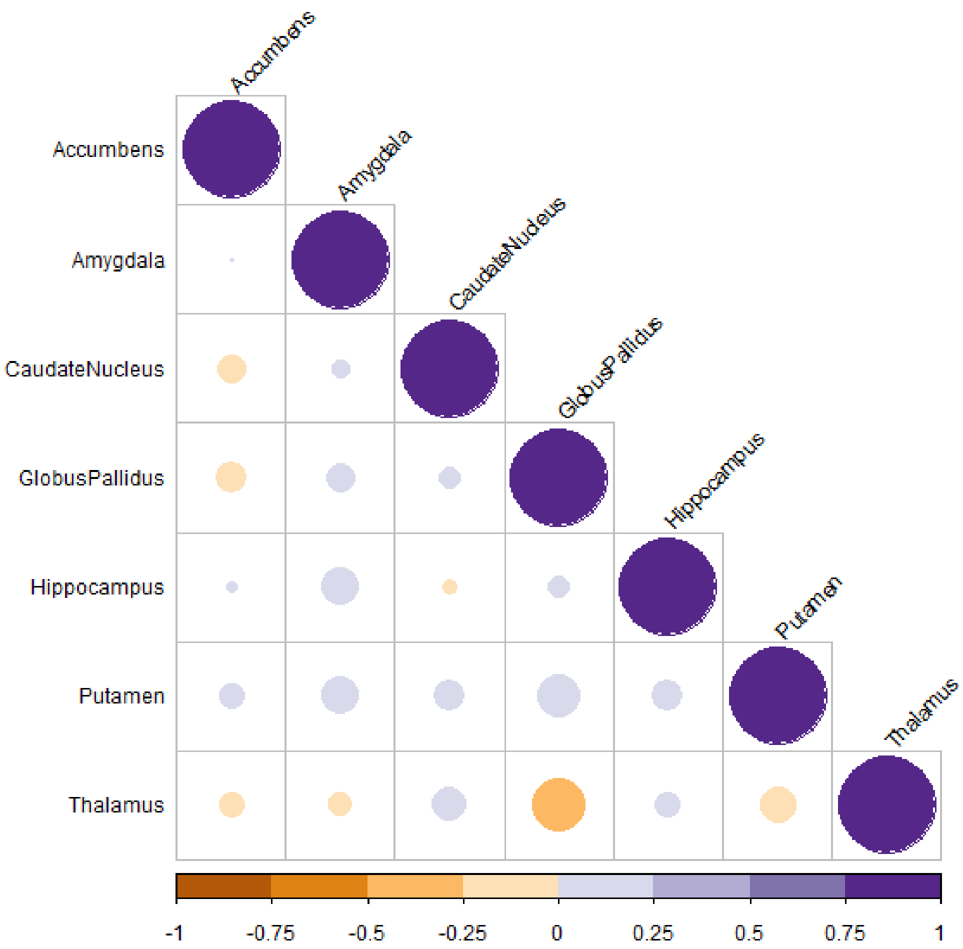

**Figure S2**. Correlations between AIs of subcortical volumes in the total study sample. Correlations ranged from -0.30 (between globus pallidus and thalamus) to 0.20 (between globus pallidus and putamen). Negative correlations are in orange and positive correlations are in purple. Color intensities and circle sizes are proportional to the magnitudes of the correlation coefficients.

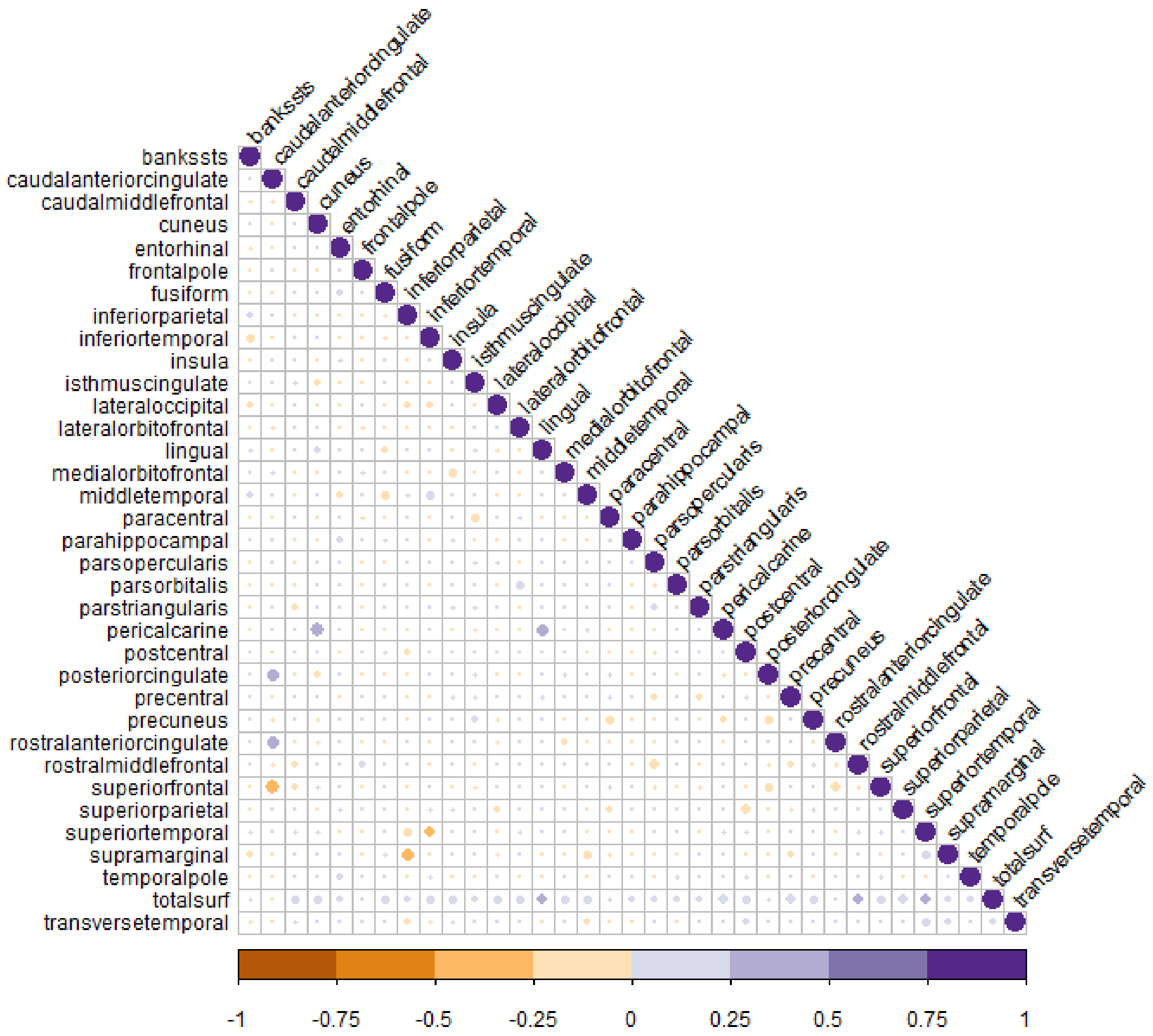

**Figure S3**. Correlations between AIs of cortical surface areas in the total study sample. Correlations ranged from -0.42 (between caudal anterior cingulate cortex and superior frontal gyrus) to 0.46 (between cuneus and pericalcarine cortex). Negative correlations are in orange and positive correlations are in purple. Color intensities and circle sizes are proportional to the magnitudes of the correlation coefficients.

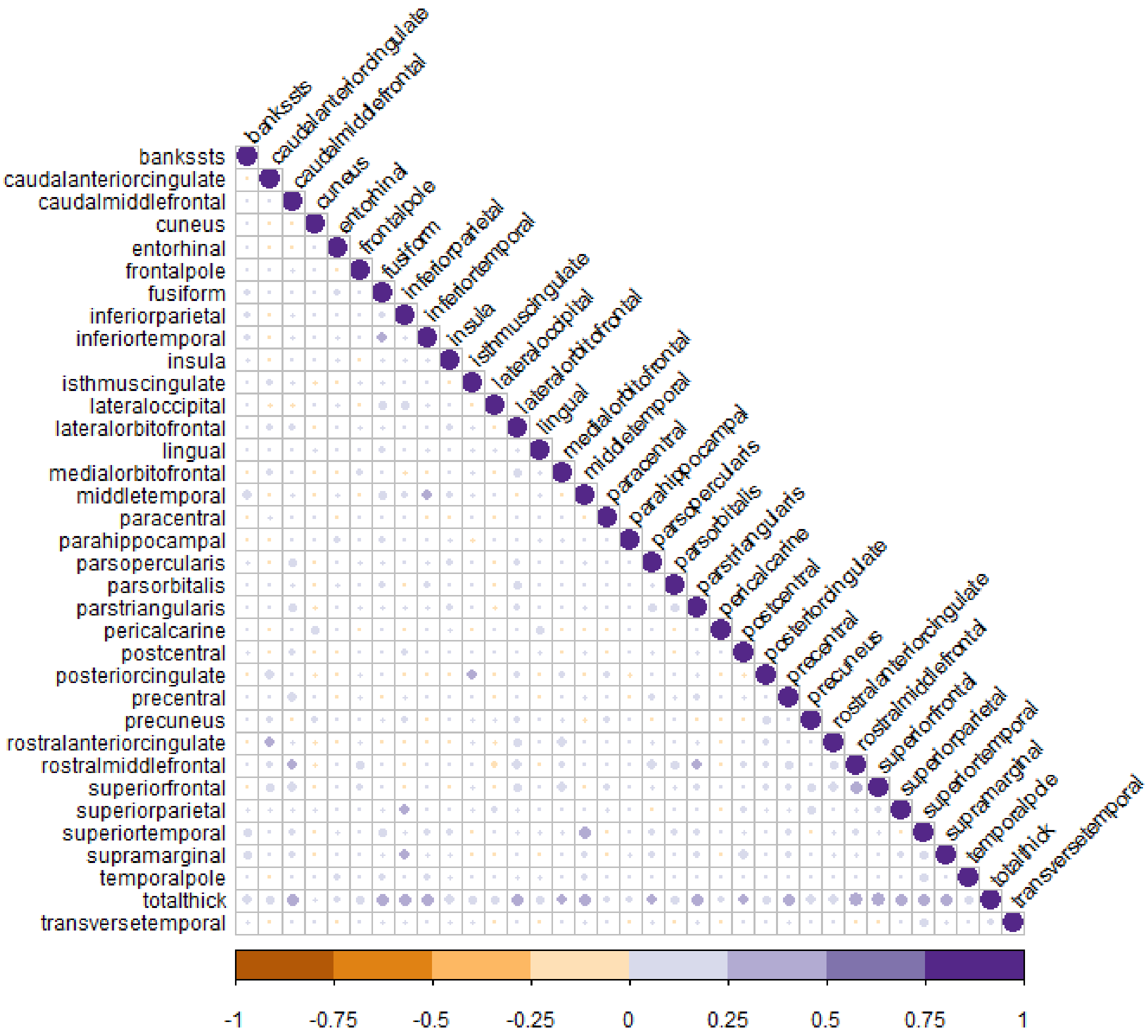

**Figure S4**. Correlations between AIs of cortical thicknes in the total study sample. Correlations ranged from -0.11 (between lateral occipital cortex and rostral middle frontal cortex) to 0.49 (between rostral middle frontal cortex and total average thickness). Negative correlations are in orange and positive correlations are in purple. Color intensities and circle sizes are proportional to the magnitudes of the correlation coefficients.

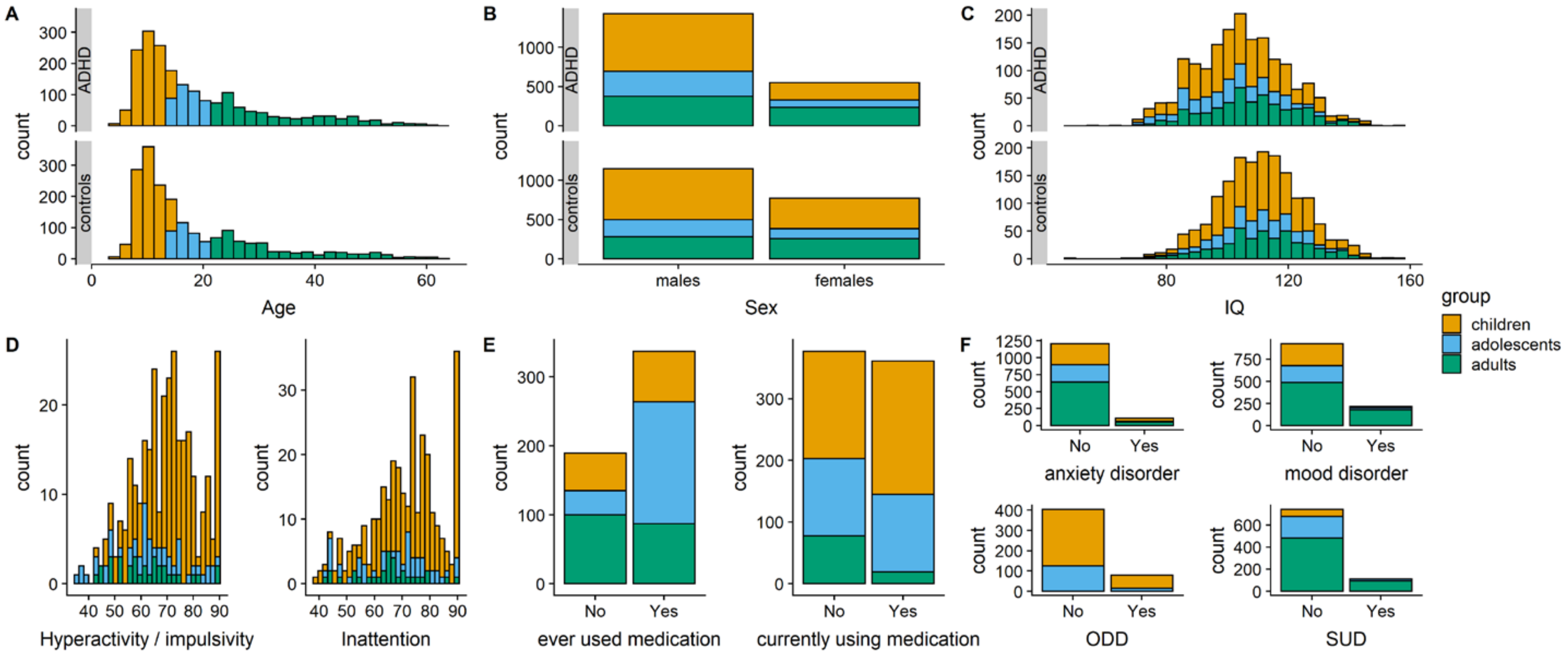

**Figure S5**. Distributions of (**A**) age, (**B**) sex and (**C**) IQ in ADHD and controls, and of (**D**) ADHD severity, (**E**) psychostimulant medication use, and (**F**) comorbidity in ADHD-only, colored by children (*orange*), adolescents (*blue*) and adults (*green*).

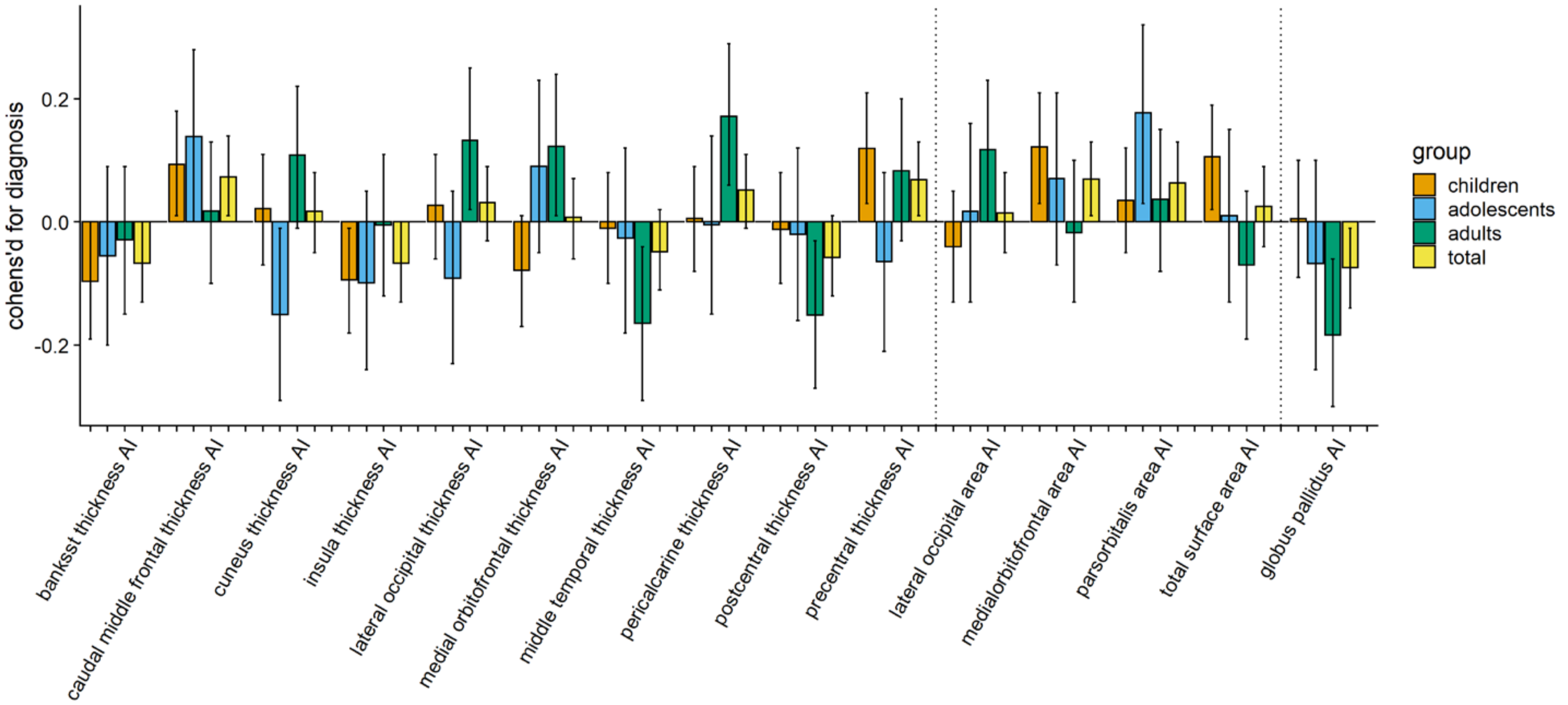

**Figure S6.** Bar plots of the Cohen's *d* effect sizes for diagnosis in the different age groups analyzed. Shown are only those AIsthat showed a nominally significant effect of diagnosis in any of the analyses. All Cohen's *d* values above zero represent a mean shift towards greater leftward or reduced rightward asymmetry in ADHD compared to controls, and those below zero represent mean shifts towards greater rightward of reduced leftward asymmetry in ADHD compared to controls. The different age groups are shown in different colors: *orange* = children; *blue* = adolescents; *green* = adults; *yellow* = all age groups combined. The solid vertical lines reflect the error bars, indicating the 95% CI interval around Cohen's *d*, and the dotted vertical lines separate the different types of measure (i.e., thickness AIs, surface area AIs, subcortical volume AIs).

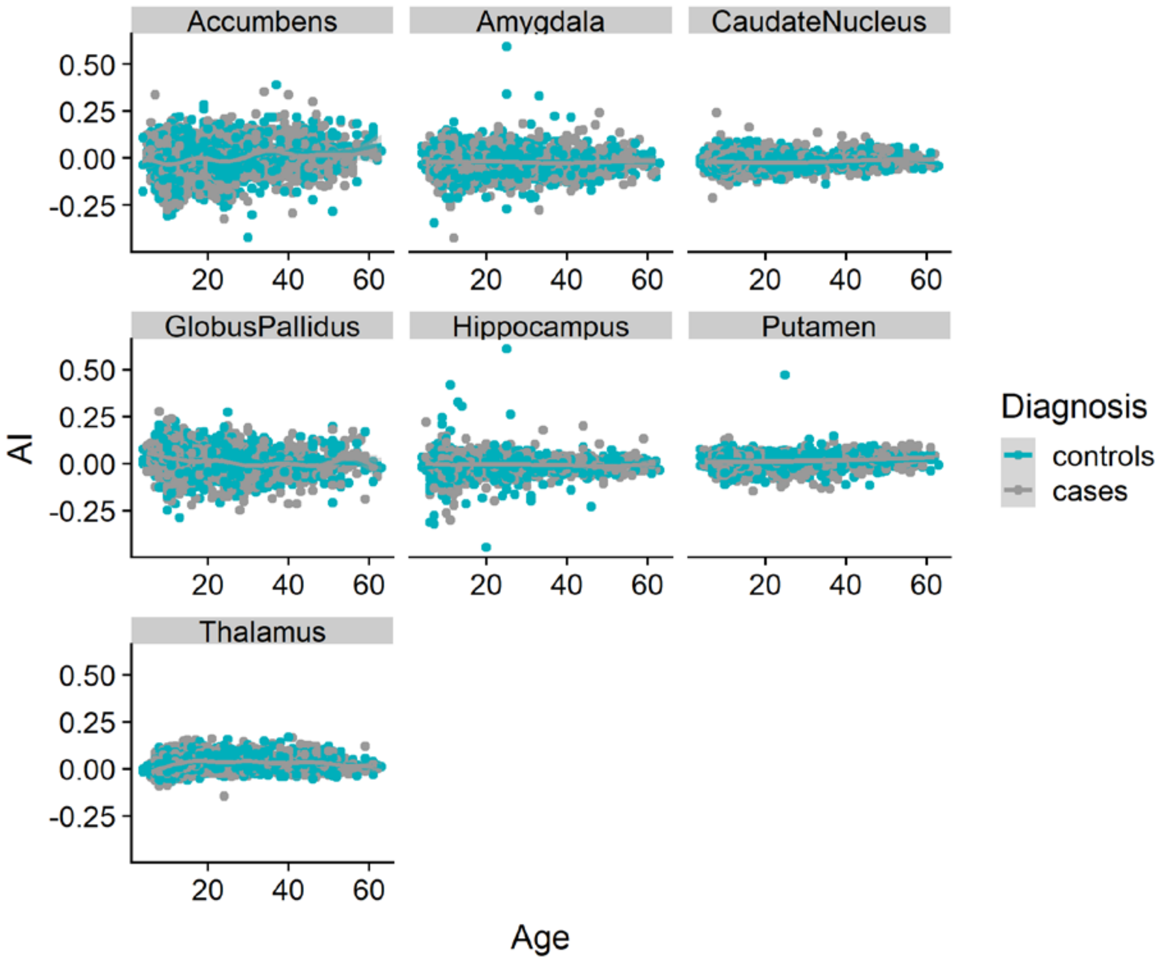

**Figure S2**. Scatter plots of the relationship between age and AIs of the subcortical volumes.

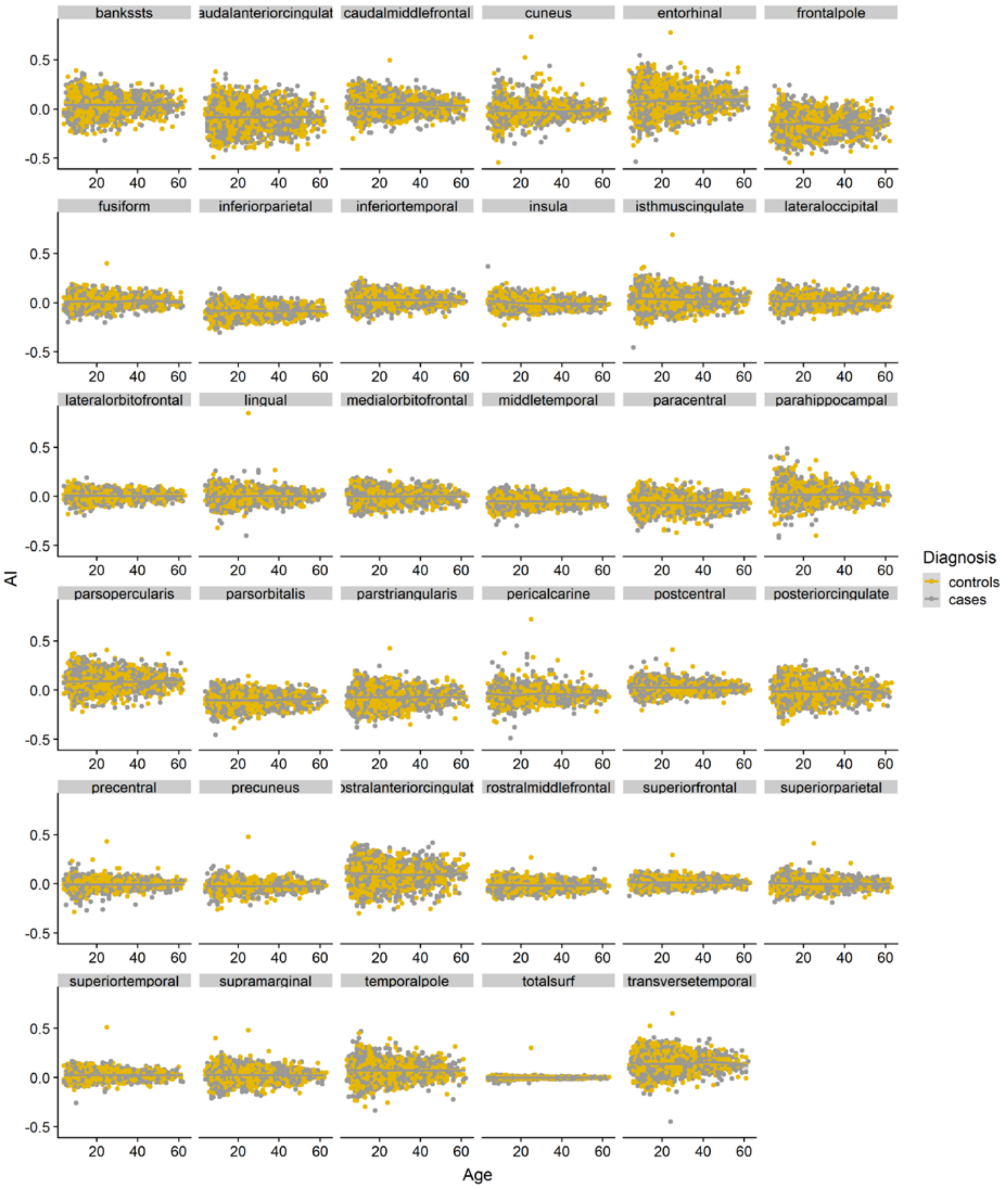

**Figure S3**. Scatter plots of the relationship between age and AIs of the cortical surface areas.

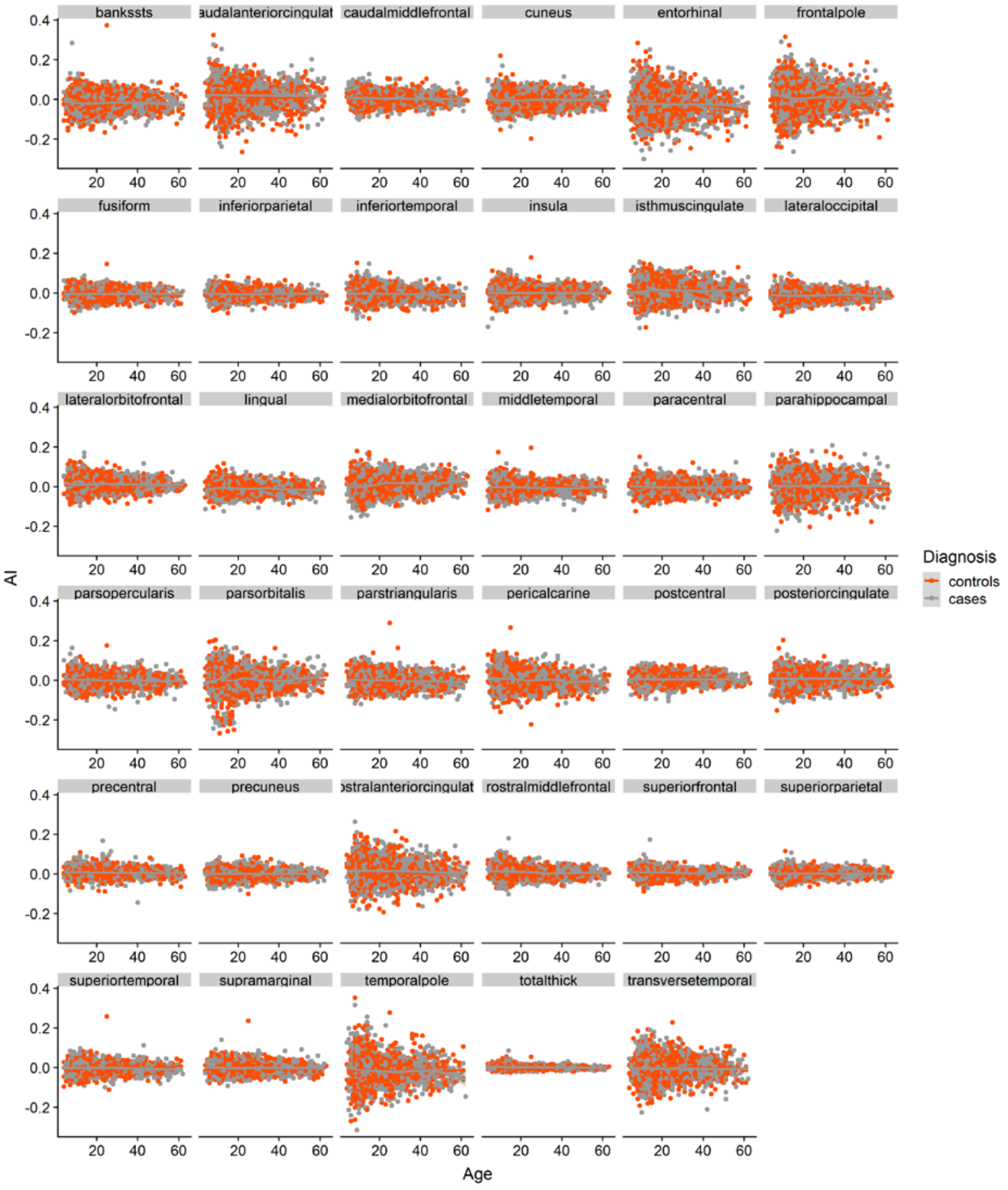

**Figure S4**. Scatter plots of the relationship between age and AIs of the cortical thickness.

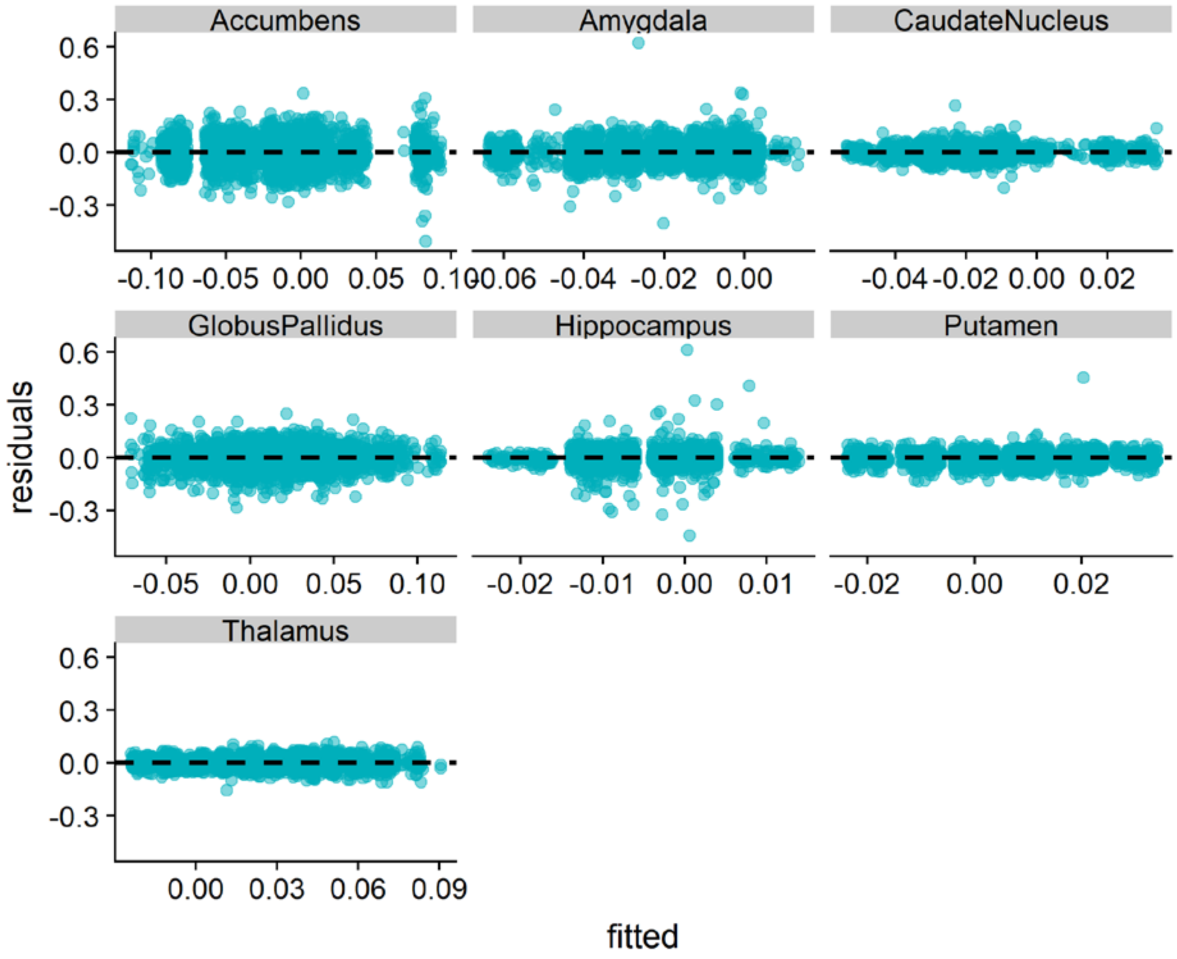

**Figure S5**. Residual plots of the linear mixed effects model analysis of subcortical volume AIs in the total study sample. The ggplot2 package in R was used to visualize residuals.

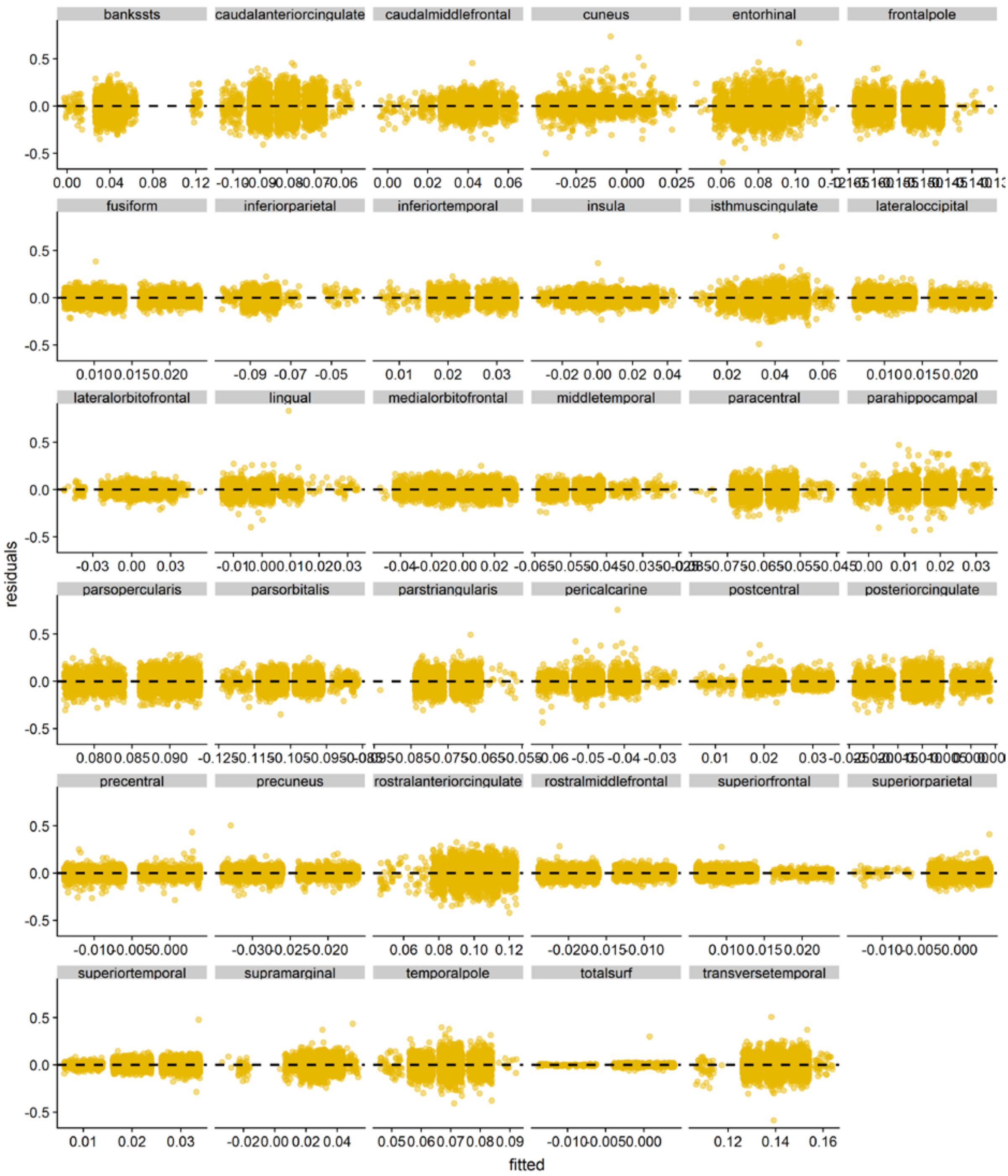

**Figure S6**. Residual plots of the linear mixed effects model analysis of cortical surface area AIs and the AI of the total average surface area (totalsurf) in the total study sample. The ggplot2 package in R was used to visualize residuals.

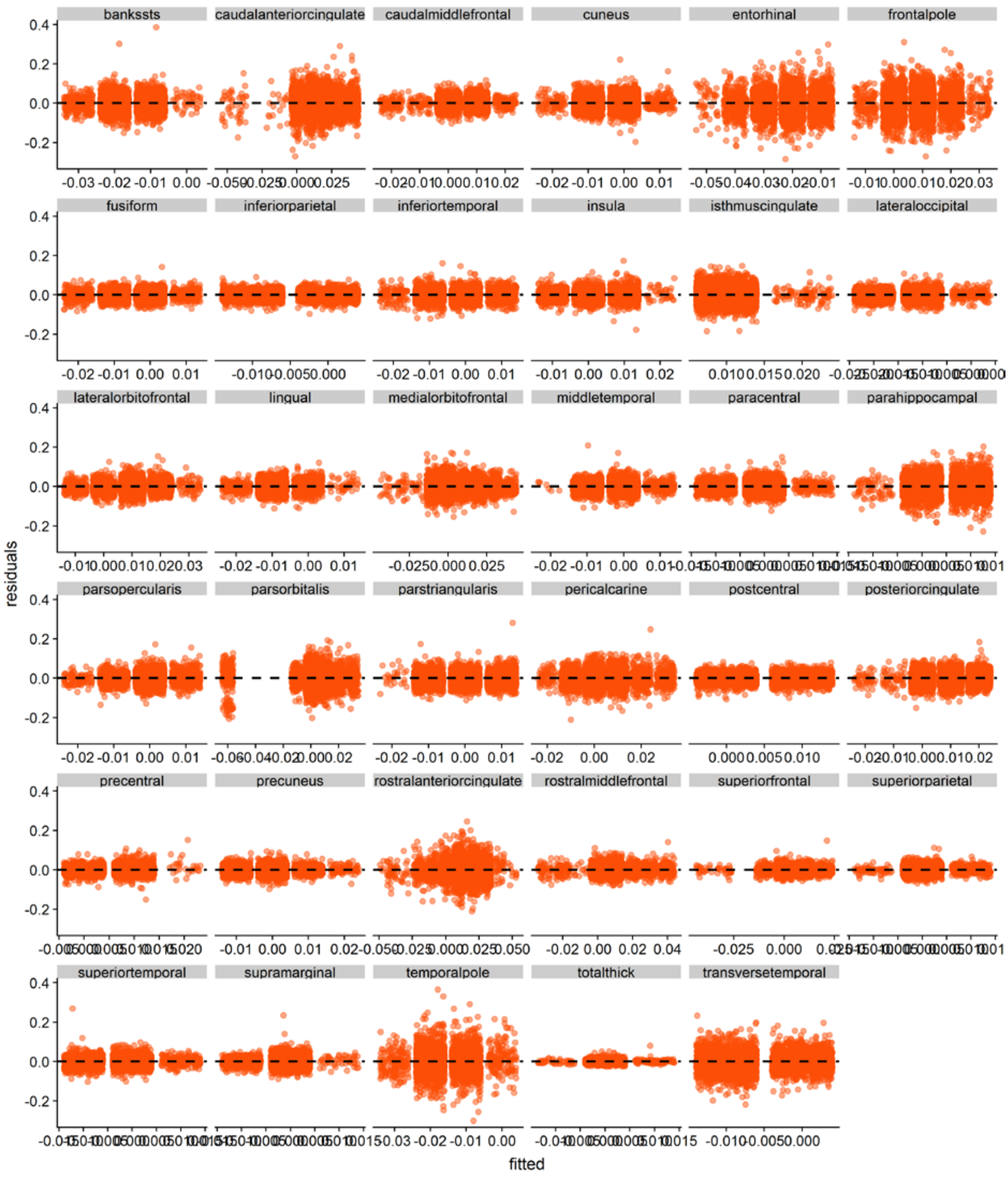

**Figure S7**. Residual plots of the linear mixed effects model analysis of cortical thickness AIs and the AI of the total average thickness (totalthick) in the total study sample. The ggplot2 package in R was used to visualize residuals.

#### Supplementary References

1. Barrantes-Vidal N, Gómez-de-Regil L, Navarro B, Vicens-Vilanova J, Obiols J, Kwapil T. Psychotic-like symptoms and positive schizotypy are associated with mixed and ambiguous handedness in an adolescent community sample. Psychiatry Research. (0).
2. Spitzer RL, Williams JB, Gibbon M, First MB. The Structured Clinical Interview for DSM-III-R (SCID). I: History, rationale, and description. Arch Gen Psychiatry. 1992;49(8):624-9.
3. Kaufman J, Birmaher B, Brent D, Rao U, Flynn C, Moreci P, et al. Schedule for Affective Disorders and Schizophrenia for School-Age Children-Present and Lifetime Version (K-SADS-PL): initial reliability and validity data. Journal of the American Academy of Child and Adolescent Psychiatry. 1997;36(7):980-8.
4. Conners CK. Rating scales in attention-deficit/hyperactivity disorder: use in assessment and treatment monitoring. J Clin Psychiatry. 1998;59 Suppl 7:24-30.
